# Microbiome Data Integration via Shared Dictionary Learning

**DOI:** 10.1101/2024.10.04.616752

**Authors:** Bo Yuan, Shulei Wang

## Abstract

Data integration is a powerful tool for facilitating a comprehensive and generalizable understanding of microbial communities and their association with outcomes of interest. However, integrating data sets from different studies remains a challenging problem because of severe batch effects, unobserved confounding variables, and high heterogeneity across data sets. We propose a new data integration method called MetaDICT, which initially estimates the batch effects by weighting methods in causal inference literature and then refines the estimation via novel shared dictionary learning. Compared with existing methods, MetaDICT can better avoid the overcorrection of batch effects and preserve biological variation when there exist unobserved confounding variables, data sets are highly heterogeneous across studies, or the batch is completely confounded with some covariates. Furthermore, MetaDICT can generate comparable embedding at both taxa and sample levels that can be used to unravel the hidden structure of the integrated data and improve the integrative analysis. Applications to synthetic and real microbiome data sets demonstrate the robustness and effectiveness of MetaDICT in integrative analysis. Using MetaDICT, we characterize microbial interaction, identify generalizable microbial signatures, and enhance the accuracy of outcome prediction in two real integrative studies, including an integrative analysis of colorectal cancer metagenomics studies and a meta-analysis of immunotherapy microbiome studies.

## 1 Introduction

Recent advances in metagenomic sequencing technologies make it possible to profile the microbiome communities from hundreds to thousands of samples in scientific studies (Turnbaugh et al., 2009; Yatsunenko et al., 2012; Franzosa et al., 2019). These microbiome studies provide glimpses into the complex microbial ecosystem and improve our understanding of the interactions between the microbes and their host. Although each study has already yielded interesting results, the findings from different studies are not always consistent and the power could be limited due to the small sample size of each study (Langdon et al., 2016; Duvallet et al., 2017). One promising strategy to obtain generalizable discoveries is the integrative analysis of the data sets from multiple studies (Wirbel et al., 2019; Ma et al., 2022). However, integrative microbiome data analysis presents unique quantitative challenges as the data from different studies are collected across times, locations, or sequencing protocols and thus suffer severe batch effects and are highly heterogeneous. When handled inappropriately, the batch effects and high heterogeneity could lead to increased false discoveries and reduced accuracy in the downstream integrative analysis (Ling et al., 2022).

In order to correct batch effect and facilitate a valid integrative analysis, one of the most popular strategies in integrative studies is to apply the regression models to correct the batch effects, where the sequencing count of each taxon is the outcome and covariates include batch and each sample’s observed covariates (Gibbons et al., 2018; Zhang et al., 2020; Ma et al., 2022; Ling et al., 2022; Ye et al., 2023). The primary assumption behind such a strategy is that the conditional distribution/expectation of sequencing count remains the same after successfully adjusting the effects of batch and observed covariates. The covariate adjustment methods can correct the batch effects efficiently when all confounding covariates are observed and adjusted appropriately. However, this assumption could be invalid, and overcorrection happens when there are some important unmeasured confounding covariates, such as lifestyle. Besides the covariate adjustment strategy, another popular strategy is to utilize the intrinsic structure of data to correct batch effects (Haghverdi et al., 2018; Butler et al., 2018; Hie et al., 2019; Korsunsky et al., 2019; Welch et al., 2019; Amodio et al., 2019; Barkas et al., 2019; Peng et al., 2021). The main advantage of such a strategy is that exploring intrinsic structure does not rely on extrinsic covariates and is thus robust to unmeasured confounding covariates. Most methods in this strategy are designed for single-cell RNA sequencing data since the covariate adjustment is not applicable in single-cell RNA sequencing data due to no access to cell-level covariates. However, this strategy does not utilize the valuable information in observed covariates when available, and the assumptions adopted in existing methods are not general enough to work for the microbiome data. For example, one commonly used assumption in these methods is that different cell types can be separated well into clusters, and cluster structure can be used as an anchor for the integration. However, this assumption is not always valid for microbiome data sets as microbiome data may be separated poorly into several groups when dependent on continuous confounding variables. The challenges above raise questions about whether it is possible to develop a new data integration method for microbiome data that combines the advantages of these two popular strategies. This paper shows that this is feasible.

This paper introduces MetaDICT, a new two-stage data integration method for microbiome data. With a similar spirit to existing covariate adjustment methods, the first stage of MetaDICT obtains an initial estimation of batch effects via adjusting commonly observed covariates. However, MetaDICT adopts the weighting method in causal inference literature to adjust covariates instead of the commonly used regression-based methods because several studies confirm that the batch effects affect the sequencing counts multiplicatively rather than additively (Harismendy et al., 2009; McLaren et al., 2019). In the second stage of MetaDICT, the estimation of batch effects is further refined via novel shared dictionary learning. Through shared dictionary learning, MetaDICT explores the intrinsic structures that are sufficiently flexible to capture the characteristics of the microbiome data and can thus disentangle the batch effect from the heterogeneous data sets robustly. Thanks to the shared dictionary learning, MetaDICT can better address the overcorrection of batch effects and preserve biological variation than existing methods in diverse settings, including ones where there are unmeasured confounding covariates, data sets are highly heterogeneous across studies, or the batch is completely confounded with some covariates. Beyond batch effect correction, shared dictionary learning in MetaDICT also generates the embedding at both taxa and sample levels that can be used to unravel the hidden structure of the integrated data and improve the integrative analysis. Comprehensive numerical experiments presented in this paper demonstrate the efficacy of MetaDICT in correcting batch effect, reducing false discoveries, and enhancing the power of integrative analysis. In particular, we apply MetaDICT to an integrative analysis of five colorectal cancer metagenomics studies where each study is conducted in a different country. In the integrative analysis, MetaDICT can successfully separate the batch effect and effect of country, reveal the microbial functional similarity, detect population structure, identify previously documented and novel microbial signatures of colorectal cancer, and improve the accuracy and generalizability of disease diagnosis. Furthermore, we also apply MetaDICT to a meta-analysis of five microbiome data sets of anti-programmed cell death protein-1 treated patients with melanoma. With the help of MetaDICT, the integrative analysis identifies consistent microbial signatures associated with clinical response and constructs an efficient predictive model for immunotherapy outcomes.

## 2 Results

### 2.1 Overview of MetaDICT

This section presents an overview of MetaDICT, while the Methods section comprehensively explains the proposed method. As summarized in Figure 1, the newly proposed MetaDICT consists of two stages: the first stage provides an initial estimation of batch effect via covariate balancing and the second stage refines the estimation by shared dictionary learning. MetaDICT defines the batch effect as heterogeneous capturing efficiency in sequencing measurement, which refers to the proportion of microbial DNA from a sample that is successfully extracted, amplified, incorporated into a sequencing library, and ultimately detected in the final sequencing data. The measurement efficiency is highly influenced by the variations in technical factors and external conditions of the sequencing process (Lander, 1999; Morgan et al., 2010) and usually affects observed sequencing count in a multiplicative way (Harismendy et al., 2009; McLaren et al., 2019). The first stage of MetaDICT estimates the measurement efficiency by the weighting method in causal inference (Imbens and Rubin, 2015), one of the most popular covariate balancing methods. The initial estimation from the weighting method is accurate when all confounding variables are observed in different studies. However, this strategy might result in overcorrection when we only observe a few common variables across studies or have difficulty in measuring some important confounding variables in practice, like lifestyle. MetaDICT further refines the estimation in the presence of unobserved confounding variables to increase the robustness.

**Figure 1:**
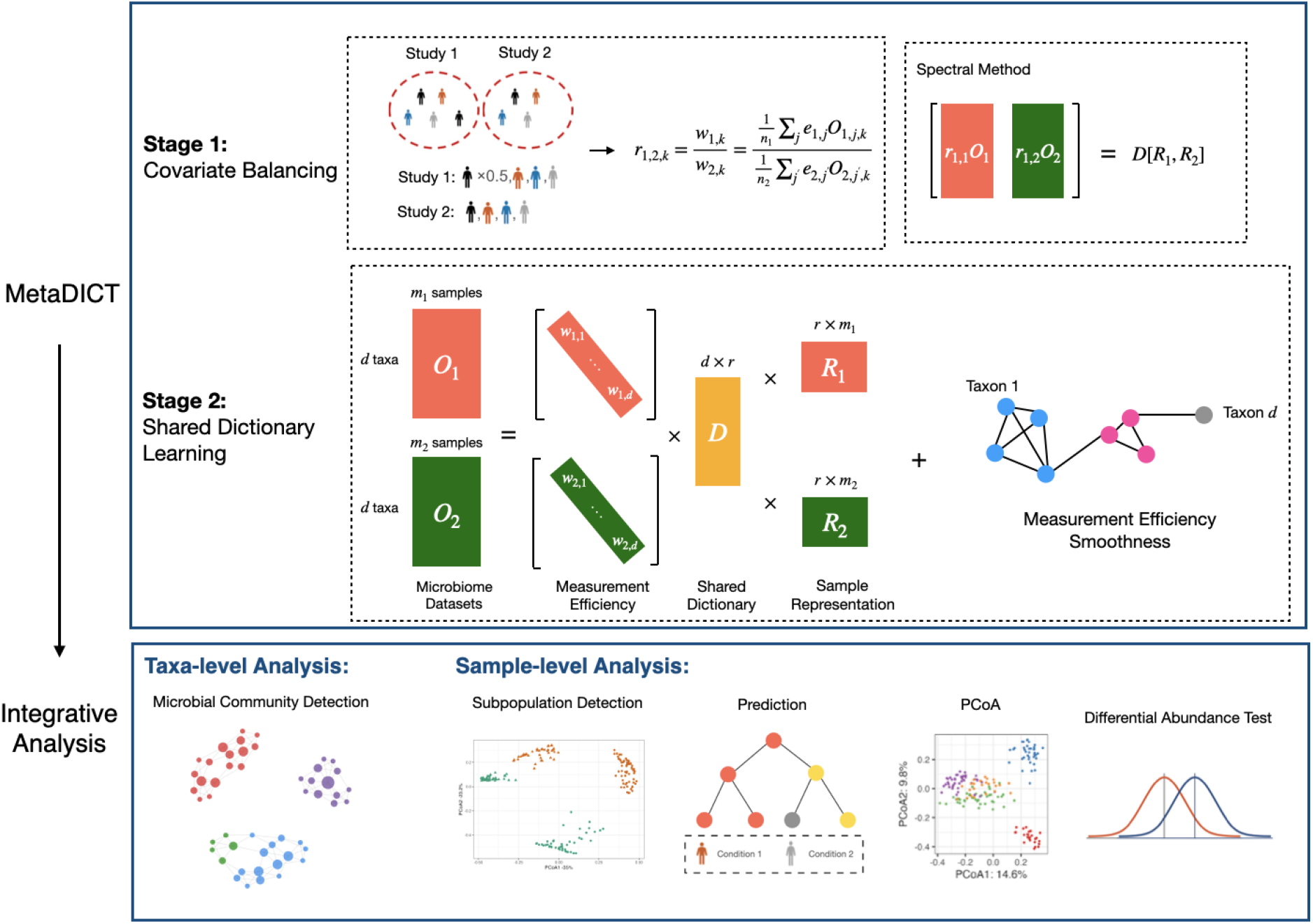
A summary of MetaDICT for integrative analysis. Stage 1: the weighting method in the causal inference literature adjusts the commonly observed covariates to estimate the batch effect initially. The weighting method balances the distribution of covariates across data sets by assigning appropriate weights to each sample. In the example, different covariates are indicated by different colors. If a certain covariate appears twice as often in group 1 compared to group 2, we assign a weight of 0.5 to individuals in group 1 with that covariate to ensure balance. Stage 2: the batch effect estimation is refined by exploring the intrinsic structures of microbiome data: the shared dictionary of microbial profile and measurement efficiency’s smoothness. In addition to batch effect correction, MetaDICT also generates embedding of taxa and samples for efficient integrative analysis.

Unlike covariate adjustment, the second stage of MetaDICT aims to improve the estimation of measurement efficiency via exploring two types of intrinsic structures in microbiome data: shared dictionary and the measurement efficiency’s smoothness. Despite the variation in sequencing procedures, the microbes interact and coexist as an ecosystem similarly in different studies (Woyke et al., 2006; Chaffron et al., 2010). Motivated by this observation, MetaDICT introduces a shared dictionary of microbial absolute abundance to capture such a universal structure across studies. As a popular representation learning method in computer vision and genomics (Elad and Aharon, 2006; Wright et al., 2008; Pan et al., 2022; Hao et al., 2024), dictionary learning aims to represent data as a linear combination of basic elements, which are called atoms and together form a dictionary. In the context of microbiome data integration, each atom represents a group of microbes of which abundance change is highly correlated, and the shared dictionary is the collection of such correlated patterns that are universal across studies. Reconstructing the microbial loads from the shared dictionary allows for identifying universal patterns and efficient data integration. Besides the shared dictionary, another important observation is that microbes with close taxonomic sequences tend to have similar capturing efficiency in each study (Krsek and Wellington, 1999; Polz and Cavanaugh, 1998; Carrigg et al., 2007; Benjamini and Speed, 2012). This observation indicates that measurement efficiency is smooth with respect to the similarity among taxa, allowing borrowing strength from similar taxa. To capture such smoothness, we adopt the graph Laplacian to measure the smoothness of measurement efficiency with respect to the graph constructed by a phylogenetic tree or taxonomic information. The second stage of MetaDICT solves a nonconvex optimization problem to explore the above two intrinsic structures of microbiome data, which is initialized by a spectral method and the estimation in the first stage. By utilizing intrinsic structures, MetaDICT can adjust the batch effect robustly, avoid overcorrection efficiently, and maintain the biological variation in the downstream integrative analysis. In addition to batch effect correction, the estimated shared dictionary in MetaDICT can naturally offer embeddings of taxa and samples, revealing the microbial communities and improving the performance of downstream analysis. In the following sections, we demonstrate the robustness and effectiveness of MetaDICT in correcting batch effect and improving the downstream integrative analysis.

### 2.2 MetaDICT Corrects Batch Effect Robustly with Unobserved Confounders

This section designs a series of numerical experiments to evaluate the MetaDICT’s performance of correcting batch effect. As summarized in Figure 1, one unique feature of the MetaDICT is to explore the intrinsic structure of microbiome data via shared dictionary learning. The first set of numerical experiments aims to assess whether the shared dictionary learning in the MetaDICT can improve the initial estimation in the first stage. To mimic the real data, we generate the synthetic data from a microbiome data set collected by He et al. (2018). We consider two criteria to assess the performance: the mean absolute error of estimated measurement efficiency at each taxon (Figure 2(a)) and the Pearson correlation coefficient between the estimated and true measurement efficiency (Figure 2(b)). These comparisons suggest that utilizing the intrinsic structure improves the accuracy of estimated measurement efficiency at both taxa and data set levels. Exploiting the smoothness of measurement efficiency is one of the major reasons why shared dictionary learning can increase accuracy. As illustrated in Figure S1(a), the penalty for measurement efficiency’s smoothness enables borrowing strength from similar taxa, thus improving the performance (Figure S1(b)). Furthermore, we also validate the robustness of MetaDICT in a wide range of settings, including experiments when the number of data sets, sample sizes per data set, the balance of data set sizes, and the measurement efficiency smoothness level are different (Figure S2). The results presented in Figure 2, S1, and S2 indicate that MetaDICT can recover measurement efficiencies accurately and robustly.

**Figure 2:**
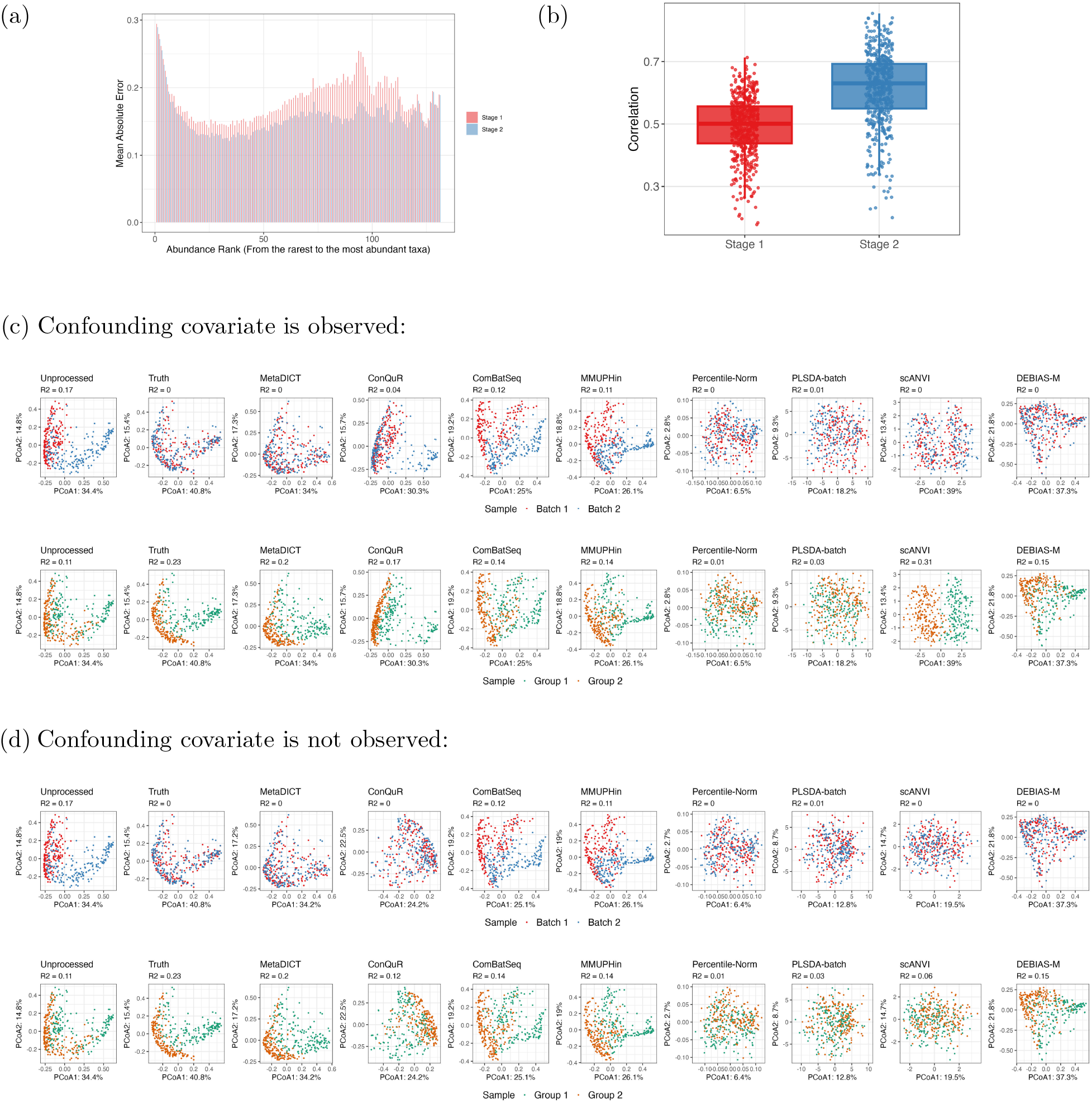
MetaDICT estimates measurement efficiency accurately and corrects batch effects robustly. Figures (a) and (b) compare the estimation accuracy of measurement efficiency between stages 1 and 2, showing that exploring intrinsic structure can significantly improve estimation accuracy. In (a), the mean absolute error of each taxon is compared, with taxa ordered from least to most abundant. In (b), the *y*-axis represents the Pearson correlation coefficients between the estimated measurement efficiency and the ground truth. These comparisons suggest that Stage 2 of MetaDICT can improve the accuracy of estimated measurement efficiency. Figures (c) and (d) compare the performance of eight data integration methods, demonstrating that MetaDICT corrects batch effects robustly and maintains biological variation effectively. In the experiment, we simulated two batches of data, and the synthetic samples were associated with a randomly generated binary biological variable (group 1 or 2). The unprocessed panels display the simulated data with batch effects, and the truth panels display the batch-effect-free data (i.e., the microbial absolute abundance). Figure (d) shows the PCoA plots and *R*^2^ in PERMANOVA when the biological variable is not observed in advance, while Figure (c) presents the results when the biological variable is used as input for all methods. The data integration methods aim to recover the truth panels as accurately as possible. Bray-Curtis dissimilarity on corrected abundances is used for all methods except scANVI and PLSDA-batch, as scANVI and PLSDA-batch provide a new representation rather than corrected counts. Euclidean distance is applied for these two methods instead.

The valid estimation of measurement efficiency leads to robust batch effect correction results, even when the initial estimation of measurement efficiency using sample covariates is inaccurate. The next sets of numerical experiments investigate whether MetaDICT can help correct batch effects and maintain the biological variations when sample covariate is unobserved. We compare MetaDICT with seven state-of-the-art data integration methods: ConQuR (Ling et al., 2022), ComBat-Seq (Zhang et al., 2020), MMUPHin (Ma et al., 2022), Percentile Normalization (Gibbons et al., 2018), PLSDA-batch (Wang and Lê Cao, 2023), DEBIAS-M (Austin et al., 2024), and an autoencoder-based method scANVI (Xu et al., 2021). The synthetic data include two simulated batches. The microbial absolute abundance relies on a binary biological variable, which can be considered as the indicator of health conditions. We adjust the abundance of a subset of taxa to introduce differences between the two biological conditions. Compared with microbial loads, the perturbation due to batch effects results in a distribution change across studies and less separated clusters between groups defined by the biological variable (Figure 2(c), (d), and S6). We consider two possible scenarios to correct the batch effect: 1) the confounding biological variable is observed and used in batch effect correction; 2) the confounding biological variable is not observed and cannot be used in batch effect correction. When the biological variable is observed, MetaDICT, ConQuR, DEBIAS-M and scANVI can adjust the effect of the biological variable well, thus successfully correcting the batch effect and maintaining a decent amount of biological variation (Figure 2(c)). On the other hand, if the biological variable is not observed, MetaDICT can still separate clusters defined by the biological variable, while ConQuR and scANVI, whose performance is highly sensitive to the observed confounding variable, wrongly reduce the biological variation (Figure 2(d)). Similar results are also observed when data sets are simulated with SparseDOSSA2 (Ma et al., 2021) (Figure S7). Besides the binary biological variable, we consider a similar experiment when the biological variable is continuous (Figure S3). The results suggest that exploring the intrinsic structure of microbiome data in MetaDICT can significantly increase the robustness of batch effect correction even when the important confounding covariate is not observed.

### 2.3 MetaDICT Prevents Overcorrection When Biological Heterogeneity Exists Across Batches

One of the major challenges in data integration is effectively removing batch effects without overcorrection when biological heterogeneity exists across studies. When there is a distribution shift in the observed sequencing counts across studies, existing data integration methods may find difficulty disentangling the variation due to the batch effects from the true biological heterogeneity, which may result in overcorrection. Motivated by this need, this section investigates whether MetaDICT can prevent overcorrection when the data sets are heterogeneous across studies. The first set of numerical experiments considers an ideal setting where no batch effects exist (the measurement efficiency are the same across batches), but a biological covariate is imbalanced across studies. In this ideal setting, the variation in sequencing counts across batches is entirely due to the imbalanced biological covariate and has nothing to do with the batch effects. Similar to the previous experiments, we apply eight different data integration methods and consider the same two possible scenarios: when the confounding biological variable is not observed or observed. As expected, the imbalanced biological variable naturally leads to a distribution shift in microbial loads across studies (truth panels in Figures 3(a) and (b)). Applying data integration methods preserves such heterogeneity if the biological variable is observed (Figure 3(a)). However, when the biological variable is not observed, most existing methods tend to overcorrect the batch effects and reduce the biological heterogeneity across batches, while only MetaDICT and PLSDA-batch remain robust under such a setting (Figure 3(b)). Similar results are observed when we vary the confounding level of biological confounding variables (Figure S5) and using SparseDOSSA2 simulated data (Figure S8). This observation indicates that some existing methods might wrongly consider the effect of confounding variables as batch effects and remove it when some important confounding variables are not observed. In contrast, MetaDICT is more robust to overcorrection than existing methods because it explores the intrinsic structure of microbiome data.

**Figure 3:**
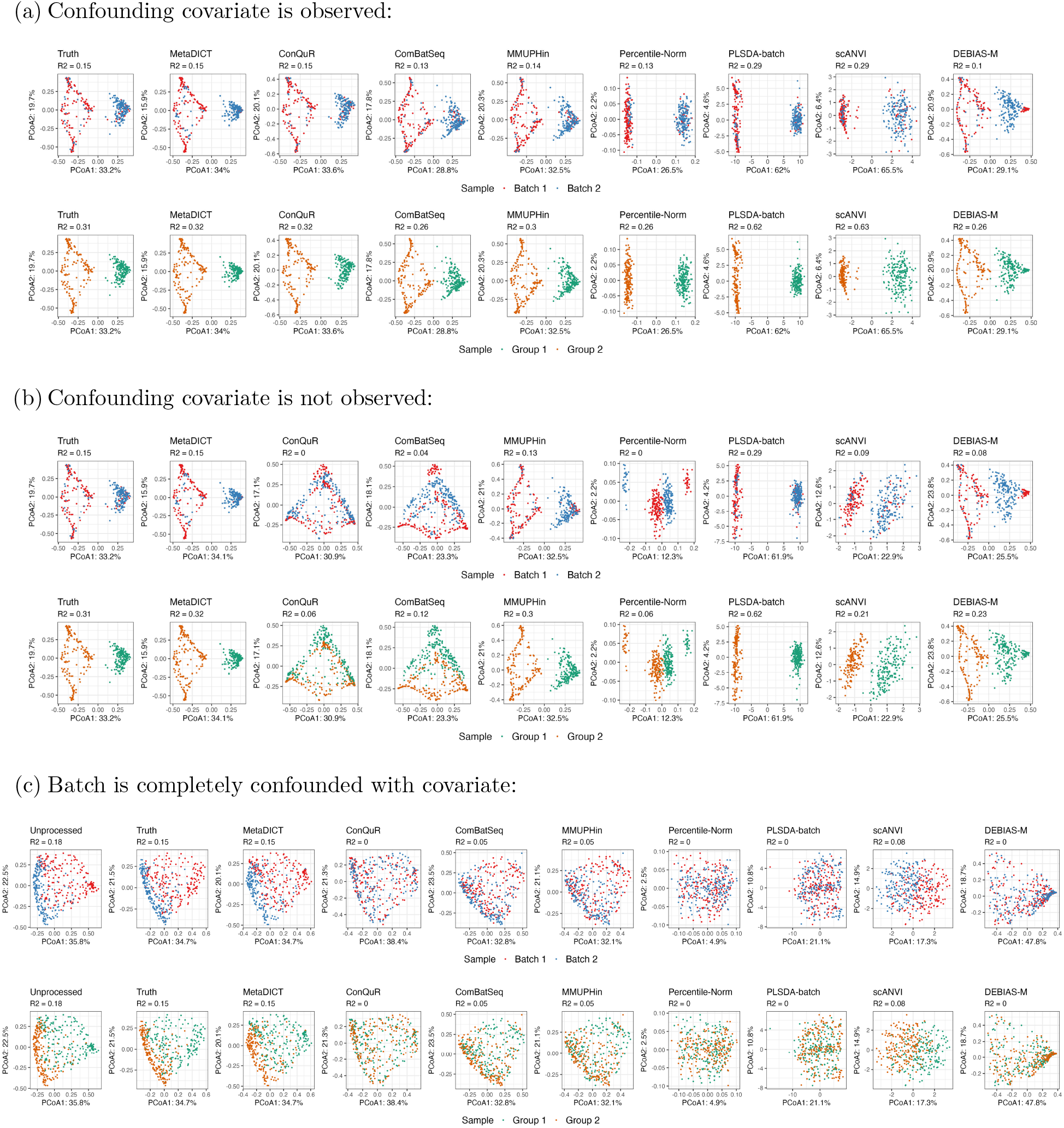
MetaDICT avoids overcorrection of batch effects in the presence of a distribution shift in absolute abundance across data sets. Figures (a) and (b) consider an ideal scenario in which there is a distribution shift in absolute abundance but no batch effect across data sets (i.e., the measurement efficiency is the same across batches). Since there is no batch effect, the observed data can be considered as ground truth as displayed in truth panels. We compare the performance of eight data integration methods via PCoA plots and *R*^2^ in PERMANOVA when the biological variable is confounded with batches. Figure (a) presents the results when the biological variable is observed in advance and used in data integrative methods, while Figure (b) presents the results when the biological variable is not observed in advance. Most existing methods tend to overcorrect batch effects (indicated by small *R*^2^ for batch variable) when the biological variable is not observed. Figure (c) compares eight data integration methods when the biological variable is completely confounded with batches (i.e., all the samples are from Group 1 in one batch and Group 2 in the other batch). All existing methods suffer from severe overcorrection because true biological variation is wrongly considered as batch effects and thus overcorrected (reduced *R*^2^ compared with the truth panel). Euclidean distance is appled for scANVI and PLSDA-batch, while Bray-Curtis dissimilarity is used for other methods. These results show that MetaDICT can perform robustly and avoid overcorrection when there is a distribution shift in absolute abundance.

The next set of numerical experiments studies a more challenging setting than the previous one, in which the batch is completely confounded with some covariates. The completely confounding case is quite common in practice. For example, the studies were conducted in different countries in the meta-analysis presented in Section 2.6, so the batch is completely confounded with the country. In the synthetic data, the binary biological variable takes 1 in one batch and 0 in the other batch, so the biological variable is completely confounding with the batch. The batch effect correction in such a setting is quite challenging as the variation in sequencing counts across batches is the interplay between batch effects and biological heterogeneity. We apply the same eight different data integration methods to correct the batch effects (Figure 3(c)). All existing methods suffer from severe overcorrection as these methods usually model the effects of batch and covariate similarly, e.g., treat the batch indicator as one of the covariates, and thus cannot disentangle the completely confounding effects. Unlike existing methods, MetaDICT can prevent overcorrection and preserve biological variation even under a completely confounding setting since effects of batch and covariate perturb the observed sequencing count differently in MetaDICT (batch effect is though multiplicative measurement efficiency and the biological variation is through shared dictionary). Besides the completely confounding case, we also design numerical experiments when the batch is confounded with biological covariates but not completely (Figure S4). Again, MetaDICT performs robustly and avoids overcorrection in this setting. All these numerical experiments demonstrate that MetaDICT better addresses the issue of overcorrection and preserves biological variation compared to existing data integration methods when biological heterogeneity exists across studies.

### 2.4 MetaDICT Reveals Hidden Structure via Embedding

Another unique feature of MetaDICT is the embedding derived from the estimated shared dictionary, and this section presents a series of numerical experiments to demonstrate its merit. We first study if the taxa embedding in MetaDICT can recover the universal structure of microbial communities. Similar to the previous section, we generate synthetic data sets with five microbial communities by modifying the real microbiome data set collected by He et al. (2018) (Figure S9). In this experiment, we consider eight community detection methods: clustering on the taxa embedding from MetaDICT, clustering on a single data set, clustering on the integrated data sets corrected by ConQuR, ComBat-Seq, MMUPHin, Percentile Normalization, PLSDA-batch and DEBIAS-M, and clustering on the combined unprocessed data sets. For a single data set, we apply clustering on each data set and present the results with the most well-separated clusters as measured by the average Silhouette score. We use the adjusted Rand index to evaluate the performance of community detection. As illustrated in Figure 4(a), the taxa embedding from MetaDICT can separate different microbial communities, while most methods tend to overestimate the number of microbial communities. We further compare the performance of these clustering methods on a wide range of experiments. Specifically, we consider various settings: 1) the confounding biological variable is observed or not (Figure 4(c)); 2) the number of data sets varies (Figure S10(a)); 3) the signal of community is different (Figure S10(b)); 4) the sample size of each data set is different (Figure S10(c)). These results suggest that the batch effect can greatly impact the microbial communities’ structure, and community detection performance relies highly on the quality of data integration. Microbial community detection could be inaccurate if we cannot correct the batch effect appropriately, such as overcorrecting the batch effect when the confounding covariate is not observed. These comparisons indicate the embedding of MetaDICT can effectively integrate the strength of multiple data sets to uncover the universal structure of microbial communities.

**Figure 4:**
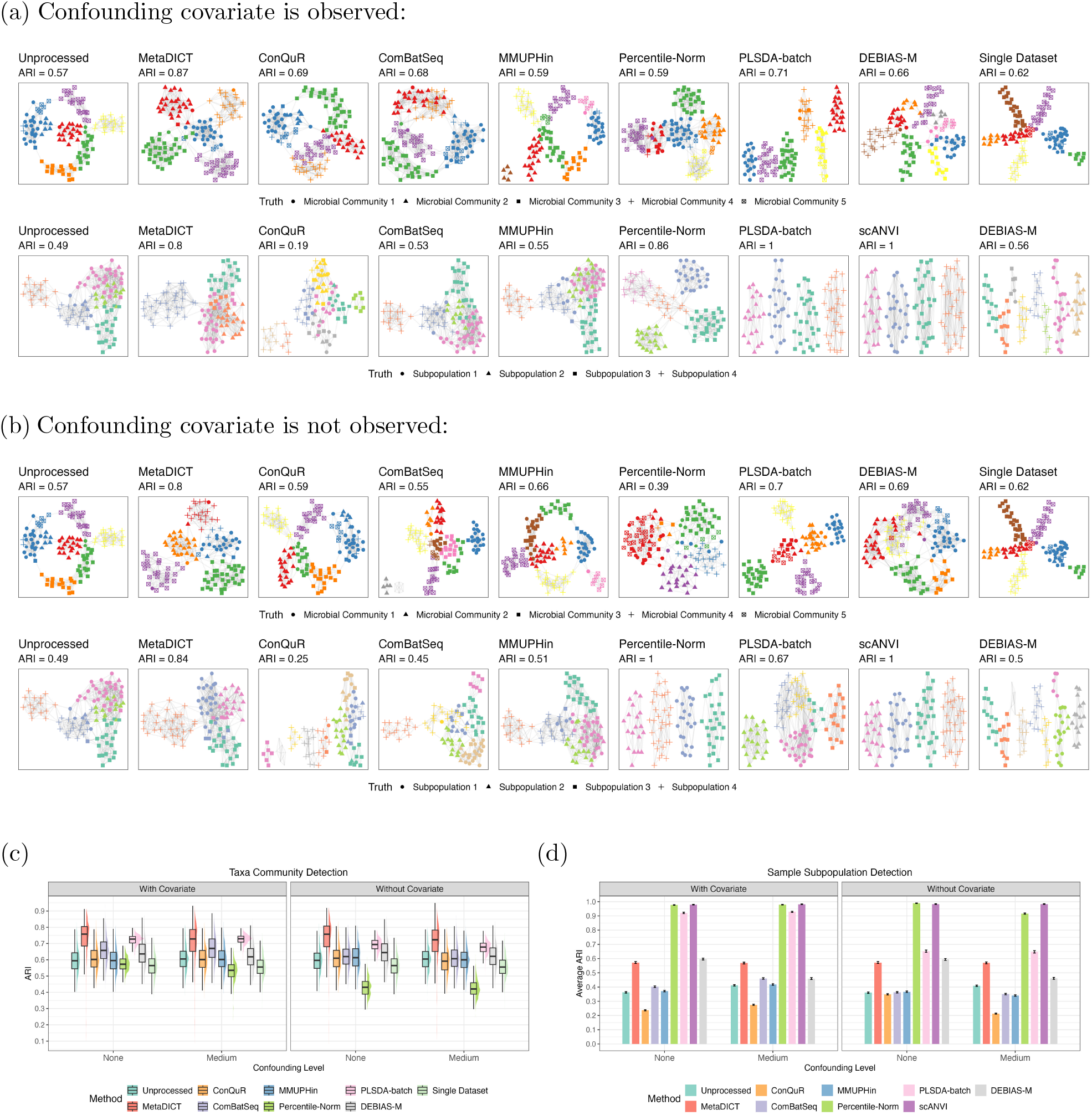
MetaDICT reveals the hidden structure of microbiome data via embedding. The embedding derived from the shared dictionary learning can provide more insight into the microbiome data. Figures (a) and (b) show examples of clustering results at taxa and samples levels when biological covariate is observed or not observed, where colors represent the detected communities and shapes represent the true communities. Adjusted Rand index is also reported in Figures (a) and (b). These examples demonstrate that the embedding in MetaDICT is effective in detecting communities at both taxonomic and sample levels. Figures (c) and (d) further compare the community detection accuracy of different methods and present adjusted Rand index in repeated experiments. Figure (c) presents the box plots of ARI for community detection at the taxa level, and Figure (d) shows the average ARI with standard error bars for community detection at the sample level. The simulated microbial communities and subpopulations are roughly equal in size. All experiments suggest MetaDICT is a reliable approach to detecting communities at taxa and samples levels.

Besides taxa embedding, MetaDICT also generates embedding at the sample level, so the next set of experiments is designed to evaluate the efficiency of sample embedding. We consider the same synthetic data sets similar to the previous set of experiments and the adjusted Rand index as the performance measure. Since we are usually interested in sample subpopulations across studies in integrative analysis, we no longer apply clustering algorithms to each individual data set. To compare different methods, we vary the access of confounding variables (Figure 4(d)), the number of data sets (Figure S11(a)), signal strength (Figure S11(b)), and sample size per data set (Figure S11(c)). Similar to the previous set of experiments, the comparisons show that the batch effects can largely perturb the community detection results at the sample level, and thus, suitable data integration is key to understanding the subpopulation of samples. In particular, the sample embedding in MetaDICT offers a concise representation of each sample and can effectively separate sample clusters even when sample covariate is not observed or confounded with batches (Figure 4(a) and (b)). In addition, the clustering at both taxa and sample levels can naturally lead to a biclustering method for integrative analysis. The results shown in Figure S9 suggest that embedding in MetaDICT is a reliable approach to unraveling the hidden communities at both taxa and sample levels.

### 2.5 MetaDICT Achieves Reliable Integrative Analysis

This section includes several numerical experiments to study how the data integration impacts the downstream integrative analysis, such as differential abundance analysis and outcome prediction. We first explore the performance of commonly used differential abundance tests on the integrated data set. Specifically, we consider the six data integration methods, including ConQuR, ComBat-Seq, MMUPHin, Percentile Normalization, DEBIAS-M, and MetaDICT, and four commonly used differential abundance tests: RDB (Wang, 2023b), LinDA (Zhou et al., 2022), ANCOM-BC (Lin and Peddada, 2020), and MaAslin2 (Mallick et al., 2021). PLSDA-batch and scANVI are not included into the comparisons as they produce embedding rather than corrected counts. In the integrative analysis, the outcome of interest in differential abundance tests is also included as an observed covariate in different data integration methods. To evaluate the performance, we consider four experiment settings: 1) there are some differentially abundant taxa and the outcome is independent of the batch (Figure S12(a)); 2) there are some differentially abundant taxa and the outcome is confounded with the batch (Figure 5(b)); 3) there is no differentially abundant taxon when there is no distribution shift in outcome across studies (Figure 5(c) and S12(b)); 4) there is no differentially abundant taxon when there is a distribution shift in outcome across studies (Figure 5(c) and S12(b)). When the outcome is independent of the batch, most data integration methods can successfully correct the batch effects, and thus, the differential abundance tests on the integrated data set can control the false discovery well and maintain a decent power. However, if the outcome of interest is a confounding variable that relies on the batch, batch effect correction becomes challenging in existing data integration methods, resulting in an inflated false discover rate and reduced power in all four differential abundance tests. Comparing the false discovery frequency of each taxon with their batch effect disturbance level suggests that the false discoveries in differential abundance analysis mainly result from remaining uncorrected batch effects (Figure 5(a)). Due to exploring the intrinsic structure of microbiome data, MetaDICT can better correct batch effect and thus lead to more reliable differential abundance analysis than existing methods when the outcome of interest is confounded with the batch. Therefore, MetaDICT is a good choice of data integration method when integrative differential abundance analysis is of interest.

**Figure 5:**
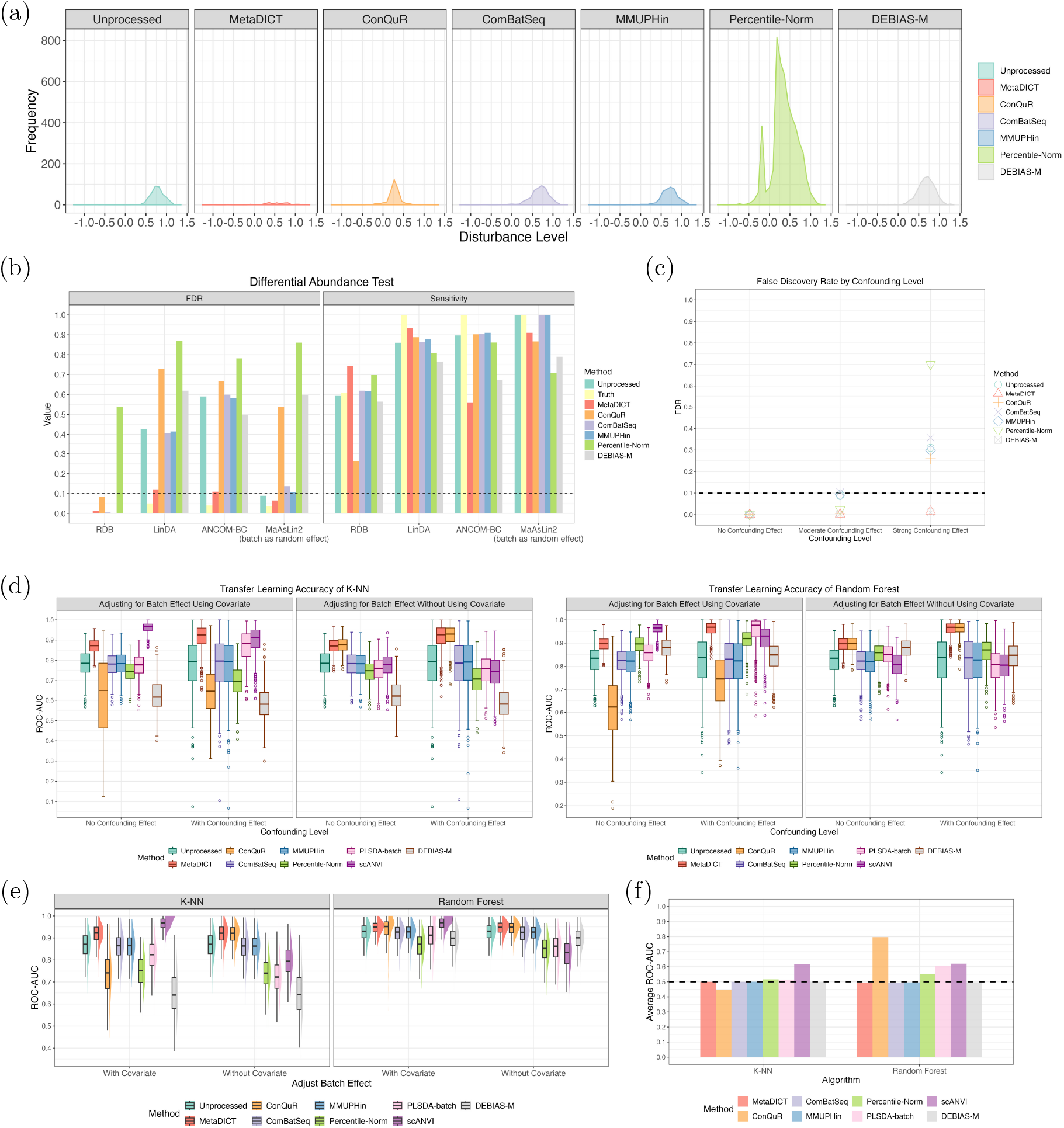
MetaDICT improves downstream integrative analyses. Figure (a) shows the frequency of false discovery at the taxa level (*y*-axis) against their disturbance level due to batch effects (*x*-axis). The results indicate that false discoveries in differential abundance tests mainly result from uncorrected batch effects, and the taxa highly affected by batch effects are more likely to be misidentified as discoveries. Figure (b) compares the sensitivity/power and false discovery rate of state-of-art differential abundance tests when data are integrated via various data integration methods. Figure (c) assesses false discovery rate inflation when the covariate of interest is independent of microbial composition but confounded with batch. The setting in Figure (b) includes differential abundant taxa, while there is no differential abundant taxon in the setting of Figure (c). The results in Figures (b) and (c) suggest that MetaDICT can better correct batch effects, leading to more reliable differential abundance analysis. Figure (d) compares the ROC-AUC of *k*-NN and random forest classifiers under a transfer learning scenario, where the training and testing data sets originate from different studies. The box covers 50% of ROC-AUC scores and whiskers extend up to 1.5 × IQR beyond upper quantile and lower quantile. Figure (e) compares the ROC-AUC of classifiers when the training and testing data sets come from the same integrated data set. To test the potential double-dipping issues, Figure (f) presents the prediction ROC-AUC when the outcome of interest is a negative control one, i.e., the outcome is fully independent of microbial compositions but is used in the data integration. These experiments suggest that MetaDICT robustly improves prediction performance while not inflating associations and leading to overly optimistic results.

The next downstream task considered in this section is outcome prediction. We consider two of the most popular classifiers in the integrative analysis: *k*-nearest neighbor classifier (*k*-NN) and random forest. When evaluating the performance by ROC-AUC and DeLong’s test (DeLong et al., 1988), we aim to answer the following two questions: 1) how does the downstream classifier perform when the batch effect is not corrected appropriately? 2) is using the same data set for data integration and classifier training safe? We consider two common prediction settings in practice to answer the first question. In the first setting, we design an experiment similar to the settings in transfer learning, where the training and testing data sets come from different studies and thus have different measurement efficiencies. The results in Figure 5(d) and S13(a) suggest that the batch effect correction between training and testing data sets is essential for building a reliable and generalizable classifier. In particular, MetaDICT leads to robust classifiers in *k*-NN and random forest while other methods present unstable performance. Besides the transfer learning setting, we also consider the second setting, where the training and testing data sets are randomly drawn from the same integrated data set (Figure 5(e) and S13(b)). The results show that classifiers trained on data sets integrated by MetaDICT achieve strong performance with *k*-NN, regardless of whether the biological covariate is observed or unobserved. This indicates that MetaDICT effectively integrates data sets across diverse scenarios. To test the potential double-dipping issues, we design an experiment with an outcome fully independent of microbial sequencing count and included it as a covariate in the data integration. The results in Figure 5(f) and S13(c) show that most data integration methods are safe when the same data set is used for data integration and classifier training. Among these methods, ConQuR, PLSDA-batch, Percentile Normalization and scANVI are more likely to achieve an over-accurate classifier in the random forest, and scANVI presents over-optimistic result for *k*-NN. A similar observation is also noted in the original ConQuR paper (Ling et al., 2022). This phenomenon, known as double dipping, often arises when a sample covariate is used repeatedly during data processing and downstream analysis, resulting in an overly optimistic association because of information leakage in data processing. Methods that heavily rely on covariate regression may inadvertently tailor or manipulate the underlying data structure to inflate the apparent association with that covariate, leading to biased results. The simulation results indicate that the double-dipping issue could be mitigated when suitable combinations of data integration methods and classifiers are used. In all the above experiments, MetaDICT consistently demonstrates robust performance in data integration, and the resulting integrated data set can significantly improve the accuracy of the downstream analysis.

### 2.6 Meta-analysis of CRC microbiome via MetaDICT

In this section, we apply MetaDICT to an integrative analysis of five colorectal cancer (CRC) metagenomic studies to further demonstrate the practical merit (Table S1). The integrative analysis aims to study the generalizable association between microbiome alterations and colorectal cancer. Although focusing on the same disease, these five studies were conducted in different countries, including the United States (US), France (FR), Austria (AT), China (CN) and Germany (DE). To reduce the technical variation, the sequencing data from these five studies were processed using consistent bioinformatics tools for taxonomic profiling. More details on sequencing and bioinformatics analysis can be found in Wirbel et al. (2019) and Supplementary Material. In the integrative analysis, we include age, gender, BMI, country, and disease status as the covariates. In particular, the study is completely confounded with the country since each study is conducted in different countries. Before conducting integrative analysis, we first explore the effect of different studies on the microbial composition and other commonly observed covariates. The results in Figure S14(c) show that the study variable had a dominant effect on the microbial profiles. Since the study is completely confounded with the county of each sample, it is difficult to tell from the explorative analysis that such a effect is due to the difference among countries or the technical variations in different studies, such as different sampling procedures, sample storage, and DNA extraction methods. In addition, there is a clear distribution shift of the observed covariates across studies (Figure S14(a)). These observations suggest that the potential batch effects could invalidate the integrative analysis, and there is a need for appropriate batch correction that can account for confounding variables and preserve biological variations. We apply MetaDICT to integrate these five studies. After processing with MetaDICT, the proportion of variability explained by the batch variable decreased from 11% to 4%, and the residual variability may reflect variation across different countries. Additionally, the proportion of variability explained by disease status increased, suggesting that true biological variation was effectively preserved.

In the integrative analysis, we first study the underlying structure of microbial communities via the taxa embedding in MetaDICT. As shown in Figure 6(b) and Table S2, the taxa embedding in MetaDICT generates a microbial interaction network and leads to 30 distinct detected microbial communities. In the microbial interaction network, 40% of the total edges are intra-phylum edges, while the expected frequency is 26% if by chance only, suggesting that the taxa from the same phylum tend to be closely connected in the network. Besides taxonomic similarities, the microbial interaction network derived from the embedding can also reflect functional similarities between taxa. For example, two subnetworks within communities 2 and 9 are butyrate generators: one subnetwork has a hub taxon *Butyricicoccus* and other taxa, like *Faecalibacterium, Blautia, Eubacterium, Dorea, Lachnospiraceae*, and the other subnetwork includes *Anaerostipes, Anaeromassilibacillus*, and *Pseudoflavonifractor*, which possess the genetic pathways necessary for converting pyruvate and acetyl-CoA into butyrate (Medvecky et al., 2018). Another highlighted example is a subnetwork within community 1 that includes several oral and periodontal pathogens, including *Porphyromonas, Peptostreptococcus, Parvimonas, Gemella, Tannerella, Lachnoanaerobaculum*, and *Solobacterium* (Hampelska et al., 2020; Sabrie et al., 2023; Ternes et al., 2020; Zwezerijnen-Jiwa et al., 2023). In addition to capturing known taxonomic and functional similarities, the detected microbial communities also suggest several interesting observations. Specifically, besides oral pathogens, community 1 also includes several other genera linked to other diseases, such as pathogens involved in opportunistic infection (*Anaerococcus, Helcococcus*, and *Providencia*) (Murphy and Frick, 2013; Lotte et al., 2015; Wie, 2015), and pathogens causing bacterial vaginosis (*Mobiluncus*) (Schwebke and Lawing, 2001), suggesting possible synergistic relationships among these pathogens. In addition, several genera from order *Eggerthellales*, including *Eggerthella* and *Adlercreutzia*, are likely to exhibit with butyrate generators in community 9, which could be explained by the hypothesis that they, as acetate consumers, could compete with microbes that convert acetate into butyrate (Noecker et al., 2023). The above observations suggest that the taxa embedding in MetaDICT can capture the taxonomic and functional similarities among taxa and offer new insights into microbial communities.

**Figure 6:**
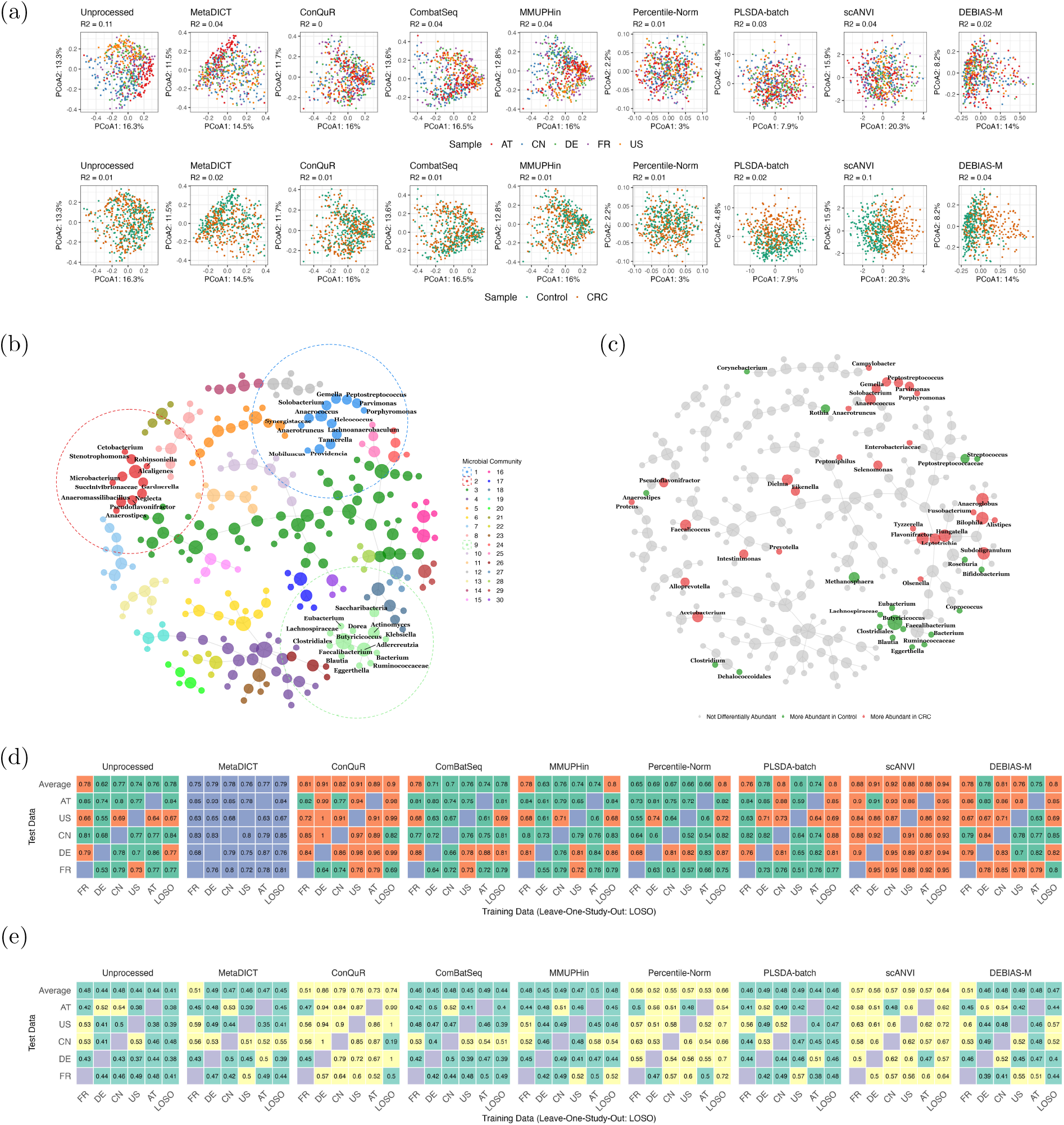
Meta-analysis of CRC microbiome via MetaDICT. Figure (a) compares PCoA plots before and after batch effect correction, while PERMANOVA *R*^2^ is shown in the subtitle. Euclidean distance is used for PLSDA-batch and scANVI while Bray-Curtis dissimilarity is applied for the rest methods. Figures (b) and (c) display the microbial interaction network derived from the taxa embedding in MetaDICT, where the community colors genera in Figure (b) and the results of differential abundance analysis color genera in Figure (c). In Figure (b), taxa circled by blue, red and green represents those from community 1, 2, 9 respectively. Figure (d) evaluates the performance of a random forest model in predicting CRC status using data processed by different data integration methods. We consider two types of classifiers: the first is trained on an individual data set and tested on another data set, while the second is trained on an integrated data set combining four studies and tested on a held-out study, referred to as LOSO. Orange in Figure (d) indicates a higher ROC-AUC than MetaDICT, while green indicates a lower ROC-AUC. Figure (e) assesses the risk of inflating associations in these data integration methods. In Figure (e), a randomly generated outcome that is independent of microbiome data is used in these data integration methods. Since there is no association between the synthetic outcome and microbiome data, the resulting classifier should perform no better than a random guess. Yellow indicates a ROC-AUC greater than 0.5, while green indicates a ROC-AUC less than 0.5 in Figure (e). Some existing methods inflate the association and lead to an over-accurate classifier.

Next, we study the association between the microbial profiles and CRC status via the integrated data set by MetaDICT. More concretely, we apply LinDA to identify differential abundance genera after adjusting the effect of observed covariates. When controlling the false discovery rate at 10%, 50 genera are detected on the integrated data set, while much fewer genera are detected on each data set (Figure S15), highlighting the increased power of integrative analysis. It is interesting to observe that the differentially abundant genera are well clustered in the microbial interaction network derived by the taxa embedding in MetaDICT (Figure 6(c)), indicating that the taxa embedding can provide insight into the pathogenic mechanism of the microbiome. In particular, most differentially abundant genera are grouped in communities 1, 9, 16, and 28, making it convenient to interpret the results from a perspective of functionality in genera. Specifically, several oral pathogens, including *Porphyromonas, Peptostreptococcus, Parvimonas, Gemella*, and *Solobacterium*, in the microbial community 1 are more abundant in CRC samples than control ones, suggesting a close connection between oral microbiome and progression of colorectal cancer (Flemer et al., 2018; Mo et al., 2022). Microbial community 16 includes three differentially abundant genera, i.e., *Fusobacterium, Bilophila*, and *Alistipes*, that can produce hydrogen sulfide, a key signaling biomolecule in colorectal cancer (Basic et al., 2017; Tilg et al., 2018; Lin et al., 2023). *Flavonifractor* and *Tyzzerella* from microbial community 28, well-known biomarkers of colorectal cancer (Yang et al., 2021; Wu et al., 2021b), are reported as differentially abundant genera. While the differentially abundant genera from the previous three communities associate with colorectal cancer positively, the ones from microbial community 9 show a negative association. The differentially abundant genera from microbial community 9 can mostly produce butyrate, which is a primary energy source for colonocytes and protects against colorectal cancer and inflammatory bowel diseases (Lopez-Siles et al., 2017; Luo et al., 2023). Besides these well-grouped genera, we also discover several isolated differentially abundant genera that can provide insight into the difference between CRC and control samples. For example, three oral-origin genera, *Eikenella, Anaeroglobus*, and *Rothia*, that are usually linked to periodontitis disease (Karim et al., 2013; Bao et al., 2017; Mazurel et al., 2023) show a significant association with the development of colorectal cancer, further underscoring the close relationship between the oral microbiome and colorectal cancer (Zepeda-Rivera et al., 2024). The above discoveries highlight the key role of MetaDICT in increasing accuracy and interpretability for downstream statistical analysis.

Lastly, we aim to use the integrated data set to train a random forest classifier that uses microbial profiles to predict CRC status in disease diagnosis. To compare the generalizability, we evaluate the performance of the classifier (measured by ROC-AUC) on the data set from one study and train the classifier on the individual data set from other studies or the integrated data set of other studies. The results are summarized in Figure 6(d). Due to the interplay between batch effects and heterogeneity of data sets, the classifiers trained by the unprocessed data cannot generalized well, e.g., the classifier trained on DE can only achieve a ROC-AUC of around 53% in FR (almost randomly guessing). After correcting batch effects by MetaDICT, resulting classifiers’ generalizability is mostly improved. For instance, when the training data are from DE and testing data are from FR, ROC-AUC increases to 76% after MetaDICT processes the data sets. As expected, the classifier training on the integrated data set is more accurate than training on the individual data set. Again, the performance of the resulting classifier is enhanced when the batch effects are corrected on the integrated data set. We also compared the results across various integrative methods using DeLong’s test, and the results are summarized in Figure S16(a). MetaDICT, ConQuR, DEBIAS-M, and scANVI achieved higher ROC-AUC values among these methods. However, ConQuR and scANVI demonstrated a greater risk of inflated ROC-AUC, as they produced overly optimistic results in predicting a randomly simulated variable that is independent of microbiome compositions (Figure 6(e) and S16(b)). In addition to the analysis at the genus level, we also train classifiers using data at the species level. After the data is preprocessed in the same way as Wirbel et al. (2019), we consider two classifiers for CRC status prediction: random forest and Lasso logistic regression, which is adopted in Wirbel et al. (2019). The results summarized in Figure S17 suggest that ROC-AUCs of Lasso logistic regression are improved by MetaDICT in most cases, and the random forest classifier trained on MetaDICT-corrected data is a more accurate classifier than the one trained in Wirbel et al. (2019) (the average ROC-AUC is increased from 82.9% to 85.6%). These comparisons underscore the importance of batch effect correction in increasing the generalizability and prediction accuracy of integrative analysis.

### 2.7 Integrative Analysis of Microbiome in PD-1 Treated Patients

We further illustrate MetaDICT using a meta-analysis of five 16S rRNA microbiome data sets of anti-programmed cell death protein-1 (PD-1) treated patients with melanoma (Table S3). The goal of this integrative study is to identify consistent microbial signatures associated with clinical response and build a predictive model for immunotherapy outcomes. In these five studies, 174 stool samples in total were collected from PD-1-treated patients at the beginning of therapy but the samples are sequenced by different protocols in different studies. Despite different protocols, all the sequence data sets are processed with the same bioinformatics pipeline (Shaikh et al., 2021). The number of responders and non-responders is highly imbalanced across five studies (Figure 7(b)), indicating a potentially strong confounding effect from the batch. The PCoA plots in Figure 7(a) show the samples from different studies are well separated due to the strong batch effects. Similar to previous sections, we apply eight data integration methods to these five data sets. After data integration, only MetaDICT and ConQuR significantly reduce the variation attributed to batches, while the MetaDICT also preserves, but does not artificially boost, the heterogeneity due to the imbalanced response status across studies.

**Figure 7:**
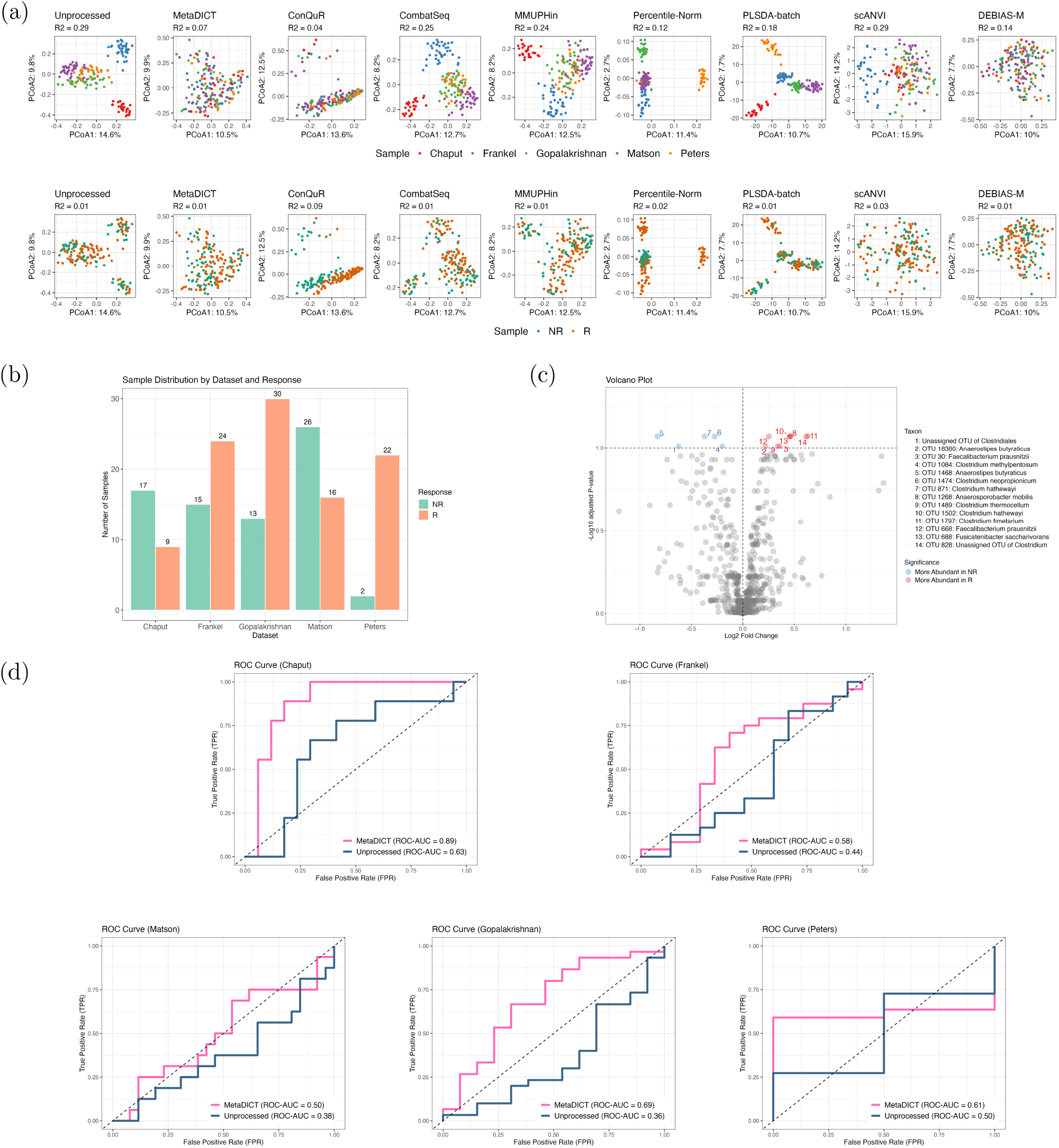
Integrative Analysis of Microbiome in PD-1 Treated Patients via MetaDICT. Figure (a) compares PCoA plots before and after batch correction using integrative methods, while PERMANOVA *R*^2^ is shown in the subtitle. Euclidean distance is used for PLSDA-batch and scANVI while Bray-Curtis dissimilarity is applied for the rest methods. Figure (b) shows the number of responders and non-responders in each data set, indicating a highly imbalanced outcome of interest across studies. Figure (c) presents volcano plot of differentially abundant taxa detected on the data set integrated by MetaDICT. Figure (d) shows ROC curves for neural network classifiers trained on four data sets and evaluated on the held-out data set, using both unprocessed data and data integrated with MetaDICT. The results show that MetaDICT can greatly correct the batch effect and boost the prediction accuracy of neural network classifiers.

Next, we use differential abundance analysis to study microbial signatures associated with therapy response. Specifically, we apply LinDA to the microbial OTU count data tables with 881 taxa when the false discovery rate is controlled at 10%. When the batch effects are not corrected, 129 taxa are reported as differentially abundant ones if we do not consider the batch in the analysis, while none of the taxa are significant if we incorporate batches as random factors. The inconsistent results suggest that the batch effects highly impact the differential abundance analysis, and there is a need for appropriate batch effect correction. We identify 14 differentially abundant taxa after MetaDICT integrates data sets from different studies (Figure 7(c)). Specifically, the differentially abundant taxa include ones from *Faecalibacterium prausnitzii, Anaerostipes butyraticus*, and five other species of the genus *Clostridium*, which produce short-chain fatty acids that have a large impact on the immune system (Wu et al., 2021a). The effect of short-chain fatty acids (particularly acetate, propionate, and butyrate) on host immunity and cancer immunotherapy has been discussed in various cancers (Zhang et al., 2019; Peng et al., 2020). The close connection between *Faecalibacterium prausnitzii* and immunotherapy response is well documented in the literature (Gopalakrishnan et al., 2018; Gao et al., 2023; Bredon et al., 2024; Formiga and Sokol, 2024). These discoveries demonstrate the efficiency of MetaDICT in batch effect correction and improving downstream analysis.

Lastly, we aim to evaluate whether data integration can improve the prediction of the immunotherapy response. We choose a two-layer neural network classifier in this section since the previously used random forest cannot fit these data sets very well and can only achieve mediocre performance for most methods without inflating the association in negative control experiments (Figure S18(a) and (b)). The accuracy of the classifier is evaluated by leave-one-study-out cross-validation, in which the classifier is trained on four out of five data sets and tested on the remaining data set. To avoid the inflated prediction resulting from data integration, we first design an experiment with a completely random simulated outcome when two-layer neural network classifiers are trained on the data set integrated by different methods (Figure S18(c)). The results show that ConQuR and scANVI highly inflate the prediction of neural network classifiers and suffer a double-dipping issue on this data set, while MetaDICT and MMUPHin prevent the over-accurate classifier. Therefore, we train a neural network classifier to predict the immunotherapy outcomes using the data set integrated by ComBat-Seq, MMUPHin, Percentile Normalization, PLSDA-batch, DEBIAS-M, and MetaDICT. We can observe a significant improvement in accuracy after these data integration methods correct the batch effects (Figure 7(d)). The comparisons suggest that MetaDICT and DEBIAS-M have a more robust performance than other methods (Figure S18(d)).

## 3 Discussion

This paper presents a new data integration method for microbiome studies, called MetaDICT. While existing methods mainly explore the relationship between microbial composition and observed covariates, MetaDICT also utilizes the intrinsic structure of microbiome data via shared dictionary learning. By taking advantage of the intrinsic structure, the new approach can better correct batch effects and preserve biological variation than existing methods, especially when unmeasured confounding variables exist, studies are highly heterogeneous in populations or the batch is completely confounded with some covariates. In addition to batch effect correction, MetaDICT generates the embedding at both taxa and sample levels to unravel the hidden structures of microbiome data. Our comprehensive numerical experiments show that the corrected count tables and embedding offered by MetaDICT can improve the commonly used integrative analysis, such as community detection, differential abundance analysis, and outcome prediction.

MetaDICT explores the assumptions and intrinsic structures in multiplicative batch effects, shared dictionary in absolute abundance, and measurement efficiency’s smoothness with respect to the taxa similarity. While these assumptions have been discussed in various aspects of microbiome literature, putting them together in MetaDICT offers a fresh perspective on how intrinsic structure can contribute to batch effect correction and data integration in microbiome data. We believe this new perspective will inspire the development of more exciting data integration methods for microbiome data. Although our discussion mainly focuses on the standard setting of microbiome study, MetaDICT can also be potentially applied to other settings, such as multiple samples observed from the same individuals and longitudinal samples, since the shared dictionary assumption can also hold in these settings. Moreover, while MetaDICT is designed for microbiome data, its framework for modeling batch effects is flexible enough to potentially work for other types of data. For example, the model adopted by MetaDICT can also be used for single-cell RNA sequencing data (Peng et al., 2021). This versatility opens up a world of possibilities for MetaDICT’s application in a broad range of data sets.

A crucial assumption in MetaDICT is that the microbial load matrix can be approximated by a product of two matrices, one of which is shared dictionary. Despite the potential nonlinear interactions among microbes, such a matrix factorization approximation in the microbiome data has been well discussed and validated in the literature (Sankaran and Holmes, 2019; Cao et al., 2020; Kim et al., 2023). A simple diagnostic tool for such an assumption is to evaluate the singular values of the sequencing count matrix in each study and see how fast the singular values decay. For example, Figure S19 displays the plots of singular values for the data sets in Sections 2.6 and 2.7. When a fast decay is observed, the matrix factorization approximation is well-held, and MetaDICT can be applied. It would also be interesting to explore the potential generalization of MetaDICT to accommodate a broader range of nonlinear settings in future work.

The workflow in MetaDICT represents just one way to utilize the intrinsic structures in microbiome data. However, there are likely more intrinsic structures to be explored or alternative ways to explore the same structure. For instance, the nonconvex formulation could be substituted with convex methods to exploit the shared dictionary in absolute abundance. Another possibility is replacing the graph Laplacian with a total variation on a graph to measure the overall smoothness of measurement efficiency (Sadhanala et al., 2016). Due to the nonconvex formulation, the results in MetaDICT could also be influenced by the choice of optimization algorithm, as these algorithms can introduce implicit bias (Gunasekar et al., 2018). This underscores the possibility for further investigation into more efficient workflows for correcting batch effects and integrating data sets than MetaDICT, inspiring the development of similar methods in the field.

Although MetaDICT formulates the batch effects as multiplicative bias in the sequencing process, MetaDICT is flexible to accommodate other types of formulation for batch effects. For example, if we would like to consider extra additive bias beyond multiplicative bias, we could consider the following optimization problem in the shared dictionary learning step

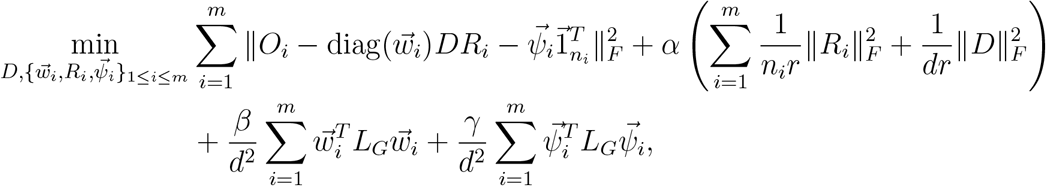

where 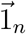 is an *n*-dimensional vector of ones, and *γ* is another tuning parameter. Here, 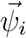 represents the potential additive bias in the sequencing process.

Compared to most existing methods, MetaDICT can better avoid overcorrection and preserve the biological variation of each data set, especially when data sets are highly heterogeneous or the biological covariate is completely confounded with batches. We demonstrate the efficacy of MetaDICT in correcting batch effects, reducing false discoveries, and enhancing statistical power in both simulations and real integrative analyses. We anticipate that MetaDICT can facilitate the identification of consistent microbial signatures and enable reliable downstream analysis results in future integrative studies.

## 4 Methods

### 4.1 A Model for Microbial Sequencing Data

In microbiome studies, researchers usually measure the abundance of different microbes in each sample by collecting their microbial sequencing data (Lozupone et al., 2007; Vandeputte et al., 2017). However, the observed sequencing count via commonly used sequencing techniques cannot accurately reflect microbial loads (absolute abundance) in each sample due to the bias introduced in the sequencing procedure (McLaren et al., 2019; Wang, 2023a). To characterize such bias, we consider a simple mathematical model to connect the absolute abundance and observed sequencing count

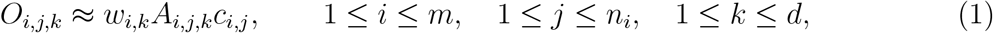

where *O*_*i,j,k*_ represents the observed sequencing count of taxon *k* in sample *j* from study *i*, and *A*_*i,j,k*_ represents the corresponding microbial loads in the sample. There are *m* studies, *n*_*i*_ samples in study *i*, and *d* different microbial taxa. In the above model, the sampling efficiency *c*_*i,j*_ characterizes the sample-specific bias related to technical factors such as sequencing depth (Robinson and Oshlack, 2010; Conesa et al., 2016; Wang, 2023b). On the other hand, *w*_*i,k*_ is the measurement efficiency of taxon *k* in study *i*. The taxon-specific bias characterized by *w*_*i,k*_ can be related to the distinct combinations of chemicals and reagents used in different DNA extraction protocols (Morgan et al., 2010), PCR binding and amplification efficiencies in a distinct agreement between primer design and sequences (Polz and Cavanaugh, 1998), and coverage variability with different sequencing platforms (Harismendy et al., 2009). While the above model seems straightforward, several studies have validated the multiplication effects of sample-specific and taxon-specific bias represented in (1) (Conesa et al., 2016; McLaren et al., 2019).

In each sample, the microbes rarely live isolated but coexist as a complex ecosystem (Woyke et al., 2006; Chaffron et al., 2010). In particular, multiple microbial communities naturally form through interactions such as nutritional cross-feeding, co-colonization, and competition (Faust et al., 2012). This observation indicates that the microbes’ abundance could change in a highly correlated way. We consider a mixed membership model for the microbial abundance profiles to capture such structural patterns in abundance. Specifically, let us write the absolute abundance vector in sample *j* from study *i* as 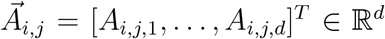. We can approximate 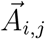 as

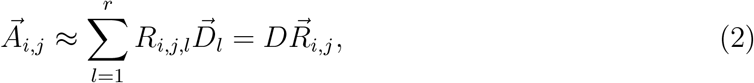

where each 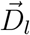 represents a group of microbes of which abundances change in a highly correlated way, and the representation 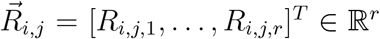 characterizes the amplitude of changes along these *r* groups. The combination of these *r* groups of microbes, 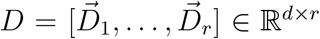, is called a shared dictionary as it represents the universal structural pattern in microbial abundance across the data sets. Although the shared dictionary is universal across studies, the distribution of the representation 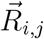 could differ for different data sets due to the potential heterogeneity across data sets. While the above type of matrix factorization structure has been widely used in the model for a single microbiome data set (Sankaran and Holmes, 2019; Cao et al., 2020; Kim et al., 2023), we extend it as a universal structure across multiple data sets. The design of the shared dictionary for absolute abundance allows for the separation of the batch effects (captured by *w*_*i,k*_) and biological variation in absolute abundance (captured by 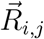), even in a completely confounding setting.

The sequence structure of each taxon mainly determines the taxon-specific measurement efficiency *w*_*i,k*_. For example, DNA extraction efficiencies are related to the cell walls and membrane structure (Krsek and Wellington, 1999; Carrigg et al., 2007), and both PCR binding and sequencing efficiencies are related to the arrangement or organization of nucleotides, especially the GC contents (Polz and Cavanaugh, 1998; Benjamini and Speed, 2012). This observation suggests that sequences with close structures can exhibit similar capture efficiencies, that is,

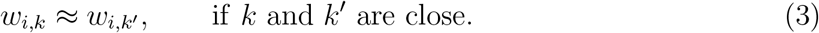

In other words, the measurement efficiency is smooth with respect to the similarity among taxa. The approximations in (1), (2), and (3) are three key assumptions in our model for microbial sequencing data.

### 4.2 Data Integration via Shared Dictionary Learning

We collect *m* microbiome data sets from different studies in the data integration. In the *i*th data set, we observe the microbial sequencing count 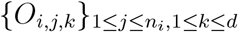 from *n*_*i*_ samples and *d* common taxa and the covariates 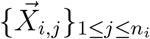 from *n*_*i*_ samples, such as age and blood type. We assume that the observed microbial sequencing data follow the model introduced in the previous section (or the three assumptions in (1), (2), and (3)). The model in (1) suggests that our observed sequencing counts are disturbed by the unobserved measurement efficiency, and thus, it is difficult to distinguish whether the variation across multiple data sets results from the confounding effect of measurement efficiency or true biological variation. Therefore, the main challenge of data integration is to remove the effect of heterogeneous measurement efficiency (also known as batch effects) while maintaining the true biological variation in the microbial loads across different data sets.

We introduce a two-stage method for integrating data sets from multiple studies to address the challenge. The first stage obtains an initial estimator by adjusting the observed covariates, while the second stage further refines the estimation by exploring the shared dictionary of microbial sequencing data. We assume *c*_*i,j*_ = 1 for the simplicity of analysis in this section, as the sampling efficiency can be normalized by the commonly used normalization methods (Bullard et al., 2010; Paulson et al., 2013; Love et al., 2014; Yuan and Wang, 2023).

#### Stage 1: Initial Estimation by Covariate Balancing

The conventional wisdom in data integration is that the difference in sequencing count distributions is due to batch effects after adjusting the effects of all possible confounding variables, and we shall remove such differences before integrating data sets (Gibbons et al., 2018; Zhang et al., 2020; Ma et al., 2022; Ling et al., 2022; Wang and Lê Cao, 2023). The main assumption behind such a strategy is that the conditional expectations of absolute abundance are the same across different data sets, that is,

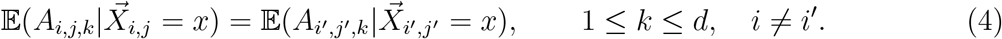

Together with the assumption in (1), this assumption naturally leads to

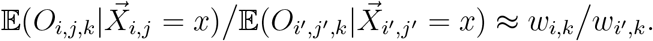

Consequently, we can adopt treatment effect estimation techniques in causal inference literature to estimate the ratio *w*_*i,k*_*/w*_*i′,k*_. In particular, the weighting method, one of the most widely used covariate balancing techniques, is particularly suitable for the estimand with the ratio form. The idea of the weighting method is to assign weights *e*_*i,j*_ to each sample so that the weighted distributions of covariates 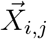 are balanced across the data sets (Imbens and Rubin, 2015). There are several ways to estimate weights in the literature, including inverse-probability weighting (Rosenbaum and Rubin, 1983; Robins et al., 2000; Hirano and Imbens, 2001) and balancing weighting (Hainmueller, 2012; Imai and Ratkovic, 2014; Zubizarreta, 2015; Chan et al., 2016; Yu and Wang, 2024). With the estimated weights *e*_*i,j*_, we can estimate the ratio *w*_*i,k*_*/w*_*i′,k*_ by

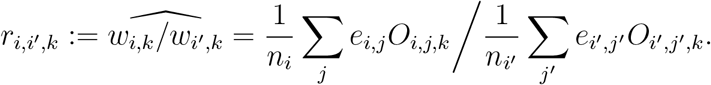

The definition suggests *r*_*i,i,k*_ = 1. The ratio estimator *r*_*i,i′,k*_ offers a straightforward way to correct batch effects and integrate data. Specifically, the corrected and comparable microbial sequencing count can be defined as

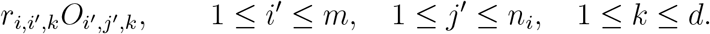

The above strategy can successfully correct the batch effects when all the confounding variables are observed and the assumption in (4) is satisfied. However, it is common that only a few covariates are observed across all data sets, and there are several important unobserved confounding variables, such as lifestyle. When the assumption in (4) is invalid, the unobserved confounding variables’ effect might be characterized as the batch effect, leading to overcorrection of the batch effect and wrong reduction of the true biological variation. To address such an issue, we propose to refine the above estimation by exploring the intrinsic structure of microbial sequencing data.

#### Stage 2: Estimation Refinement by Shared Dictionary Learning

The first intrinsic structure explored here is the shared dictionary structure introduced in assumption (2). The shared dictionary structure and assumption in (1) suggest that the matrix of observed sequencing count in each data set, 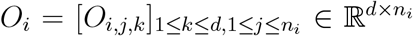, can be factorized as a product of three matrices

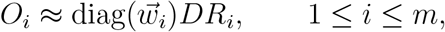

where 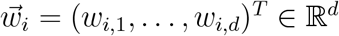 is the vector of measurement efficiency in the *i*th data set, 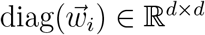 is a diagonal matrix with diagonal entries as 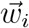, and 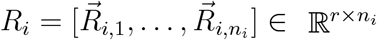 is the representation matrix of *i*th data set. As one would prefer a concise model, it is natural to assume both *D* and *R*_*i*_ have some low-rank structure. To capture such shared and low-rank dictionary structure, we can consider the following loss function

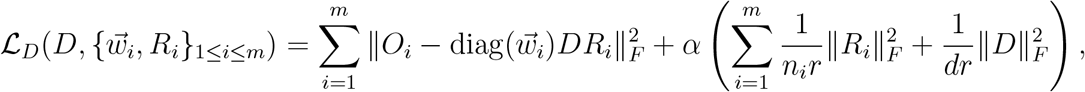

where ∥ · ∥_*F*_ is the Frobenius norm of a matrix. The first term in the above loss function measures the discrepancy between the observation and matrix product, and the second term promotes the low-rank structure in *D* and {*R*_*i*_}_1≤*i*≤*m*_ (Srebro et al., 2004). Another intrinsic structure we can explore here is the smoothness of measurement efficiency as characterized by the assumption in (3). To characterize the similarity of the sequence structure of taxa, we can construct a graph *G* = (*V, E*) such that each vertex is a taxon and two taxa are connected if they are similar in the sequence structure. For example, the sequence similarity between taxa can be measured by phylogenetic distance or taxonomic similarity. Given the graph *G*, we can consider the graph Laplacian to measure the overall smoothness of measurement efficiency with respect to the graph

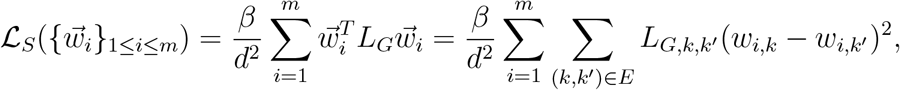

where *L*_*G*_ is the graph Laplacian matrix, and *L*_*G,k,k*′_ is the weight of the edge (*k, k*^*′*^). Putting these two loss functions together yields the following optimization problem

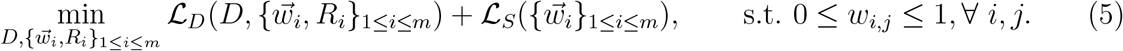

In the above optimization problem, the estimation of measurement efficiency is further refined by treating the shared dictionary as an anchor. To solve the above optimization problem, we can set the results from stage 1 as the initial point so that the estimation for measurement efficiency can be further refined. Since the low-rank matrix factorization has the effect of denoising (Chi et al., 2019), our final corrected and comparable microbial sequencing data are 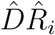 for 1 ≤ *i* ≤ *m*, where 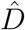 and 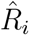 are the outputs of the above optimization problem. Notably, the output of the nonconvex optimization problem in (5) highly relies on the choice of initial points, so the estimation in stage 1 is also critical.

### 4.3 A Practical Workflow of MetaDICT

While the last section introduces a general methodology for data integration, we present a detailed workflow used in all numerical experiments.

1. **Preprocessing** Before correcting batch effects, we apply normalization methods to remove the effect of unobserved sampling fraction *c*_*i,j*_ for microbiome data sets. For ASV/OTU data, we recommend using RSim (Yuan and Wang, 2023), and for data at other resolutions, we apply the Upper Quantile (UQ) method (Bullard et al., 2010). In particular, ASV/OTU data sets typically exhibit a high prevalence of zeros (greater than 95%), where RSim performs better than UQ; conversely, when taxa are aggregated at higher taxonomic levels, UQ is faster and more robust. Despite our recommendation, other normalization methods may also apply. When a few taxa are missing in some data sets, we fill the missing values with zeros, a commonly used technique in low-rank matrix completion. The matrix factorization employed in shared dictionary learning can aid in recovering these missing values (Chi et al., 2019). The effectiveness of this zero-filling strategy relies on the overlap of dominated taxa across the data sets. When poor overlap among dominated taxa is observed, data integration becomes quite a difficult task. We do not recommend extensive filtering in MetaDICT as it can break the shared dictionary structure among taxa and reduce accuracy.
2. **Covariate Balancing** In the initial estimation, we choose the inverse-probability weighting to balance the distribution of covariates (Rosenbaum and Rubin, 1983; Robins et al., 2000), where the propensity score is estimated by logistic regression. With estimated weights, the ratio of measurement efficiency is estimated by a weighted average of sequencing count. While MetaDICT can preserve biological variation despite unobserved confounding variables, we recommend that users include all the shared covariates across data sets in the balancing step to reduce computational time and improve performance in the next step.
3. **Shared Dictionary Learning** In the optimization problem of (5), we use the phylogenetic tree to construct a *p*-nearest nearest neighbors graph for taxa. When the phylogenetic tree is unavailable, the graph can be constructed by connecting the taxa within the same taxonomic group. The optimization problem is solved by the limited-memory Broyden-Fletcher-Goldfarb-Shanno algorithm (L-BFGS) (Byrd et al., 1995), a quasi-Newton method particularly suitable for solving large non-linear optimization problems subject to simple bounds. The initial point for 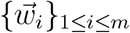 is constructed by the estimation in the previous step, while the initial point for dictionary *D* and representation matrix {*R*_*i*_}_1≤*i*≤*m*_ are the singular value decomposition of concatenating corrected microbial sequencing data matrix 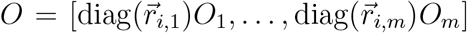. Optionally, to preserve the original microbiome data structure, MetaDICT converts the small values or negative values resulting from optimization to zeros when they are zeros in the original data set.

In the above workflow, we also need to choose four tuning parameters:

- The parameter *p* represents the number of nearest neighbors to construct the graph *G*. In our numerical experiments, we select *p* from 5 to 10.
- *r* is an important parameter in the size of *D* and {*R*_*i*_}_1≤*i*≤*m*_. In our numerical experiments, we select *r* as the estimated rank of *O*. Since we put some penalty in place to promote a low-rank structure, the rank of the resulting *D* and {*R*_*i*_}_1≤*i*≤*m*_ might be smaller than *r*.
- Parameters *α* and *β* control the importance of the penalty in the optimization problem. The effects of *α* and *β* are illustrated in Figure S1. The large tuning parameter *α* can promote the low rank of shared dictionary *D* and increase convergence time, which is usually used in a setting with strong heterogeneity across data sets. A larger tuning parameter *β* can enforce more smoothness of measurement efficiency. In practice, we recommend a large *β* when the graph is constructed by a phylogenetic tree and a small *β* when the taxonomic information constructs the graph. We recommend the choices of *β* ≤ *α <* 1. In our numerical experiments, we usually choose *β* = 0.01 when taxonomic information characterizes the taxa’s similarity and increase it to *β* = 0.1 if the phylogenetic tree is available. After *β* is selected, we often choose *α* between *β* and 1 and prefer a large *α* when we can achieve the similar fitness of the model.

Since the optimization problem in MetaDICT is nonconvex, we do not expect to converge a global minimizer but a local minimizer. To check whether a local minimizer is satisfied, we suggest evaluating two quantities

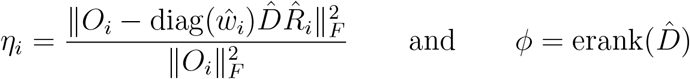

where erank is the effective rank of a matrix (Roy and Vetterli, 2007), a more computable way to measure the dimensionality of the shared dictionary. *η*_*i*_ is expected to be between 0 and 1 and measures how well the estimated measurement efficiency and shared dictionary can fit the observed data in the *i*th study. We always hope that a small *η*_*i*_ is achieved and that a smaller *ϕ* is better given small *η*_*i*_. We achieve a satisfactory convergence when we observe small *η*_*i*_ and small *ϕ*. When large *η*_*i*_ is observed, we can tune the parameters *α* and *β* for better convergence. In our numerical experiments, we usually expect satisfactory convergence when *η*_*i*_ ≤ 10^−8^ for 1 ≤ *i* ≤ *m* and gradually increase *α* until the smallest *ϕ* is observed and *η*_*i*_ ≤ 10^−8^ for 1 ≤ *i* ≤ *m*.

The computational time of MetaDICT depends on many factors, including the number of taxa, the number of samples, the choice of optimization algorithm, and tuning parameter choices. The computational complexity can be written as *O*((*d* + ∑_*i*_ *n*_*i*_)*r*), so we can expect the algorithm to be efficiently applied to large data sets as it scales linearly with the number of taxa and samples. The choice of tuning parameters also greatly impacts the number of iterations needed for optimization, affecting the computation time.

### 4.4 Batch Correction When Shared Dictionary is Available

In practice, the data set, *O*_*m*+1_, from a new study may become available after we integrate the data sets, *O*_1_, …, *O*_*m*_, from *m* studies and train a machine learning model on the integrated data set. Due to the batch effect, the trained machine learning model from *m* studies cannot be applied to *O*_*m*+1_ directly. Can we correct the batch effect for *O*_*m*+1_ without processing *O*_1_, …, *O*_*m*_ in such a case? We now show an algorithm to correct the batch effect when the shared dictionary is available. Suppose 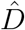 is the shared dictionary estimated by *O*_1_, …, *O*_*m*_.

1. **Preprocessing:** Apply the same normalization method to *O*_*m*+1_.
2. **Covariate Balancing:** Apply the weighting method to estimate the initial measurement efficiency with the covariates shared by new and previous studies.
3. **Shared Dictionary Learning:** We solve the following optimization problem:

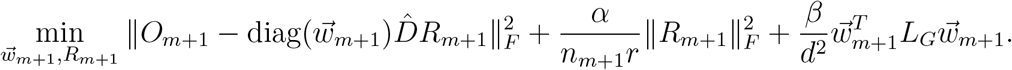

The corrected sequencing count table is 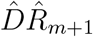, where 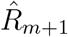 is the result of the optimization problem above.

The above pipeline allows batch effects to be quickly corrected for the new data set without retraining the shared dictionary on all the data sets. This pipeline can work well when a diverse and representative source of data sets estimates the shared dictionary 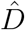. If *O*_1_, …, *O*_*m*_ are not representative enough, we recommend applying MetaDICT to all data sets *O*_1_, …, *O*_*m*+1_ to correct batch effects.

### 4.5 Representation Learning in MetaDICT

While the proposed method can correct the batch effects robustly, the output from our shared dictionary learning can provide more insight into the microbial sequencing data. Like other matrix factorization methods, the resulting dictionary *D* and representation matrix {*R*_*i*_}_1≤*i*≤*m*_ can naturally lead to the embedding of taxa and samples. It is worth noting that the decomposition of *D* and *R*_*i*_ is unique up to a rotation. We apply Varimax, a technique to find a good rotation, to make the shared dictionary interpretable (Kaiser, 1958).

#### Embedding of Taxa

As discussed in the model for microbial sequencing data, each dictionary column represents a direction in which the microbes are likely to change systemically. This interpretation suggests that we can consider each dictionary row as taxon’s embedding. When two rows/representations are similar, the abundance of these two taxa constantly changes similarly. These embeddings of taxa can provide more understanding of taxa. For example, taxa representation can help detect the taxa communities. In practice, we can select the first *r* columns of *D* as the taxa representations, where *r* is the elbow point of the singular values of *D*. Given this new taxa representations, we can construct a *k*-nearest neighbor graph via the distance matrix of taxa representations and apply some clustering algorithms, such as spectral clustering, the Louvain algorithm (Blondel et al., 2008), or the Walktrap algorithm (Pons and Latapy, 2005). These detected taxa communities can help reveal universal microbial co-occurrence relationships in multiple data sets.

#### Embedding of Samples

Besides the shared dictionary, the representation matrix can provide an important source for understanding these microbial data. In particular, each column of the representation matrix can be interpreted as a sample representation, as it reflects the coordinate in the low-dimensional space spanned by the shared dictionary. Due to the corrected batch effects, these sample representations are cast in a common space and thus are comparable across multiple data sets. Therefore, these representations can be directly useful in the downstream sample analysis, such as sample clustering and classifier construction.

### 4.6 Simulation Setup and Implementation of Existing Methods

All synthetic data sets are generated using the microbiome data set collected by He et al. (2018). The absolute abundance/microbial loads of each simulated data set are generated by perturbing the observed sequencing count in the real data set, so the characteristics of the microbiome data set, such as sparsity and high skewness, can be well preserved in the synthetic data sets. Specifically, a subset of samples are randomly selected from the real data set, and each sample’s observed/unobserved covariates are generated from some pre-specified probability distributions, such as the Bernoulli distribution or uniform distribution. The simulated microbial loads are the modified counts of the selected samples after we insert the signal of the observed/unobserved covariates and add small random perturbation, such as Dirichlet perturbation. After the microbial loads are generated, the observed sequencing counts are generated based on the model in Equation 1, where the measurement efficiency of each data set and sample-specific bias of each sample are generated randomly. Due to the above design, the synthetic data sets can capture the unique characteristics of the real microbiome data sets well. In different numerical experiments, the detailed data generation choices may differ slightly. See the details of the numerical experiment setup in Supplementary Material.

In methods comparison, other existing data integration methods are used as recommended in the original paper or package tutorial:

1. ConQuR: call the `ConQuR` function in R package `ConQuR`, and observed sequencing counts are used as input.
2. ComBatSeq: call the `ComBat_seq` function in R package `sva`, and observed sequencing counts are used as input.
3. MMUPHin: call the `adjust batch` function in R package `MMUPHin`, and observed sequencing counts are used as input.
4. PLSDA-batch: call the `PLSDA_batch` function in R package `PLSDAbatch`, and centered log ratio transformed data is used as input. For imbalanced data sets, we set the parameter `balance` to be `FALSE`.
5. Percentile Normalization: call the `percentilenorm` function in R package `PLSDAbatch`, and relative abundance is used as input.
6. DEBIAS-M: call the `debiasm` python package, and observed sequencing counts are used as input.
7. scANVI: call the `scvi-tools` python package, and observed sequencing counts are used as input.

All the methods are run with the default parameters. In experiments involving unobserved confounding variables, we generate an uninformative binary variable following a Bernoulli distribution with *p* = 0.5 and use it as the outcome variable for all methods.

### 4.7 Implementation of Machine Learning Algorithms

In numerical experiments of Section 2.5, random forest and *k*-NN classifiers are applied to corrected counts by different data integrative methods. Random forest algorithm is implemented via R package `caret` (Kuhn, 2008). We choose the number of trees in the random forest algorithm as 500. Model parameters are selected through five-fold cross-validation under the ROC-AUC metric. The *k*-NN algorithm is implemented using the R package `caret`, with tuning over five different values of *k* and 5-fold cross-validation. Each experiment is repeated 500 times, and the average ROC-AUC score on the test data set is compared across integrative methods. In the meta-analysis of the CRC data set presented in Section 2.6, the random forest algorithm is implemented in the same manner described in Section 2.5 and the same data split is used for cross-validation during the training process. In the meta-analysis of responses to PD-1 therapy in Section 2.7, the random forest is trained in the same way as in Section 2.6. The two-layer neural network classifier is implemented using `pytorch`. In the first fully connected layer, the features are reduced from 881 dimensions to 64 dimensions with the ReLU activation function. The second layer is also a fully connected layer with a sigmoid function. Cross-entropy loss is used in classifier training.

## Acknowledgments

The authors acknowledge support from National Science Foundation Grants.

## Data Availability

Data set in He et al. (2018) can be downloaded from Qiita (https://qiita.ucsd.edu/) under study ID 11757. Data sets in (Duvallet et al., 2017) can be found in Zenodo under the identifier No. 840333. In Section 2.6, data sets in Wirbel et al. (2019) can be found in Zenodo (https://zenodo.org/) under the identifier No. 3517209. In Section 2.7, the data set in Chaput et al. (2017) can be retrieved with accession number PRJNA379764; the data set in Frankel et al. (2017) can be retrieved with accession number PRJNA397906; the data set in Matson et al. (2018) can be retrieved with accession number PRJNA399742; and the data set in Peters et al. (2019) can be retrieved with accession number PRJNA541981; the data set in Gopalakrishnan et al. (2018) can be retrieved with accession number PRJEB22894, PRJEB22874, PRJEB22895, PRJEB22893.

## Code Availability

The R package is available at https://github.com/BoYuan07/MetaDICT. All analyses can be found under https://github.com/BoYuan07/MetaDICT_manuscript_code.

## Author Contributions

B. Y. and S. W. conceptualized the problem, developed the methodology, and prepared the manuscript. B. Y. performed the numerical studies, visualized the results, and developed software. S. W. supervised the work.

## References

M. Amodio, D. Van Dijk, K. Srinivasan, W. S. Chen, H. Mohsen, K. R. Moon, A. Campbell, Y. Zhao, X. Wang, M. Venkataswamy, A. Desai, V. Ravi, P. Kumar, R. Montgomery, G. Wolf, and S. Krishnaswamy. Exploring single-cell data with deep multitasking neural networks. Nature Methods, 16(11):1139–1145, 2019.

G. I. Austin, A. B. Kav, H. Park, J. Biermann, A.-C. Uhlemann, and T. Korem. Processingbias correction with debias-m improves cross-study generalization of microbiome-based prediction models. bioRxiv, 2024.

K. Bao, N. Bostanci, T. Thurnheer, and G. N. Belibasakis. Proteomic shifts in multi-species oral biofilms caused by anaeroglobus geminatus. Scientific Reports, 7(1):4409, 2017.

N. Barkas, V. Petukhov, D. Nikolaeva, Y. Lozinsky, S. Demharter, K. Khodosevich, and P. V. Kharchenko. Joint analysis of heterogeneous single-cell rna-seq dataset collections. Nature Methods, 16(8):695–698, 2019.

A. Basic, M. Blomqvist, G. Dahlén, and G. Svensäter. The proteins of fusobacterium spp. involved in hydrogen sulfide production from l-cysteine. BMC Microbiology, 17:1–10, 2017.

Y. Benjamini and T. P. Speed. Summarizing and correcting the gc content bias in highthroughput sequencing. Nucleic Acids Research, 40(10):e72–e72, 2012.

V. D. Blondel, J. Guillaume, R. Lambiotte, and E. Lefebvre. Fast unfolding of communities in large networks. Journal of Statistical Mechanics: Theory and Experiment, 2008(10):P10008, 2008.

M. Bredon, K. le Malicot, C. Louvet, Lu. Evesque, D. Gonzalez, D. Tougeron, and H. Sokol. Faecalibacterium prausnitzii is associated with clinical response to immune checkpoint inhibitors in patients with advanced gastric adenocarcinoma: Results of microbiota analysis of prodige 59-ffcd 1707-durigast trial. Gastroenterology, 2024.

J. H. Bullard, E. Purdom, K. D. Hansen, and S. Dudoit. Evaluation of statistical methods for normalization and differential expression in mrna-seq experiments. BMC Bioinformatics, 11(1):1–13, 2010.

A. Butler, P. Hoffman, P. Smibert, E. Papalexi, and R. Satija. Integrating single-cell transcriptomic data across different conditions, technologies, and species. Nature Biotechnology, 36(5):411–420, 2018.

R. H. Byrd, P. Lu, J. Nocedal, and C. Zhu. A limited memory algorithm for bound constrained optimization. SIAM Journal on Scientific Computing, 16(5):1190–1208, 1995.

Y. Cao, A. Zhang, and H. Li. Multisample estimation of bacterial composition matrices in metagenomics data. Biometrika, 107(1):75–92, 2020.

C. Carrigg, O. Rice, S. Kavanagh, G. Collins, and V. O’Flaherty. Dna extraction method affects microbial community profiles from soils and sediment. Applied Microbiology and Biotechnology, 77:955–964, 2007.

S. Chaffron, H. Rehrauer, J. Pernthaler, and C. Von Mering. A global network of coexisting microbes from environmental and whole-genome sequence data. Genome Research, 20(7):947–959, 2010.

K. C. G. Chan, S. C. P. Yam, and Z. Zhang. Globally efficient non-parametric inference of average treatment effects by empirical balancing calibration weighting. Journal of the Royal Statistical Society: Series B (Statistical Methodology), 78(3):673–700, 2016.

N. Chaput, P. Lepage, C. Coutzac, E. Soularue, K. Le Roux, C. Monot, L. Boselli, E. Routier, L. Cassard, M. Collins, T. Vaysse, L. Marthey, A. Eggermont, V. Asvatourian, E. Lanoy, C. Mateus, C. Robert, and F. Carbonnel. Baseline gut microbiota predicts clinical response and colitis in metastatic melanoma patients treated with ipilimumab. Annals of Oncology, 28(6):1368–1379, 2017.

Y. Chi, Y. M. Lu, and Y. Chen. Nonconvex optimization meets low-rank matrix factorization: An overview. IEEE Transactions on Signal Processing, 67(20):5239–5269, 2019.

A. Conesa, P. Madrigal, S. Tarazona, D. Gomez-Cabrero, A. Cervera, A. McPherson, M. W. Szczésniak, D. J. Gaffney, L. L. Elo, X. Zhang, and A. Mortazavi. A survey of best practices for rna-seq data analysis. Genome Biology, 17(1):1–19, 2016.

E. R. DeLong, D. M. DeLong, and D. L. Clarke-Pearson. Comparing the areas under two or more correlated receiver operating characteristic curves: a nonparametric approach. Biometrics, pages 837–845, 1988.

C. Duvallet, S. M. Gibbons, T. Gurry, R. A. Irizarry, and E. J. Alm. Meta-analysis of gut microbiome studies identifies disease-specific and shared responses. Nature Communications, 8(1):1784, 2017.

M. Elad and M. Aharon. Image denoising via sparse and redundant representations over learned dictionaries. IEEE Transactions on Image Processing, 15(12):3736–3745, 2006.

K. Faust, J. F. Sathirapongsasuti, J. Izard, N. Segata, D. Gevers, J. Raes, and C. Huttenhower. Microbial co-occurrence relationships in the human microbiome. PLoS Computational Biology, 8(7):e1002606, 2012.

B. Flemer, R. D. Warren, M. P. Barrett, K. Cisek, A. Das, I. B. Jeffery, E. Hurley, O. Micheal, F.s Shanahan, and W. Paul. The oral microbiota in colorectal cancer is distinctive and predictive. Gut, 67(8):1454–1463, 2018.

R. Formiga and H. Sokol. Faecalibacterium prausnitzii: one species with multiple potential implications in cancer research. Gut, 2024.

A. E. Frankel, L. A. Coughlin, J. Kim, T. W. Froehlich, Y. Xie, E. P. Frenkel, and A. Y. Koh. Metagenomic shotgun sequencing and unbiased metabolomic profiling identify specific human gut microbiota and metabolites associated with immune checkpoint therapy efficacy in melanoma patients. Neoplasia, 19(10):848–855, 2017.

E. A. Franzosa, A. Sirota-Madi, J. Avila-Pacheco, N. Fornelos, H. J. Haiser, S. Reinker, T. Vatanen, A. B. Hall, H. Mallick, L. J. McIver, J. S. Sauk, R. G. Wilson, B. W. Stevens, J. M. Scott, K. Pierce, A. A. Deik, K. Bullock, F. Imhann, J. A. Porter, A. Zhernakova, J. Fu, R. K. Weersma, C. Wijmenga, C. B. Clish, H. Vlamakis, C. Huttenhower, and R. J. Xavier. Gut microbiome structure and metabolic activity in inflammatory bowel disease. Nature Microbiology, 4(2):293–305, 2019.

Y. Gao, P. Xu, D. Sun, Y. Jiang, X. Lin, T. Han, J. Yu, C. Sheng, H. Chen, J. Hong, Y. Chen, X. Xiao, and J. Fang. Faecalibacterium prausnitzii abrogates intestinal toxicity and promotes tumor immunity to increase the efficacy of dual ctla4 and pd-1 checkpoint blockade. Cancer Research, 83(22):3710–3725, 2023.

S. M. Gibbons, C. Duvallet, and E. J. Alm. Correcting for batch effects in case-control microbiome studies. PLoS Computational Biology, 14(4):e1006102, 2018.

V. Gopalakrishnan, N. C. Spencer, L. Nezi, A. Reuben, C. M. andrews, V. T. Karpinets, A. P. Prieto, D. Vicente, K. Hoffman, C. S. Wei, P. A. Cogdill, L. Zhao, W. C. Hudgens, S. D. Hutchinson, T. Manzo, M. Petaccia de Macedo, T. Cotechini, T. Kumar, S. W. Chen, M. S. Reddy, R. Szczepaniak Sloane, J. Galloway-Pena, H. Jiang, L. P. Chen, J. E. Shpall, K. Rezvani, M. A. Alousi, F. R. Chemaly, S. Shelburne, M. L. Vence, C. P. Okhuysen, B. V. Jensen, G. A. Swennes, F. McAllister, M. E. Riquelme Sanchez, Y. Zhang, E. Le Chatelier, L. Zitvogel, N. Pons, L. J. Austin-Breneman, E. L. Haydu, M. E. Burton, M. J. Gardner, E. Sirmans, J. Hu, J. A. Lazar, T. Tsujikawa, A. Diab, H. Tawbi, C. I. Glitza, J. W. Hwu, P. S. Patel, E. S. Woodman, N. R. Amaria, A. M. Davies, E. J. Gershenwald, P. Hwu, E. J. Lee, J. Zhang, M. L. Coussens, A. Z. Cooper, A. P. Futreal, R. C. Daniel, J. N. Ajami, F. J. Petrosino, T. M. Tetzlaff, P. Sharma, P. J. Allison, R. R. Jenq, and A. J. Wargo. Gut microbiome modulates response to anti–pd-1 immunotherapy in melanoma patients. Science, 359(6371):97–103, 2018.

S. Gunasekar, J. Lee, D. Soudry, and N. Srebro. Characterizing implicit bias in terms of optimization geometry. In International Conference on Machine Learning, pages 1832– 1841, 2018.

L. Haghverdi, A. T. Lun, M. D. Morgan, and J. C. Marioni. Batch effects in single-cell rna-sequencing data are corrected by matching mutual nearest neighbors. Nature Biotechnology, 36(5):421–427, 2018.

J. Hainmueller. Entropy balancing for causal effects: A multivariate reweighting method to produce balanced samples in observational studies. Political Analysis, pages 25–46, 2012.

K. Hampelska, M. M. Jaworska, Z. Babalska, and T.M. Karpiński. The role of oral microbiota in intra-oral halitosis. Journal of Clinical Medicine, 9(8):2484, 2020.

Y. Hao, T. Stuart, M. H. Kowalski, S. Choudhary, P. Hoffman, A. Hartman, A. Srivastava, G. Molla, S. Madad, C. Fernandez-Granda, and R. Satija. Dictionary learning for integrative, multimodal and scalable single-cell analysis. Nature Biotechnology, 42(2):293–304, 2024.

O. Harismendy, P. C. Ng, R. L. Strausberg, X. Wang, T. B. Stockwell, K. Y. Beeson, N. J. Schork, S. S. Murray, E. J. Topol, S. Levy, and K. A. Frazer. Evaluation of next generation sequencing platforms for population targeted sequencing studies. Genome Biology, 10:1– 13, 2009.

Y. He, W. Wu, H. Zheng, P. Li, D. McDonald, H. Sheng, M. Chen, Z. Chen, G. Ji, Z. Zheng, P. Mujagond, X. Chen, Z. Rong, P. Chen, L. Lyu, X. Wang, C. Wu, N. Yu, Y. Xu, J. Yin, J. Raes, R. Knight, W. Ma, and H. Zhou. Regional variation limits applications of healthy gut microbiome reference ranges and disease models. Nature Medicine, 24(10):1532–1535, 2018.

B. Hie, B. Bryson, and B. Berger. Efficient integration of heterogeneous single-cell transcriptomes using scanorama. Nature Biotechnology, 37(6):685–691, 2019.

K. Hirano and G. W. Imbens. Estimation of causal effects using propensity score weighting: An application to data on right heart catheterization. Health Services and Outcomes Research Methodology, 2(3-4):259–278, 2001.

K. Imai and M. Ratkovic. Covariate balancing propensity score. Journal of the Royal Statistical Society: Series B (Statistical Methodology), 76(1):243–263, 2014.

G. W. Imbens and D. B. Rubin. Causal inference in statistics, social, and biomedical sciences. Cambridge University Press, 2015.

H. F. Kaiser. The varimax criterion for analytic rotation in factor analysis. Psychometrika, 23(3):187–200, 1958.

M. M. Karim, T. Hisamoto, T. Matsunaga, Y. Asahi, Y. Noiri, S. Ebisu, A. Kato, and H. Azakami. Luxs affects biofilm maturation and detachment of the periodontopathogenic bacterium eikenella corrodens. Journal of Bioscience and Bioengineering, 116(3):313–318, 2013.

A. Kim, S. Sevanto, E. R. Moore, and N. Lubbers. Latent dirichlet allocation modeling of environmental microbiomes. PLoS Computational Biology, 19(6):e1011075, 2023.

I. Korsunsky, N. Millard, J. Fan, K. Slowikowski, F. Zhang, K. Wei, Y. Baglaenko, M. Brenner, P. Loh, and S. Raychaudhuri. Fast, sensitive and accurate integration of single-cell data with harmony. Nature Methods, 16(12):1289–1296, 2019.

M. Krsek and E. Wellington. Comparison of different methods for the isolation and purification of total community dna from soil. Journal of Microbiological Methods, 39(1):1–16, 1999.

M. Kuhn. Building predictive models in r using the caret package. Journal of Statistical Software, 28:1–26, 2008.

E. S. Lander. Array of hope. Nature Genetics, 21(1):3–4, 1999.

A. Langdon, N. Crook, and G. Dantas. The effects of antibiotics on the microbiome throughout development and alternative approaches for therapeutic modulation. Genome Medicine, 8:1–16, 2016.

H. Lin and S. D. Peddada. Analysis of compositions of microbiomes with bias correction. Nature Communications, 11(1):3514, 2020.

H. Lin, Y. Yu, L. Zhu, N. Lai, L. Zhang, Y. Guo, X. Lin, D. Yang, N. Ren, Z. Zhu, and Q. Dong. Implications of hydrogen sulfide in colorectal cancer: Mechanistic insights and diagnostic and therapeutic strategies. Redox Biology, 59:102601, 2023.

W. Ling, J. Lu, N. Zhao, A. Lulla, A. M. Plantinga, W. Fu, A. Zhang, H. Liu, H. Song, Z. Li, J. Chen, T. W. Randolph, W. A. Koay, J. R. White, L. J. Launer, A. A. Fodor, K. A. Meyer, and M. C. Wu. Batch effects removal for microbiome data via conditional quantile regression. Nature Communications, 13(1):5418, 2022.

M. Lopez-Siles, S. H. Duncan, L. J. Garcia-Gil, and M. Martinez-Medina. Faecalibacterium prausnitzii: from microbiology to diagnostics and prognostics. The ISME Journal, 11(4):841–852, 2017.

R. Lotte, L. Lotte, N. Degand, A. Gaudart, S. Gabriel, M. Ben H’dech, M. Blois, J. Rinaldi, and R. Ruimy. Infectious endocarditis caused by helcococcus kunzii in a vascular patient: a case report and literature review. BMC Infectious Diseases, 15:1–7, 2015.

M. I. Love, W. Huber, and S. anders. Moderated estimation of fold change and dispersion for rna-seq data with deseq2. Genome Biology, 15(12):1–21, 2014.

C. Lozupone, M. Hamady, S. T. Kelley, and R. Knight. Quantitative and qualitative β diversity measures lead to different insights into factors that structure microbial communities. Applied and Environmental Microbiology, 73(5):1576–1585, 2007.

Q. Luo, P. Zhou, S. Chang, Z. Huang, and X. Zeng. Characterization of butyrate-metabolism in colorectal cancer to guide clinical treatment. Scientific Reports, 13(1):5106, 2023.

S. Ma, B. Ren, H. Mallick, Y. S. Moon, E. Schwager, S. Maharjan, T. L. Tickle, Y. Lu,R. N. Carmody, E. A. Franzosa, L. Janson, and C. Huttenhower. A statistical model for describing and simulating microbial community profiles. PLoS Computational Biology, 17 (9):e1008913, 2021.

S. Ma, D. Shungin, H. Mallick, M. Schirmer, L. H. Nguyen, R. Kolde, E. Franzosa, H. Vlamakis, R. Xavier, and C. Huttenhower. Population structure discovery in meta-analyzed microbial communities and inflammatory bowel disease using mmuphin. Genome Biology, 23(1):208, 2022.

H. Mallick, A. Rahnavard, L. J. McIver, S. Ma, Y. Zhang, L. H. Nguyen, T. L. Tickle, G. Weingart, B. Ren, E. H. Schwager, S. Chatterjee, K. N. Thompson, J. E. Wilkinson, A. Subramanian, Y. Lu, L. Waldron, J. N. Paulson, E. A. Franzosa, H. C. Bravo, and C. Huttenhower. Multivariable association discovery in population-scale meta-omics studies. PLoS Computational Biology, 17(11):e1009442, 2021.

V. Matson, J. Fessler, R. Bao, T. Chongsuwat, Y. Zha, M.-L. Alegre, J. J. Luke, and T. F. Gajewski. The commensal microbiome is associated with anti–pd-1 efficacy in metastatic melanoma patients. Science, 359(6371):104–108, 2018.

D. Mazurel, M. Carda-Diéguez, T. Langenburg, M. Žiemytė, W. Johnston, C. P. Martínez, F. Albalat, C. Llena, N. Al-Hebshi, S. Culshaw, A. Mira, and B. T. Rosier. Nitrate and a nitrate-reducing rothia aeria strain as potential prebiotic or synbiotic treatments for periodontitis. npj Biofilms and Microbiomes, 9(1):40, 2023.

M. R. McLaren, A. D. Willis, and B. J. Callahan. Consistent and correctable bias in metagenomic sequencing experiments. Elife, 8:e46923, 2019.

M. Medvecky, D. Cejkova, O. Polansky, D. Karasova, T. Kubasova, A. Cizek, and I. Rychlik. Whole genome sequencing and function prediction of 133 gut anaerobes isolated from chicken caecum in pure cultures. BMC Genomics, 19:1–15, 2018.

S. Mo, H. Ru, M. Huang, L. Cheng, X. Mo, and L. Yan. Oral-intestinal microbiota in colorectal cancer: inflammation and immunosuppression. Journal of Inflammation Research, pages 747–759, 2022.

J. L. Morgan, A. E. Darling, and J. A. Eisen. Metagenomic sequencing of an in vitrosimulated microbial community. PloS One, 5(4):e10209, 2010.

E. C. Murphy and I. Frick. Gram-positive anaerobic cocci–commensals and opportunistic pathogens. FEMS Microbiology Reviews, 37(4):520–553, 2013.

C. Noecker, J. Sanchez, J. E. Bisanz, V. Escalante, M. Alexander, K. Trepka, A. Heinken, Y. Liu, D.n Dodd, I. Thiele, B. C. DeFelice, and P. J. Turnbaugh. Systems biology elucidates the distinctive metabolic niche filled by the human gut microbe eggerthella lenta. PLoS Biology, 21(5):e3002125, 2023.

J. Pan, J. J. Kwon, J. A. Talamas, A. A. Borah, F. Vazquez, J. S. Boehm, A. Tsherniak, M. Zitnik, J. M. McFarland, and W. C. Hahn. Sparse dictionary learning recovers pleiotropy from human cell fitness screens. Cell Systems, 13(4):286–303, 2022.

J. N. Paulson, O. C. Stine, H. C. Bravo, and M. Pop. Differential abundance analysis for microbial marker-gene surveys. Nature Methods, 10(12):1200–1202, 2013.

M. Peng, Y. Li, B. Wamsley, Y. Wei, and K. Roeder. Integration and transfer learning of single-cell transcriptomes via cfit. Proceedings of the National Academy of Sciences, 118 (10):e2024383118, 2021.

Z. Peng, S. Cheng, Y. Kou, Z. Wang, R. Jin, H. Hu, X. Zhang, J. Gong, J. Li, M. Lu, X. Wang, J. Zhou, Z. Lu, Q. Zhang, D. Tzeng, D. Bi, Y. Tan, and L. Shen. The gut microbiome is associated with clinical response to anti–pd-1/pd-l1 immunotherapy in gastrointestinal cancer. Cancer Immunology Research, 8(10):1251–1261, 2020.

B. A. Peters, M. Wilson, U. Moran, A. Pavlick, A. Izsak, T. Wechter, J. S. Weber, I. Osman, and J. Ahn. Relating the gut metagenome and metatranscriptome to immunotherapy responses in melanoma patients. Genome Medicine, 11:1–14, 2019.

M. F. Polz and C. M. Cavanaugh. Bias in template-to-product ratios in multitemplate pcr. Applied and Environmental Microbiology, 64(10):3724–3730, 1998.

P. Pons and M. Latapy. Computing communities in large networks using random walks. In Computer and Information Sciences-ISCIS 2005: 20th International Symposium, Istanbul, Turkey, October 26-28, 2005. Proceedings 20, pages 284–293, 2005.

J. M. Robins, M. A. Hernan, and B. Brumback. Marginal structural models and causal inference in epidemiology. Epidemiology, 11(5):550–560, 2000.

M. D. Robinson and A. Oshlack. A scaling normalization method for differential expression analysis of rna-seq data. Genome Biology, 11:1–9, 2010.

P. R. Rosenbaum and D. B. Rubin. The central role of the propensity score in observational studies for causal effects. Biometrika, 70(1):41–55, 1983.

O. Roy and M. Vetterli. The effective rank: A measure of effective dimensionality. In 2007 15th European Signal Processing Conference, pages 606–610. IEEE, 2007.

N. A. Salad Sabrie, S. Halani, F. Maguire, P. Aftanas, R. Kozak, and N. andany. Lachnoanaerobaculum orale bacteremia in a patient with acute myeloid leukemia and stomatitis: An emerging pathogen. IDCases, 33:e01837, 2023.

V. Sadhanala, Y. Wang, and R. J. Tibshirani. Total variation classes beyond 1d: Minimax rates, and the limitations of linear smoothers. Advances in Neural Information Processing Systems, 29, 2016.

K. Sankaran and S. P. Holmes. Latent variable modeling for the microbiome. Biostatistics, 20(4):599–614, 2019.

J. R. Schwebke and L. F. Lawing. Prevalence of mobiluncus spp among women with and without bacterial vaginosis as detected by polymerase chain reaction. Sexually Transmitted Diseases, 28(4):195–199, 2001.

Y. F. Shaikh, R. J. White, J. J. Gills, T. Hakozaki, C. Richard, B. Routy, Y. Okuma, M. Usyk, A. Pandey, S. J. Weber, J. Ahn, J. E. Lipson, J. Naidoo, M. D. Pardoll, and L. C. Sears. A uniform computational approach improved on existing pipelines to reveal microbiome biomarkers of nonresponse to immune checkpoint inhibitors. Clinical Cancer Research, 27(9):2571–2583, 2021.

N. Srebro, J. Rennie, and T. Jaakkola. Maximum-margin matrix factorization. Advances in Neural Information Processing Systems, 17, 2004.

D. Ternes, J. Karta, M. Tsenkova, P. Wilmes, S. Haan, and E. Letellier. Microbiome in colorectal cancer: how to get from meta-omics to mechanism? Trends in Microbiology, 28 (5):401–423, 2020.

H. Tilg, T. E. Adolph, R. R. Gerner, and A. R. Moschen. The intestinal microbiota in colorectal cancer. Cancer Cell, 33(6):954–964, 2018.

P. J. Turnbaugh, M. Hamady, T. Yatsunenko, B. L. Cantarel, A. Duncan, R. E. Ley, M. L. Sogin, W. J. Jones, B. A. Roe, J. P. Affourtit, M. Egholm, B. Henrissat, A. C. Heath, R. Knight, and J. I. Gordon. A core gut microbiome in obese and lean twins. Nature, 457 (7228):480–484, 2009.

D. Vandeputte, G. Kathagen, K. D’hoe, S. Vieira-Silva, M. Valles-Colomer, J. Sabino, J. Wang, R. Y. Tito, L. De Commer, Y. Darzi, S. Vermeire, G. Falony, and J. Raes. Quantitative microbiome profiling links gut community variation to microbial load. Nature, 551(7681):507–511, 2017.

S. Wang. Multiscale adaptive differential abundance analysis in microbial compositional data. Bioinformatics, 39(4):btad178, 2023a.

S. Wang. Robust differential abundance test in compositional data. Biometrika, 110(1):169–185, 2023b.

Y. Wang and K. Lê Cao. Plsda-batch: a multivariate framework to correct for batch effects in microbiome data. Briefings in Bioinformatics, 24(2):bbac622, 2023.

J. D. Welch, V. Kozareva, A. Ferreira, C. Vanderburg, C. Martin, and E. Z. Macosko. Singlecell multi-omic integration compares and contrasts features of brain cell identity. Cell, 177 (7):1873–1887, 2019.

S. Wie. Clinical significance of providencia bacteremia or bacteriuria. The Korean Journal of Internal Medicine, 30(2):167, 2015.

J. Wirbel, P. T. Pyl, E. Kartal, K. Zych, A. Kashani, A. Milanese, J. S. Fleck, A. Y. Voigt, A. Palleja, R. Ponnudurai, S. Sunagawa, L. P. Coelho, P. Schrotz-King, E. Vogtmann, N. Habermann, E. Niméus, A. M. Thomas, P. Manghi, S. Gandini, D. Serrano, S. Mizutani, H. Shiroma, S. Shiba, T. Shibata, S. Yachida, T. Yamada, L. Waldron, A. Naccarati, N. Segata, R. Sinha, C. M. Ulrich, H. Brenner, M. Arumugam, P. Bork, and G. Zeller. Meta-analysis of fecal metagenomes reveals global microbial signatures that are specific for colorectal cancer. Nature Medicine, 25(4):679–689, 2019.

T. Woyke, H. Teeling, N. N. Ivanova, M. Huntemann, M. Richter, F. O. Gloeckner, D. Boffelli, I. J. anderson, K. W. Barry, H. J. Shapiro, E. Szeto, N. C. Kyrpides, M. Mussmann, R. Amann, C. Bergin, C. Ruehland, E. M. Rubin, and N. Dubilier. Symbiosis insights through metagenomic analysis of a microbial consortium. Nature, 443(7114):950–955, 2006.

J. Wright, A. Y. Yang, A. Ganesh, S. S. Sastry, and Y. Ma. Robust face recognition via sparse representation. IEEE Transactions on Pattern Analysis and Machine Intelligence, 31(2):210–227, 2008.

J. Wu, S.n Wang, B. Zheng, X. Qiu, H. Wang, and L. Chen. Modulation of gut microbiota to enhance effect of checkpoint inhibitor immunotherapy. Frontiers in Immunology, 12:669150, 2021a.

Y. Wu, N. Jiao, R. Zhu, Y. Zhang, D. Wu, A. Wang, S. Fang, L. Tao, Y. Li, S. Cheng, X. He, P. Lan, C. Tian, N. Liu, and L. Zhu. Identification of microbial markers across populations in early detection of colorectal cancer. Nature Communications, 12(1):3063, 2021b.

C. Xu, R. Lopez, E. Mehlman, J. Regier, M. I. Jordan, and N. Yosef. Probabilistic harmonization and annotation of single-cell transcriptomics data with deep generative models. Molecular Systems Biology, 17(1):e9620, 2021.

Y. Yang, L. Du, D. Shi, C. Kong, J. Liu, G. Liu, X. Li, and Y. Ma. Dysbiosis of human gut microbiome in young-onset colorectal cancer. Nature Communications, 12(1):6757, 2021.

T. Yatsunenko, F. E. Rey, M. J. Manary, I. Trehan, M. G. Dominguez-Bello, M. Contreras, M. Magris, G. Hidalgo, R. N. Baldassano, A. P. Anokhin, A. C. Heath, B. Warner, J. Reeder, J. Kuczynski, J. G. Caporaso, C. A. Lozupone, C. Lauber, J. C. Clemente, D. Knights, R. Knight, and J. I. Gordon. Human gut microbiome viewed across age and geography. Nature, 486(7402):222–227, 2012.

H. Ye, X. Zhang, C. Wang, E. L. Goode, and J. Chen. Batch-effect correction with sample remeasurement in highly confounded case-control studies. Nature Computational Science, 3(8):709–719, 2023.

R. Yu and S. Wang. Treatment effects estimation by uniform transformer. In The Twelfth International Conference on Learning Representations, 2024.

B. Yuan and S. Wang. Rsim: A reference-based normalization method via rank similarity. PLoS Computational Biology, 19(9):e1011447, 2023.

M. Zepeda-Rivera, S. S. Minot, H. Bouzek, H. Wu, A. Blanco-Míguez, P. Manghi, D. S. Jones, K. D. LaCourse, Y. Wu, E. F. McMahon, S. Park, Y. K. Lim, A. G. Kempchinsky, A. D. Willis, S. L. Cotton, S. C. Yost, E. Sicinska, J. Kook, F. E. Dewhirst, N. Segata, S. Bullman, and C. D. Johnston. A distinct fusobacterium nucleatum clade dominates the colorectal cancer niche. Nature, 628(8007):424–432, 2024.

Y. Zhang, G. Parmigiani, and W. E. Johnson. Combat-seq: batch effect adjustment for rna-seq count data. NAR Genomics and Bioinformatics, 2(3):qaa078, 2020.

Z. Zhang, H. Tang, P. Chen, H. Xie, and Y. Tao. Demystifying the manipulation of host immunity, metabolism, and extraintestinal tumors by the gut microbiome. Signal Transduction and Targeted Therapy, 4(1):41, 2019.

H. Zhou, K. He, J. Chen, and X. Zhang. Linda: linear models for differential abundance analysis of microbiome compositional data. Genome Biology, 23(1):95, 2022.

J. R. Zubizarreta. Stable weights that balance covariates for estimation with incomplete outcome data. Journal of the American Statistical Association, 110(511):910–922, 2015.

F. H. Zwezerijnen-Jiwa, H. Sivov, P. Paizs, K. Zafeiropoulou, and J. Kinross. A systematic review of microbiome-derived biomarkers for early colorectal cancer detection. Neoplasia, 36:100868, 2023.

