## Supplemental Materials for "Microbiome Data Integration via Shared Dictionary Learning"

### Supplementary Material to “Microbiome Data Integration via Shared Dictionary Learning”

Bo Yuan* and Shulei Wang*
University of Illinois at Urbana-Champaign

(June 12, 2025)

### Numerical Experiment Setup

All simulated datasets are generated using a microbiome dataset collected in He et al. (2018). Count vectors aggregated at the order level from samples of individuals under the age of 30 are used as the basis. In total, the dataset consists 539 samples and 131 taxa.

**Summary of Notations** We denote the number of datasets by $m$, sample size per dataset by $n$, and the number of taxa by $d$. $N_{i,j,k}$ represents the sequencing count from He et al. (2018). $A_{i,j,k}$ denotes the simulated microbial loads, and $O_{i,j,k}$ denotes the simulated observed abundance. Measurement efficiency is denoted by $w_{i,k}$, and sample-specific bias is represented by $c_{i,j}$. Sample covariate is denoted by $X_{i,j}$.

#### Setting for Section 2.2

**Figure 2(a) and (b)** We simulate $m$ datasets with known measurement efficiency. First, we randomly draw $m$ subsets of $n$ samples, $N_1, \ldots, N_m$, from the microbiome dataset in He et al. (2018). The microbial loads for dataset $i$, sample $j$ is simulated as follows:

$$\vec{A}_{i,j,\cdot} \sim \text{Dirichlet}(\vec{N}_{i,j,\cdot} + 0.1), \qquad i = 1, \ldots, m, \quad j = 1, \ldots, n.$$

Adding a small value 0.1 ensures that observed zeros are mostly sampling zeros.

Then we generate different measurement efficiencies $\vec{w}_i$ for each dataset. A neighborhood graph $G$ is constructed based on the cophenetic distance among taxa on the phylogenetic tree. The weight on the edge between taxa $k_1$ and $k_2$ is $\exp(-\text{dist}(k_1, k_2)^2/\sigma)$. For dataset $i$, the measurement efficiency vector $\vec{w}_i = [w_{i,1}, \ldots, w_{i,d}]^T$ is defined as a linear combination of the last 10 eigenvectors $\vec{e}_1, \ldots, \vec{e}_{10}$ of the graph Laplacian matrix $L_G$:

$$\vec{w}_i = \sum_{h=1}^{10} u_{i,h} \vec{e}_h$$

where $u_{i,h} \sim \text{Unif}(-1, 1)$ is the coefficient of eigenvector $\vec{e}_h$. The vector $\vec{w}_i$ is rescaled such that the measurement efficiency for each taxon lies within the range $[0, 1]$. The observed abundance is simulated as follows:

$$O_{i,j,k} = w_{i,k} A_{i,j,k} c_{i,j}, \qquad i = 1, \ldots, m, \quad j = 1, \ldots, n, \quad k = 1, \ldots, d$$

where $c_{i,j} \sim \text{Unif}(10000, 15000)$ represents the sample-specific bias for sample $j$ in dataset $i$.

This set of experiment aims to assess whether the shared dictionary learning in MetaDICT can improve the initial estimation of measurement efficiency. Specifically, we compare the estimation accuracy of measurement efficiency with and without the second stage. In (a), the mean absolute error of estimated measurement efficiency for taxon with abundance rank $k$ is defined as

$$\hat{\text{MAE}}(k) = \frac{1}{T} \sum_{t=1}^{T} \frac{1}{m} \sum_{i=1}^{m} |\hat{w}_{i,r_k}^{(t)} - w_{i,r_k}^{(t)}|$$

where $r_k$ represents the taxon with rank $k$ defined by the total absolute abundance across $m$ datasets, $T$ represents the total iterations of simulation experiments. In (b), we calculate the average Pearson correlation between the flattened measurement efficiency vector and ground truth. The experiments are repeated 500 times.

**Figure ??** This set of experiments is aimed at illustrating the effect of $\alpha$ and $\beta$. The basic settings are the same as in Figure 2(a) and (b). Figure (a) compares the measurement efficiency estimation when $\beta = 0$ and $\beta > 0$. Figure (b) shows the average Pearson correlation between the measurement efficiency estimates and the ground truth across 500 iterations when $\beta = 0$ and $\beta > 0$. Figure (c) plots the singular values of the estimated shared dictionary $D$ and compares the trends between $\alpha = 0.1$ and $\alpha = 1$.

**Figure ??** We investigate the robustness of our method in measurement efficiency estimation under various settings. Basic setting is same with Figure 2(a) and (b). In (a), the number of datasets is set to be 3, 5, 7; in (b), the sample size per dataset is set to be 50, 100, 150; in (c), we compare the estimation accuracy between the case when all the datasets have the same sample size with the case when sample size varies across datasets; in (d), we change the smoothness of true measurement efficiency by increasing the number of included eigenvectors from 5 to 15 in measurement efficiency generation. The average Pearson correlation across 500 iterations is reported.

**Figure 2(c), (d) and ??** This experiment compares the performance of MetaDICT and other data integration methods in PCoA analyses. We sample two subsets of 200 samples, $N_1$ and $N_2$, from He et al. (2018) as basis for simulated datasets. Then we randomly generate a binary label $X_{i,j} \sim \text{Bernoulli}(0.5)$ as target covariate of interest. The set of samples with $X_{i,j} = 0$ is denoted as biological group $\mathcal{G}_1$, and the set with $X_{i,j} = 1$ is denoted as biological group $\mathcal{G}_2$. We let the microbial loads vary between $\mathcal{G}_1$ and $\mathcal{G}_2$:

$$\vec{A}'_{i,j,\cdot} \sim \text{Dirichlet}(\vec{N}_{i,j,\cdot} + 0.1), \qquad i = 1, \ldots, m, \quad j = 1, \ldots, n$$

$$A_{i,j,k} = \begin{cases} sA'_{i,j,k}, & k \in \mathcal{J}, j \in \mathcal{G}_1 \\ A'_{i,j,k}, & \text{otherwise} \end{cases},$$

where $\mathcal{J}$ is the set of differentially abundant taxa between $\mathcal{G}_1$ and $\mathcal{G}_2$, and $s = 5$. In the experiment, $\mathcal{J}$ is a randomly selected set of 40 taxa. The observed abundances are generated in the following way:

$$O_{i,j,k} = w_{i,k} A_{i,j,k} c_{i,j}, \qquad i = 1, \ldots, m, \quad j = 1, \ldots, n, \quad k = 1, \ldots, d,$$

where sample-specific bias $c_{i,j} \sim \text{Unif}(10000, 15000)$. We also create an uninformative label $X'_{i,j} \sim \text{Bernoulli}(0.5)$ which is unrelated to absolute abundance. For experiments where biological covariates are observed, both $X_{i,j}$ and $X'_{i,j}$ are used as input in data integration methods; for experiments where biological covariates are not observed, only $X'_{i,j}$ is used as input in data integration methods.

**Figure ??** This experiment compares the performance of MetaDICT and other methods in PCoA analyses when biological variable is continuous. We sample two subsets of 200 samples, $N_1$ and $N_2$, from He et al. (2018) as basis for simulated datasets. A latent variable $X_{i,j} \sim \text{Unif}(1, 10)$ is introduced as a continuous sample covariate. A random set of 15 taxa, denoted as $\mathcal{J}$, is selected. The abundance of these taxa is varied as a function of $X$ to simulate realistic biological changes:

$$
\begin{aligned}
&\vec{A}'_{i,j,\cdot} \sim \text{Dirichlet}(\vec{N}_{i,j,\cdot} + 0.1), \qquad i = 1, \dots, m, \quad j = 1, \dots, n \\
&A_{i,j,k} = \begin{cases} A'_{i,j,k} X_{i,j}, & k \in \mathcal{J} \\ A'_{i,j,k}, & \text{otherwise} \end{cases} \\
&O_{i,j,k} = w_{i,k} A_{i,j,k} c_{i,j}, \qquad i = 1, \dots, m, \quad j = 1, \dots, n, \quad k = 1, \dots, d.
\end{aligned}
$$

where sample-specific bias $c_{i,j} \sim \text{Unif}(10000, 15000)$. We also create an uninformative label $X'_{i,j} \sim \text{Bernoulli}(0.5)$ which is unrelated to microbial compositions. For experiments where biological covariates are observed, both $X_{i,j}$ and $X'_{i,j}$ are used as input in data integration methods; for experiments where biological covariates are not observed, only $X'_{i,j}$ is used as input in data integration methods.

#### Settings for Section 2.3

**Figure 3(a) and (b)** This set of experiments aims to investigate whether integrative methods can avoid overcorrection when biological variables are confounded with batches. We consider an ideal case, where there is no batch effects but distribution shifts in the absolute abundance between two studies. In the experiment, we sample two subsets of 200 samples $N_1$, $N_2$ from He et al. (2018) as basis for simulated datasets. Then we assign each sample a binary label $X_{i,j} \sim \text{Bernoulli}(p_i)$ as indicator of biological conditions. Samples with $X_{i,j} = 0$ are grouped into the biological group $\mathcal{G}_1$, while those with $X_{i,j} = 1$ form the group $\mathcal{G}_2$. We let the microbial loads vary between $\mathcal{G}_1$ and $\mathcal{G}_2$, while the measurement efficiency remains the same between the two studies:

$$
\begin{aligned}
&A'_{i,j,k} = \begin{cases} s(N_{i,j,k}/\sum \vec{N}_{i,j,\cdot} + 0.1), & k \in \mathcal{J}, j \in \mathcal{G}_1 \\ N_{i,j,k}/\sum \vec{N}_{i,j,\cdot}, & \text{otherwise} \end{cases} \\
&\vec{A}_{i,j,\cdot} \sim \text{Dirichlet}(\vec{A}'_{i,j,\cdot}), \\
&O_{i,j,k} = A_{i,j,k} c_{i,j}, \qquad i = 1, \dots, m, \quad j = 1, \dots, n, \quad k = 1, \dots, d.
\end{aligned}
$$

where $c_{i,j} \sim \text{Unif}(1000, 1500)$, $s = 10$, $\mathcal{J}$ is a randomly selected set of 40 taxa that are set to be differentially abundant between $\mathcal{G}_1$ and $\mathcal{G}_2$. Note that there is no variation of batch effects and sequencing depth between batches.

To mimic the case when covariate distribution shift exists between studies, we let the parameter of Bernoulli distribution $p_1 \neq p_2$. Specifically, we let $p_1 = 1/6, p_2 = 5/6$. In this case, samples in $\mathcal{G}_1$ are significantly more than samples in $\mathcal{G}_2$ for study 1, while samples in

$\mathcal{G}_2$ are significantly more than samples in $\mathcal{G}_1$ for study 2. We also create an uninformative label $X'_{i,j} \sim \text{Bernoulli}(0.5)$ which is unrelated to absolute abundance. For experiments where biological covariates are observed, both $X_{i,j}$ and $X'_{i,j}$ are used as input in data integration methods; for experiments where biological covariates are not observed, only $X'_{i,j}$ is used as input in data integration methods.

**Figure ??** The basic setup is the same as in Figure 3(a) and (b). When there is no confounding effect between biological conditions and batches, the Bernoulli distribution coefficients $p_1 = p_2 = 1/2$. For the medium-level confounding effect, we set $p_1 = 1/4$ and $p_2 = 3/4$. In the experiment with high-level confounding effect, $p_1 = 1/6$ and $p_2 = 5/6$. Each experiment is repeated 500 times, and we calculate the PERMANOVA $R^2$ statistic to measure the variation in abundances due to the biological variable $X_{i,j}$ and batch effects. Overcorrection is indicated when the $R^2$ statistics for both batches and biological groups are lower than the ground truth.

**Figure ?? and Figure 3(c)** The basic setup is the same as in Figure 2(c) and (d), except that the samples are unevenly divided into two biological groups, $\mathcal{G}_1$ and $\mathcal{G}_2$. We split samples into different groups based on the value of $X_{i,j} \sim \text{Bernoulli}(p_i)$. For Figure **??**, we set $p_1 = 1/4$, $p_2 = 3/4$, and the sample sequence depth $c_{i,j} \sim \text{Unif}(10000, 11000)$, and $s = 5$. For Figure 3(c), we set $p_1 = 1$, $p_2 = 0$, $c_{i,j} \sim \text{Unif}(1000, 1500)$, and $s = 10$, so the batches are completely confounded with the biological groups.

**Figure ??** We use SparseDOSSA2 (Ma et al., 2021) to generate simulated data. 30 control samples from Zeller et al. (2014) are used to fit the basis model for simulated dataset 1, while 30 control samples from Baxter et al. (2016) are used for simulated dataset 2. Each simulated dataset includes 100 samples. Instead of artificially simulating batch bias, we leverage the natural variation between the two datasets as batch effects. A binary phenotype label is introduced using SparseDOSSA2, affecting 5% of the taxa abundances and prevalences with a log fold change of 5.

**Figure ??** We use SparseDOSSA2 (Ma et al., 2021) to generate simulated data. To mimic a scenario with no batch effects but with distribution shifts in absolute abundance between two studies, we simulate two datasets with 100 samples using a single basis model fitted on the real dataset from Zeller et al. (2014). A binary phenotype label is introduced using SparseDOSSA2, affecting 5% of taxa abundances and prevalences with a log fold change of 5. To simulate covariate distribution shifts between studies, we ensure that samples with phenotype 1 are significantly more prevalent than those with phenotype 2 in dataset 1, while the reverse is true in dataset 2.

#### Setting for Section 2.4

In this section, we consider a more challenging setting, where both sample-specific bias and measurement efficiency are confounded with batches. Meanwhile, we directly use subsets of the microbiome dataset in He et al. (2018) as absolute abundance so that the simulated data maintains a similar zero prevalence to the real data.

**Figure 4** We sample two subsets of 50 samples $N_1, N_2$ from the microbiome dataset in He et al. (2018). Then we randomly split samples in each dataset $i$ into four groups $\mathcal{G}_1, \ldots, \mathcal{G}_4$ with probability $\vec{p}_i \in \mathbb{R}^4$. The sample group label is denoted as $X_{i,j}$. We also randomly split taxa into five groups $\mathcal{J}_1, \ldots, \mathcal{J}_5$. Taxa in $\mathcal{J}_1, \ldots, \mathcal{J}_4$ are differential abundant among sample groups, and taxa in $\mathcal{J}_5$ are not differentially abundant across sample groups. The observed abundance is generated in the following way:

$$A_{i,j,k} = \begin{cases} s(N_{i,j,k}+1), & k \in \mathcal{J}_t, j \in \mathcal{G}_t, t = 1,2,3,4, \\ N_{i,j,k}, & \text{otherwise} \end{cases}$$
$$O_{i,j,k} = w_{i,k} A_{i,j,k} c_{i,j}, \qquad i = 1,\ldots,m, \quad j = 1,\ldots,n, \quad k = 1,\ldots,d,$$

where $s = 50$, $c_{i,j}$ is the sample-specific bias. We let $c_{i,j}$ vary across datasets: $c_{i,j} \sim \text{Unif}(100, 500)$ for dataset 1 and $c_{i,j} \sim \text{Unif}(10, 50)$ for dataset 2. Besides, we create an uninformative label $X'_{i,j} \sim \text{Bernoulli}(0.5)$ which is unrelated to microbial compositions. For experiments where biological covariates are observed, both $X_{i,j}$ and $X'_{i,j}$ are used as input in data integration methods; for experiments where biological covariates are not observed, only $X'_{i,j}$ is used as input in data integration methods.

To identify microbial communities (subpopulations), we first construct a $k$-nearest neighbor graph based on the Euclidean distance between taxa (samples). We then apply Louvain algorithm to detect communities (Blondel et al., 2008). To determine the optimal number of neighbors, we try various values of $k$ within a specified range and select the one that yields the highest average Silhouette score. For MetaDICT, we use shared dictionary $D$ to detect microbial community and representation $R$ to detect subpopulations. We select the first 50 columns of $D$ for taxa community detection, where 50 represents the estimated rank of $D$. For other integrative methods, we use corrected count data to detect both microbial community and subpopulations. To compare the microbial community results between single dataset and integrated data, we detect microbial community on each dataset, and present the result with maximum average Silhouette score. The performance is quantified using the adjusted Rand index (ARI) between estimated clusters and ground truth.

In the experiment where batches are independent of sample groups, we let $\vec{p}_1 = \vec{p}_2 = [1/4, 1/4, 1/4, 1/4]^T$; In the experiment where batches are confounded with sample groups, we let

$$\vec{p}_1 = [1/8, 1/2, 1/8, 1/2]$$
$$\vec{p}_2 = [1/2, 1/8, 1/2, 1/8].$$

The average adjusted Rand index between estimated clusters with ground truth is calculated across 500 iterations.

**Figure ??** The setting is the same as Figure 4 when batches are independent with sample groups.

**Figure ??** We investigate the robustness of our method in microbial community estimation. In (a), we set the number of datasets $m$ to be 2, 5, 7, $n = 50$, $s = 50$; in (b), we set $s = 30, 50, 70$, $m = 5$, $n = 50$; in (c), we set the sample size per dataset to be 25, 50, 100, $m = 5$, $s = 50$. Sample groups are independent with batches. The average adjusted Rand index between estimated microbial communities with ground truth is calculated across 500 iterations.

**Figure ??** The setting is the same with Figure **??**.

#### Setting for Section 2.5

**Figure 5(b) and ??(a)** We sample five subsets of 50 samples $N_1, \dots, N_5$ from He et al. (2018), which are used as basis of simulated datasets. Samples in each dataset $i$ are randomly split into two groups $\mathcal{G}_1$ and $\mathcal{G}_2$ with probability $\vec{p}_i \in \mathbb{R}^2$. We denote the group label as $X_{i,j}$. A set of taxa $\mathcal{J} = \mathcal{J}_1 \cup \mathcal{J}_2$ is selected to be differentially abundant between sample groups $\mathcal{G}_1$, and $\mathcal{G}_2$. Taxa in $\mathcal{J}_1$ are more abundant in $\mathcal{G}_1$ than $\mathcal{G}_2$, while taxa in $\mathcal{J}_2$ are more abundant in $\mathcal{G}_2$ than $\mathcal{G}_1$. The observed abundance is generated in the following way:

$$\begin{aligned} A_{i,j,k} &= \begin{cases} sN_{i,j,k}, & k \in \mathcal{J}_1, j \in \mathcal{G}_1 \\ s^{-1}N_{i,j,k}, & k \in \mathcal{J}_2, j \in \mathcal{G}_1 \\ N_{i,j,k}, & \text{otherwise} \end{cases} \\ O_{i,j,k} &= w_{i,k} A_{i,j,k} c_{i,j}, \qquad i = 1, \dots, m, \quad j = 1, \dots, n, \quad k = 1, \dots, d, \end{aligned}$$

where $s = 5$, $c_{i,j}$ is the sample-specific bias. In the experiment, we selected $\mathcal{J}$ from the top 5% most abundant taxa to ensure detectable difference between groups. We let $c_{i,j}$ vary across datasets: $c_{i,j} \sim \text{Unif}(100, 500)$ when $i = 1, 3, 5$ and $c_{i,j} \sim \text{Unif}(10, 100)$ when $i = 2, 4$.

In the setting that sample groups are independent of batches, we let

$$\vec{p}_1 = \cdots = \vec{p}_5 = [1/2, 1/2]^T.$$

In the setting that sample groups are confounded with batches, we let

$$\begin{aligned} \vec{p}_1 = \vec{p}_3 = \vec{p}_5 &= [1/4, 3/4]^T \\ \vec{p}_2 = \vec{p}_4 &= [3/4, 1/4]^T. \end{aligned}$$

Sample covariate $X_{i,j}$ is used as the input of all the integrative methods. We apply $t$-test, RDB, LinDA, ANCOM-BC and MaAsLin2 to detect differential abundant taxa between $\mathcal{G}_1$ and $\mathcal{G}_2$. We choose significance level $\alpha = 0.1$ and use the Benjamini-Hochberg procedure to adjust $p$-values. The experiment is repeated 500 times. Sensitivity and FDR are reported.

**Figure 5(c)** In this experiment, we simulate the case when there is no differential abundant taxon but a distribution shift in outcome across studies. We sample three subsets of 100 samples $N_1, N_2, N_3$ from the microbiome dataset in He et al. (2018) and generate observed abundance in the following way:

$$A_{i,j,k} = N_{i,j,k}$$
$$O_{i,j,k} = w_{i,k} A_{i,j,k} c_{i,j}, \qquad i = 1, \ldots, m, \quad j = 1, \ldots, n, \quad k = 1, \ldots, d.$$

where $c_{i,j}$ varies across datasets: $c_{i,j} \sim \text{Unif}(100, 500)$ when $i = 1, 3$ and $c_{i,j} \sim \text{Unif}(10, 100)$ when $i = 2$. We randomly generate a label $X'_{i,j} \sim \text{Bernoulli}(p_i)$ for samples in dataset $i$ which is independent of the abundance. As a result, there is no differential abundan taxa between samples with $X'_{i,j} = 0$ and $X'_{i,j} = 1$. To mimic the confounding effect between sample covariate and batches, we let the parameter of Bernoulli distribution vary across datasets. Specifically, we consider three scenarios:

1. No confounding effect between batches and $X'_{i,j}$ ($p_1 = p_2 = p_3 = 0.5$): The proportion between samples with different $X'$ values is roughly the same across datasets.

2. Median-level confounding effects between batches and $X'_{i,j}$ ($p_1 = 0.8, p_2 = 0.2, p_3 = 0.8$): The proportion between samples with different $X'$ values significantly varies across datasets.

3. High-level confounding effects between batches and $X'_{i,j}$ ($p_1 = 0.99, p_2 = 0.01, p_3 = 0.99$): The proportion between samples with different $X'$ values extremely varies across datasets.

Then we detect differentially abundant taxa between groups $X'_{i,j} = 0$ and $X'_{i,j} = 1$ using two-sample $t$-test. We choose significance level $\alpha = 0.1$, and the Benjamini-Hochberg procedure to adjust $p$-values. The experiment is repeated 500 times and FDR is reported.

**Figure 5(a)** The basic setting is the same with Figure 5(b) when the sample groups are confounded with batches. We generate two datasets and quantify the disturbance level using standardized mean difference of individual taxa abundance between two datasets. A histogram displaying the frequency distribution of disturbance levels in false discoveries is presented.

**Figure ??(b)** We integrate healthy samples from three studies Zeller et al. (2014); Zackular et al. (2014); Baxter et al. (2016), all sequenced in the same 16S gene region and processed using the same bioinformatics pipeline Gibbons et al. (2018). We create a random label $X'_{i,j} \sim \text{Bernoulli}(p_i)$ which is independent of the count table. Then we use $X'$ as a covariate in all the integrative methods. We consider three cases:

1. No confounding effect between batches and $X'_{i,j}$ ($p_1 = p_2 = p_3 = 0.5$): The proportion between samples with different $X'$ values is roughly the same across datasets.

2. Median-level confounding effects between batches and $X'_{i,j}$ ($p_1 = 0.85, p_2 = 0.5, p_3 = 0.15$): The proportion between samples with different $X'$ values significantly varies across datasets.

3. High-level confounding effects between batches and $X'_{i,j}$ ($p_1 = 0.99, p_2 = 0.5, p_3 = 0.01$): The proportion between samples with different $X'$ values extremely varies across datasets.

Then we detect differentially abundant taxa between groups $X'_{i,j} = 0$ and $X'_{i,j} = 1$ using two-sample $t$-test. We choose significance level $\alpha = 0.1$, and the Benjamini-Hochberg procedure to adjust $p$-values. The experiment is repeated 500 times and FDR is reported.

**Figure 5(d) and Figure ??(a)** We sampled two subsets of 100 samples each, denoted as $N_1$ and $N_2$, from He et al. (2018) to serve as the basis for our simulated datasets. In each dataset $i$, samples were randomly partitioned into two groups, $\mathcal{G}_1$ and $\mathcal{G}_2$, according to a probability vector $\vec{p_i} \in \mathbb{R}^2$. We denote the group label for sample $j$ in dataset $i$ as $X_{i,j}$. The top 5% most abundant taxa, denoted by $\mathcal{J}$, were selected as differentially abundant between the two groups. The observed abundance is generated as follows:

$$A_{i,j,k} = \begin{cases} s(N_{i,j,k}+1), & k \in \mathcal{J}, j \in \mathcal{G}_1 \\ N_{i,j,k}, & \text{otherwise} \end{cases}$$
$$O_{i,j,k} = w_{i,k} A_{i,j,k} c_{i,j}, \qquad i = 1,\ldots,m, \quad j = 1,\ldots,n, \quad k = 1,\ldots,d.$$

where $s = 4$, $c_{i,j}$ is the sample-specific bias. We let $c_{i,j}$ vary across datasets: $c_{i,j} \sim \text{Unif}(1,5)$ when $i = 1$ and $c_{i,j} = 1$ when $i = 2$. Besides, we create an uninformative label $X'_{i,j} \sim \text{Bernoulli}(0.5)$ used as negative control variable. For experiments where biological covariates are observed, both $X_{i,j}$ and $X'_{i,j}$ are used as input in data integration methods; for experiments where biological covariates are not observed, only $X'_{i,j}$ is used as input in data integration methods.

We train the classifier on dataset 1 and test the performance on dataset 2. For the setting where no distribution shift exists across dataset, we let

$$\vec{p_1} = \vec{p_2} = [1/2, 1/2]^T.$$

For the setting where distribution shift exists, we let

$$\vec{p_1} = [1/4, 3/4]^T$$
$$\vec{p_2} = [1/6, 5/6]^T.$$

Each experiment is repeated 500 times, and the average ROC-AUC score on the test dataset is compared across integrative methods.

To statistically compare the ROC-AUC scores, we applied a one-sided DeLong's test to evaluate differences between the ROC-AUC scores of models trained on MetaDICT-processed data and those trained on data integrated using other methods. DeLong's test (DeLong et al., 1988) was implemented using the `roc.test` function from the `pROC` R package (Robin et al., 2011). In each experiment, we compared the ROC-AUC of a classifier trained on MetaDICT-processed data against that of a classifier trained on data processed by an alternative integration method. The null hypothesis is that there is no significant difference between the two AUCs, while the alternative hypothesis states that the AUC of the classifier using MetaDICT-processed data is greater than that using the other method. We set the significance level to 0.1 and repeated each experiment 500 times. Figure **??**(a) shows the proportion of experiments in which the null hypothesis was rejected ($p$-value $\leq 0.1$).

**Figure 5(e) and Figure ??(b)** Data generation process is the same as Figure 5(d) when where no distribution shift exists across dataset. Instead of using one dataset as training set and one as test set, we combine all the samples together and split them into the training and test set with a ratio of 3:1. Similar to Figure **??**(a), one-sided DeLong's test was applied to compare the ROC-AUCs scores of models trained on MetaDICT-processed data against that of a classifier trained on data processed by an alternative integration method.

**Figure 5(f) and Figure ??(c)** Data generation process is the same as Figure 5(e). We use the random forest and $k$-NN to predict uninformative covariate $X'_{i,j}$. We applied DeLong's test to investigate whether the ROC-AUCs of classifiers trained on integrated data is significantly larger than 0.5. To simulate the output of a random-guess classifier, we set predicted probabilities of each label category as 0.5 for each sample. In each experiment, we applied a one-sided DeLong's test. The null hypothesis is that AUC of classifier is not significantly larger than 0.5, while the alternative hypothesis states that the AUC of the classifier trained on integrated data is greater than 0.5. We set the significance level to 0.1 and repeated the experiment 500 times. Figure **??**(c) shows the proportion of experiments in which the null hypothesis was rejected ($p$-value $\leq 0.1$).

#### Setting for Section 2.6

| Study | Region | No. of samples | Phenotype(s) | Reference |
|---|---|---|---|---|
| US | United States | 104 | CRC 52/CTR 52 | Vogtmann et al. (2016) |
| DE | Germany | 120 | CRC 60/CTR 60 | Wirbel et al. (2019) |
| CN | China | 128 | CRC 74/CTR 54 | Yu et al. (2017) |
| AT | Austria | 109 | CRC 46/CTR 63 | Feng et al. (2015) |
| FR | France | 114 | CRC 53/CTR 61 | Zeller et al. (2014) |

Table S1: **Five CRC datasets included in the data integration.**

##### Genus-level Analysis

We integrate five datasets related to colorectal cancer, with basic information presented in Table S1. Taxonomy was assigned using mOTUs2 (Milanese et al., 2019), and taxa are aggregated to the genus level based on taxonomic profiles. Sample covariates included age, gender, BMI, country, and disease status. Samples with missing values in any of these covariates were filtered out. In total, there are 272 taxa and 567 samples. The sample compositions after filtering are shown in Figure **??** (a).

**Implementation of MetaDICT** We selected parameters of MetaDICT as $\alpha = 1$, $\beta = 0.01$, $\gamma = 1$. Since phylogenetic tree is not available, we used taxonomic information to construct taxa similarity graph. In this graph, taxa within the same family were connected with each other.

**Microbial Community Detection.** Microbial communities are detected using the first 50 columns of dictionary $D$ learned by MetaDICT, employing the method described in Section 2.4. Here we select the number of neighbors as 2 and use the Walktrap algorithm to detect microbial community.

**Genus-Level Association Analysis.** LinDA is applied to test the association between genera abundance and disease status, while controlling for the confounding effects of age, gender, BMI, and country. The Benjamini-Hochberg procedure is used to adjust $p$-values, with a significance level of $\alpha = 0.1$.

**Random Forest Prediction.** We consider two types of random forest classifiers. The first type of classifier is trained on each individual study and evaluated on the other four studies. The second type of classifier is trained on all-but-one study and evaluated on the left study. The random forest algorithm is implemented using the R package `caret` (Kuhn, 2008), with the number of decision trees also set to 500. Model parameters are selected through five-fold cross-validation under the ROC-AUC metric. The two-sided DeLong's test was applied to compare the ROC-AUCs of classifier trained on MetaDICT-processed data and classifiers trained on other integrated data. The significance level is selected to be 0.1 and the result is shown in Figure **??**(a).

We first apply the classifier to predict CRC status and compare the ROC-AUC score. To investigate whether these integration methods lead to over-optimistic results, we randomly generate a binary variable from a Bernoulli(0.5) distribution and use it as input for the data integration methods. Then, we apply the classifier to predict this uninformative variable and compare the ROC-AUC score with 0.5. A one-sided DeLong's test was applied to assess whether the ROC-AUC of classifiers trained on integrated data was significantly greater than a random-guess classifier. The null hypothesis is that the model performs no better than random guessing, while the alternative hypothesis states that the ROC-AUC is significantly higher than 0.5. The significance level was set to 0.1. The results are presented in Figure **??**(b).

**Species-level Analysis**

We investigated the prediction performance of random forest and Lasso logistic regression models on species-level integrated data. In total, there are 1,911 species. To compare with the results of (Wirbel et al., 2019), no sample filtering was applied and all 575 samples were included in the analysis. Disease status was used as input for all data integration methods. The taxa similarity graph was constructed by connecting species that belong to the same genus for MetaDICT.

**Prediction Using Lasso Logistic Regression** To compare our results with those reported in Wirbel et al. (2019), we followed the same data processing pipeline. First, we filtered out non-abundant taxa, reducing the number of species from 1,911 to 849. We then applied various data integration methods to reduce batch effects. After adding a pseudocount of $1 \times 10^{-5}$ to the compositional data, we standardized the features as $z$-scores. A Lasso logistic regression model was trained using 10-fold cross-validation, repeated 10 times, as implemented in the `SIAMCAT` R package (Wirbel et al., 2021). The final ROC-AUC score was computed as the average across 100 cross-validated models. We compared the ROC-AUC scores of the Lasso logistic regression model trained on unprocessed data with those trained on data processed by different integration methods. The results are shown in Figure **??**(a).

**Random Forest Prediction for LOSO experiments** We evaluated the performance of random forest classifiers in a leave-one-study-out (LOSO) experiment, where the model was trained on four datasets and tested on the remaining study. All 1911 species were used as input features. Both unprocessed count data and MetaDICT-processed data were used to train the classifier. The random forest algorithm was implemented using the `caret` R package (Kuhn, 2008), with the number of decision trees set to 500. Model parameters were tuned using three-fold cross-validation based on the ROC-AUC metric. The results are presented in Figure **??**(b).

**Microbial Community Detection Results**

| Microbial Community | Genera |
|---|---|
| 1 | *Porphyromonas*, *Peptostreptococcus*, *Lachnoanaerobaculum*, *Anaerococcus*, *Solobacterium*, *Providencia*, *Anaerotruncus*, *Parvimonas*, *Gemella*, *Mobiluncus*, *Helcococcus*, *Tannerella*, unknown genus of *Synergistaceae*. |
| 2 | *Gardnerella*, *Alcaligenes*, *Stenotrophomonas*, *Microbacterium*, *Robinsoniella*, *Anaerostipes*, *Cetobacterium*, *Pseudoflavonifractor*, *Anaeromassilibacillus*, unknown genus of *Succinivibrionaceae*, *Neglecta*. |
| 3 | *Megasphaera*, *Streptococcus*, *Lactobacillus*, unknown genus of *Enterobacteriaceae*, *Salmonella*, *Citrobacter*, *Clostridioides*, *Raoultella*, *Acetobacter*, *Comamonas*, *Edwardsiella*, *Enterococcus*, *Aggregatibacter*, *Weissella*, *Neisseria*, *Aeromonas*, *Acidaminococcus*, *Leptotrichia*, *Selenomonas*, *Veillonella*, *Morganella*, *Kingella*, *Hungatella*, unknown genus of *Erysipelotrichaceae*, *Parabacteroides*, *Oribacterium*, unknown genus of *Prevotellaceae*, *Acidiphilium*, unknown genus of *Peptostreptococcaceae*, *Kluyvera*, unknown genus of *Verrucomicrobia*, unknown genus of *Flavobacteriaceae*, *Butyricimonas*, *Faecalicoccus*, *Holdemania*, *Actinobaculum*, unknown genus of *Methanomethylophilus*, unknown genus of *Tissierellia*, *Succinivibrio*, *Intestinibacter*, *Oxalobacter*, *Catonella*, *Cardiobacterium*, *Stomatobaculum*, *Lautropia*, *Centipeda*, unknown genus of *Selenomonadaceae*, *Mailhella*, unknown genus of *Veillonellaceae*, unknown genus of *Lactobacillales*, unknown genus of *Flavobacteriia*, unknown genus of *Tenericutes*, *Colibacter*, *Niameybacter*. |
| 4 | unknown genus of *Bacillales*, *Propionibacterium*, *Burkholderia*, *Rhodococcus*, *Arcobacter*, *Xanthomonas*, *Carnobacterium*, *Leuconostoc*, *Alloscardovia*, *Brochothrix*, *Dermabacter*, *Finegoldia*, *Paracoccus*, *Terrisporobacter*, *Elusimicrobium*, *Marvinbryantia*, *Abiotrophia*, *Cellulosilyticum*, *Sneathia*, unknown genus of *Coriobacteriales*, *Lawsonella*. |
| 5 | *Pseudomonas*, *Psychrobacter*, *Pasteurella*, *Rothia*, *Parascardovia*, *Paeniclostridium*, *Shuttleworthia*, *Enterorhabdus*, *Mitsuokella*, *Scardovia*, *Rikenella*, *Pseudopropionibacterium*, unknown genus of *Bacilli*. |
| 6 | *Solibacillus*, *Helicobacter*, *Pantoea*, *Phocaeicola*, *Bavariicoccus*, *Acholeplasma*, unknown genus of *Comamonadaceae*, *Desulfovibrio*, *Macrococcus*, *Filifactor*, *Enhydrobacter*, unknown genus of *Bacteroidia*. |

| | |
|---|---|
| 7 | *Sporosarcina*, *Pediococcus*, *Proteus*, *Lysinibacillus*, *Paraclostridium*, *Eggerthia*, *Methanomassiliicoccus*, *Anaerofustis*, *Bulleidia*, *Fenollaria*. |
| 8 | *Ralstonia*, *Eremococcus*, *Meiothermus*, *Facklamia*, *Brevundimonas*, unknown genus of *Rhodospirillales*, unknown genus of *Odoribacteraceae*, unknown genus of *Enterobacterales*, unknown genus of *Fusobacteria*. |
| 9 | *Klebsiella*, *Actinomyces*, *Blautia*, *Eggerthella*, unknown genus of *Lachnospiraceae*, unknown genus of *Clostridiales*, *Dorea*, *Faecalibacterium*, *Eubacterium*, *Bacterium*, *Butyricicoccus*, *Adlercreutzia*, unknown genus of *Ruminococcaceae*, unknown genus of *Saccharibacteria*. |
| 10 | *Escherichia*, *Brachyspira*, *Acinetobacter*, *Peptoniphilus*, *Coprobacillus*, *Parasutterella*, *Varibaculum*, *Levyella*, unknown genus of *Eggerthellales,Bacteria*. |
| 11 | *Dielma*, *Methanobrevibacter*, *Collinsella*, *Senegalimassilia*, *Eikenella*, *Holdemanella*, unknown genus of *Burkholderiales*, unknown genus of *Eggerthellaceae*. |
| 12 | *Serratia*, *Yersinia*, *Campylobacter*, *Treponema*, *Stoquefichus*, *Cryptobacterium*, unknown genus of *Selenomonadales*. |
| 13 | *Oscillibacter*, *Methanobacterium*, *Pyramidobacter*, *Coraliomargarita*, *Alloprevotella*, *Odoribacter*, unknown genus of *Sutterellaceae*, *Merdibacter*. |
| 14 | *Corynebacterium*, *Kosakonia*, *Franconibacter*, *Brevibacterium*, *Pluralibacter*, *Leclercia*. |
| 15 | *Prevotella*, *Desulfovibrio*, *Intestinimonas*, *Paraprevotella*, unknown genus of *Lentisphaerae*. |
| 16 | *Fusobacterium*, *Sutterella*, *Alistipes*, *Phascolarctobacterium*, *Bilophila*, *Akkermansia*, *Anaeroglobus*. |
| 17 | *Hafnia*, unknown genus of *Coriobacteriaceae*, *Corallococcus*, *Methanosphaera*, unknown genus of *Atopobiaceae*, *Phoenicibacter*. |
| 18 | *Coprococcus*, *Atopobium*, *Slackia*, *Olsenella*, *Mogibacterium*, *Granulicatella*. |
| 19 | *Azospirillum*, *Acetobacterium*, *Synergistes*, *Barnesiella*, unknown genus of *Clostridia*. |
| 20 | *Staphylococcus*, *Mycoplasma*, *Butyrivibrio*, unknown genus of *Dehalococcoidales*. |
| 21 | *Bacillus*, *Enorma*, unknown genus of *Clostridiaceae*, *Libanicoccus*, unknown genus of *Methanomicrobia*. |

| | |
|---|---|
| 22 | *Achromobacter*, *Sodalis*, *Tropheryma*, *Dolosigranulum*, *Pseudoramibacter*. |
| 23 | *Paraburkholderia*, *Cronobacter*, *Cedecea*, *Anoxybacillus*, *Dysgonomonas*. |
| 24 | *Enterobacter*, *Haemophilus*, *Megamonas*, unknown genus of *Pasteurellaceae*. |
| 25 | *Lactococcus*, *Faecalitalea*, *Turicibacter*, unknown genus of *Massiliomicrobiota*. |
| 26 | *Shewanella*, *Desulfotomaculum*, *Anaerobiospirillum*, unknown genus of *Melainabacteria*. |
| 27 | unknown genus of *Bacteroidales*, *Porphyromonadaceae*, *Roseburia*, *Bacteroidaceae*. |
| 28 | *Bacteroides*, *Flavonifractor*, *Dialister*, *Tyzzerella*. |
| 29 | *Bifidobacterium*, *Ruminococcus*, unknown genus of *Firmicutes*, *Subdoligranulum*. |
| 30 | *Succinatimonas*, unknown genus of *Rhodospirillaceae*, unknown genus of *Saccharomycetaceae*. |

Table S2: Microbial Community Detection Results

#### Setting for Section 2.7

| Study | Region | No. of samples | Phenotype(s) | Reference |
|---|---|---|---|---|
| Chaput | France | 26 | NR 17/R 9 | Chaput et al. (2017) |
| Gopalakrishnan | United States | 43 | NR 13/R 30 | Gopalakrishnan et al. (2018) |
| Frankel | United States | 39 | NR 15/R 24 | Frankel et al. (2017) |
| Matson | United States | 42 | NR 26/R 16 | Matson et al. (2018) |
| Peters | United States | 24 | NR 2/R 22 | Peters et al. (2019) |

Table S3: **Five PD-1 therapy microbiome datasets included in the data integration.**

We integrate five 16S rRNA datasets of PD-1 treated patients and aim to study the association between gut microbiome and responses to therapy (Table S3). 174 stool samples were collected from PD-1-treated patients with melanoma at the beginning of therapy and sequenced by different protocols for different studies. Sequences are processed with the same bioinformatics pipeline in Shaikh et al. (2021). The union of taxa across datasets is treated as an integrated feature, and we filter out taxa that appear in less than 10% samples of concatenated data. In total, there are 881 taxa at the OTU level. Only response status is included in all the meta tables.

**Implementation of MetaDICT** We selected parameters of MetaDICT as $\alpha = 0.01$, $\beta = 0.01$, $\gamma = 1$. Since phylogenetic tree is not available, we used taxonomic information to construct taxa similarity graph. In this graph, taxa within the same family were connected with each other.

**OTU-Level Association Analysis** LinDA is applied to test the association between genera abundance and response status. The Benjamini-Hochberg procedure adjusts $p$-values with a significance level of $\alpha = 0.1$.

**Response Status Prediction** The random forest model has the same setup as Section 2.6. The classifier is trained on all but one data set and evaluated on the left one. The two-layer neural network classifier is implemented using `pytorch`. In the first fully connected layer, the features are reduced from 881 dimensions to 64 dimensions with the ReLU activation function. The second layer is also a fully connected layer with a sigmoid function. Cross-entropy loss is used in classifier training.

We first apply the classifier to predict response status and compare the ROC-AUC score. To investigate whether these integration methods lead to over-optimistic results, we randomly generate a binary variable from a Bernoulli(0.5) distribution and use it as input for the data integration methods. Then, we apply the classifier to predict this uninformative variable and compare the ROC-AUC score with 0.5.

#### References


N. T. Baxter, M. T. Ruffin, M. A. Rogers, and P. D. Schloss. Microbiota-based model improves the sensitivity of fecal immunochemical test for detecting colonic lesions. *Genome Medicine*, 8:1–10, 2016.

V. D. Blondel, J. Guillaume, R. Lambiotte, and E. Lefebvre. Fast unfolding of communities in large networks. *Journal of Statistical Mechanics: Theory and Experiment*, 2008(10): P10008, 2008.

N. Chaput, P. Lepage, C. Coutzac, E. Soularue, K. Le Roux, C. Monot, L. Boselli, E. Routier, L. Cassard, M. Collins, T. Vaysse, L. Marthey, A. Eggermont, V. Asvatourian, E. Lanoy, C. Mateus, C. Robert, and F. Carbonnel. Baseline gut microbiota predicts clinical response and colitis in metastatic melanoma patients treated with ipilimumab. *Annals of Oncology*, 28(6):1368–1379, 2017.

E. R. DeLong, D. M. DeLong, and D. L. Clarke-Pearson. Comparing the areas under two or more correlated receiver operating characteristic curves: a nonparametric approach. *Biometrics*, pages 837–845, 1988.

Q. Feng, S. Liang, H. Jia, A. Stadlmayr, L. Tang, Z. Lan, D. Zhang, H. Xia, X. Xu, Z. Jie, L. Su, X. Li, X. Li, J. Li, L. Xiao, U. Huber-Schönauer, D. Niederseer, X. Xu, J. Al-Aama, H. Yang, J. Wang, K. Kristiansen, M. Arumugam, H. Tilg, C. Datz, and J. Wang. Gut microbiome development along the colorectal adenoma–carcinoma sequence. *Nature Communications*, 6(1):6528, 2015.

A. E. Frankel, L. A. Coughlin, J. Kim, T. W. Froehlich, Y. Xie, E. P. Frenkel, and A. Y. Koh. Metagenomic shotgun sequencing and unbiased metabolomic profiling identify specific human gut microbiota and metabolites associated with immune checkpoint therapy efficacy in melanoma patients. *Neoplasia*, 19(10):848–855, 2017.

S. M. Gibbons, C. Duvallet, and E. J. Alm. Correcting for batch effects in case-control microbiome studies. *PLoS Computational Biology*, 14(4):e1006102, 2018.

V. Gopalakrishnan, N. C. Spencer, L. Nezi, A. Reuben, C. M. andrews, V. T. Karpinets, A. P. Prieto, D. Vicente, K. Hoffman, C. S. Wei, P. A. Cogdill, L. Zhao, W. C. Hudgens, S. D. Hutchinson, T. Manzo, M. Petaccia de Macedo, T. Cotechini, T. Kumar, S. W. Chen, M. S. Reddy, R. Szczepaniak Sloane, J. Galloway-Pena, H. Jiang, L. P. Chen, J. E. Shpall, K. Rezvani, M. A. Alousi, F. R. Chemaly, S. Shelburne, M. L. Vence, C. P. Okhuysen, B. V. Jensen, G. A. Swennes, F. McAllister, M. E. Riquelme Sanchez, Y. Zhang, E. Le Chatelier, L. Zitvogel, N. Pons, L. J. Austin-Breneman, E. L. Haydu, M. E. Burton, M. J. Gardner, E. Sirmans, J. Hu, J. A. Lazar, T. Tsujikawa, A. Diab, H. Tawbi, C. I. Glitza, J. W. Hwu, P. S. Patel, E. S. Woodman, N. R. Amaria, A. M. Davies, E. J. Gershenwald, P. Hwu, E. J. Lee, J. Zhang, M. L. Coussens, A. Z. Cooper, A. P. Futreal, R. C. Daniel, J. N. Ajami, F. J. Petrosino, T. M. Tetzlaff, P. Sharma, P. J. Allison, R. R. Jenq, and A. J.

Wargo. Gut microbiome modulates response to anti–pd-1 immunotherapy in melanoma patients. *Science*, 359(6371):97–103, 2018.

Y. He, W. Wu, H. Zheng, P. Li, D. McDonald, H. Sheng, M. Chen, Z. Chen, G. Ji, Z. Zheng, P. Mujagond, X. Chen, Z. Rong, P. Chen, L. Lyu, X. Wang, C. Wu, N. Yu, Y. Xu, J. Yin, J. Raes, R. Knight, W. Ma, and H. Zhou. Regional variation limits applications of healthy gut microbiome reference ranges and disease models. *Nature Medicine*, 24(10):1532–1535, 2018.

M. Kuhn. Building predictive models in r using the caret package. *Journal of Statistical Software*, 28:1–26, 2008.

S. Ma, B. Ren, H. Mallick, Y. S. Moon, E. Schwager, S. Maharjan, T. L. Tickle, Y. Lu, R. N. Carmody, E. A. Franzosa, L. Janson, and C. Huttenhower. A statistical model for describing and simulating microbial community profiles. *PLoS Computational Biology*, 17 (9):e1008913, 2021.

V. Matson, J. Fessler, R. Bao, T. Chongsuwat, Y. Zha, M.-L. Alegre, J. J. Luke, and T. F. Gajewski. The commensal microbiome is associated with anti–pd-1 efficacy in metastatic melanoma patients. *Science*, 359(6371):104–108, 2018.

A. Milanese, D. R. Mende, L. Paoli, G. Salazar, H. Ruscheweyh, M. Cuenca, P. Hingamp, R. Alves, P. I. Costea, L. P. Coelho, T. S. B. Schmidt, A. Almeida, A. L. Mitchell, R. D. Finn, J. Huerta-Cepas, P. Bork, G. Zeller, and S. Sunagawa. Microbial abundance, activity and population genomic profiling with motus2. *Nature Communications*, 10(1):1014, 2019.

B. A. Peters, M. Wilson, U. Moran, A. Pavlick, A. Izsak, T. Wechter, J. S. Weber, I. Osman, and J. Ahn. Relating the gut metagenome and metatranscriptome to immunotherapy responses in melanoma patients. *Genome Medicine*, 11:1–14, 2019.

X. Robin, N. Turck, A. Hainard, N. Tiberti, F. Lisacek, J.-C. Sanchez, and M. Müller. proc: an open-source package for r and s+ to analyze and compare roc curves. *BMC bioinformatics*, 12:1–8, 2011.

Y. F. Shaikh, R. J. White, J. J. Gills, T. Hakozaki, C. Richard, B. Routy, Y. Okuma, M. Usyk, A. Pandey, S. J. Weber, J. Ahn, J. E. Lipson, J. Naidoo, M. D. Pardoll, and L. C. Sears. A uniform computational approach improved on existing pipelines to reveal microbiome biomarkers of nonresponse to immune checkpoint inhibitors. *Clinical Cancer Research*, 27(9):2571–2583, 2021.

E. Vogtmann, X. Hua, G. Zeller, S. Sunagawa, A. Y. Voigt, R. Hercog, J. J. Goedert, J. Shi, P. Bork, and R. Sinha. Colorectal cancer and the human gut microbiome: reproducibility with whole-genome shotgun sequencing. *PloS One*, 11(5):e0155362, 2016.

J. Wirbel, P. T. Pyl, E. Kartal, K. Zych, A. Kashani, A. Milanese, J. S. Fleck, A. Y. Voigt, A. Palleja, R. Ponnudurai, S. Sunagawa, L. P. Coelho, P. Schrotz-King, E. Vogtmann,

N. Habermann, E. Niméus, A. M. Thomas, P. Manghi, S. Gandini, D. Serrano, S. Mizutani, H. Shiroma, S. Shiba, T. Shibata, S. Yachida, T. Yamada, L. Waldron, A. Naccarati, N. Segata, R. Sinha, C. M. Ulrich, H. Brenner, M. Arumugam, P. Bork, and G. Zeller. Meta-analysis of fecal metagenomes reveals global microbial signatures that are specific for colorectal cancer. *Nature Medicine*, 25(4):679–689, 2019.

J. Wirbel, K. Zych, M. Essex, N. Karcher, E. Kartal, G. Salazar, P. Bork, S. Sunagawa, and G. Zeller. Microbiome meta-analysis and cross-disease comparison enabled by the siamcat machine learning toolbox. *Genome Biology*, 22(1):93, 2021.

J. Yu, Q. Feng, S. Wong, D. Zhang, Q. yi Liang, Y. Qin, L. Tang, H. Zhao, J. Stenvang, Y. Li, X. Wang, X. Xu, N.g Chen, W. Wu, J. Al-Aama, H. J. Nielsen, P. Kiilerich, B. Jensen, T. Yau, Z. Lan, H. Jia, J. Li, L. Xiao, T. Lam, S. Ng, A. Cheng, V. Wong, F. Chan, Xun Xu, H. Yang, L. Madsen, C. Datz, H. Tilg, J. Wang, N. Brünner, K. Kristiansen, J. Arumugam, M.and Sung, and J. Wang. Metagenomic analysis of faecal microbiome as a tool towards targeted non-invasive biomarkers for colorectal cancer. *Gut*, 66(1):70–78, 2017.

J. P. Zackular, M. A. Rogers, M. T. Ruffin IV, and P. D. Schloss. The human gut microbiome as a screening tool for colorectal cancer. *Cancer Prevention Research*, 7(11):1112–1121, 2014.

G. Zeller, J. Tap, A. Y. Voigt, S. Sunagawa, J. R. Kultima, P. I. Costea, A. Amiot, J. Böhm, F. Brunetti, N. Habermann, R. Hercog, M. Koch, A. Luciani, D. R. Mende, M. A. Schneider, P. Schrotz-King, C. Tournigand, J. Tran Van Nhieu, T. Yamada, J. Zimmermann, V. Benes, M. Kloor, C. M. Ulrich, M. von Knebel Doeberitz, I. Sobhani, and P. Bork. Potential of fecal microbiota for early-stage detection of colorectal cancer. *Molecular Systems Biology*, 10(11):766, 2014.
