## Supplemental Figures for "Microbiome Data Integration via Shared Dictionary Learning"

### Supplementary Figures to "Microbiome Data Integration via Shared Dictionary Learning"

Bo Yuan* and Shulei Wang*
University of Illinois at Urbana-Champaign

(June 12, 2025)

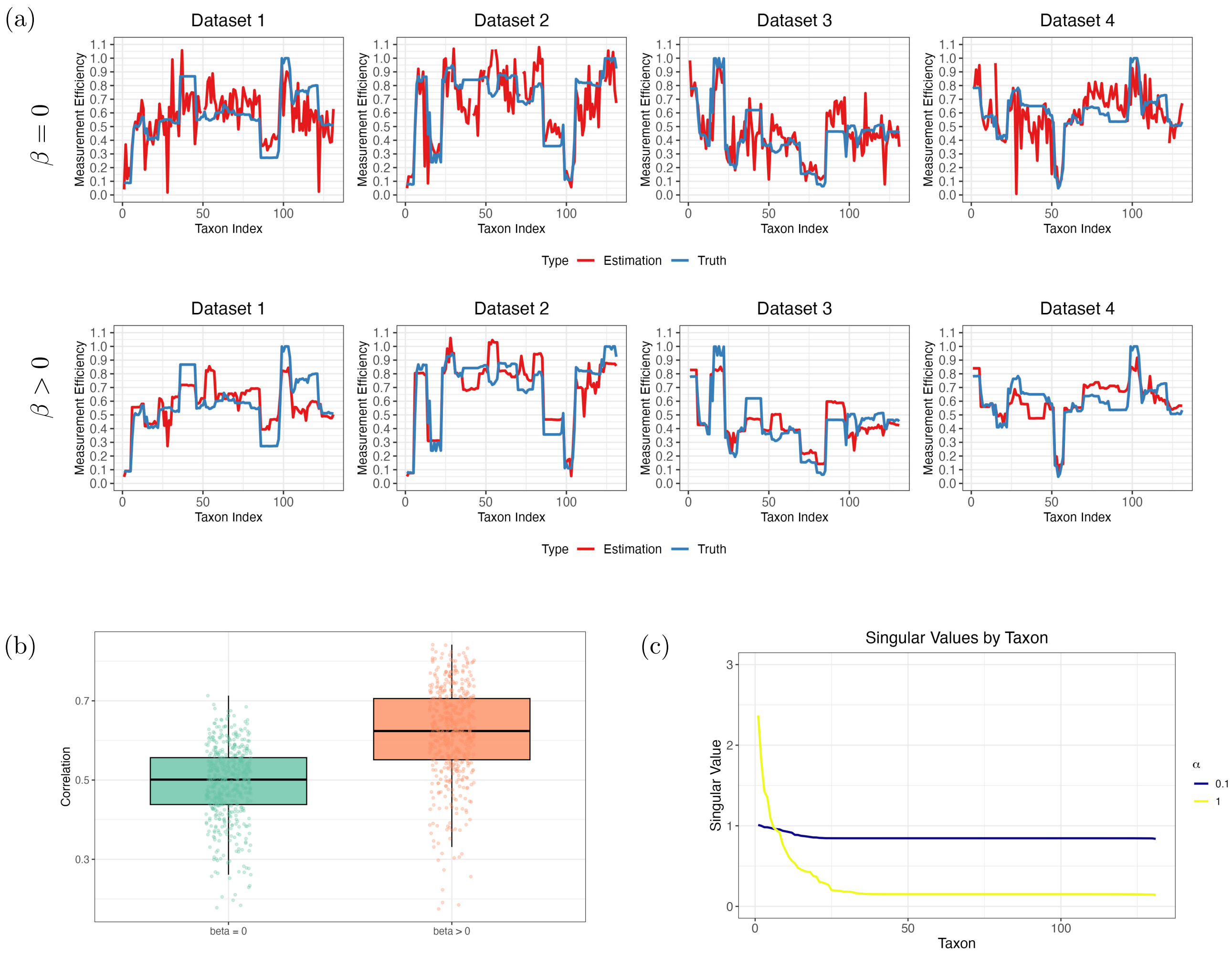


Figure S1: **Effects of tuning parameters $\alpha$ and $\beta$.** In Figures (a) and (b), we demonstrate the importance of incorporating taxa sequence similarity into the estimation of measurement efficiency. In Figure (a), the estimated measurement efficiency is compared with the ground truth in a simulation integrating four datasets, and the results show that using taxa sequence similarity ($\beta > 0$) yields estimates that are both more accurate and less noisy. Figure (b) compares the Pearson correlation between the estimated and true measurement efficiencies for the cases of $\beta > 0$ and $\beta = 0$, further highlighting the benefits of including taxa sequence similarity. In Figure (c), the impact of the parameter $\alpha$ is illustrated: the ranked singular values of the learned shared dictionary $D$ are plotted, showing that a larger $\alpha$ results in a lower rank of $D$.

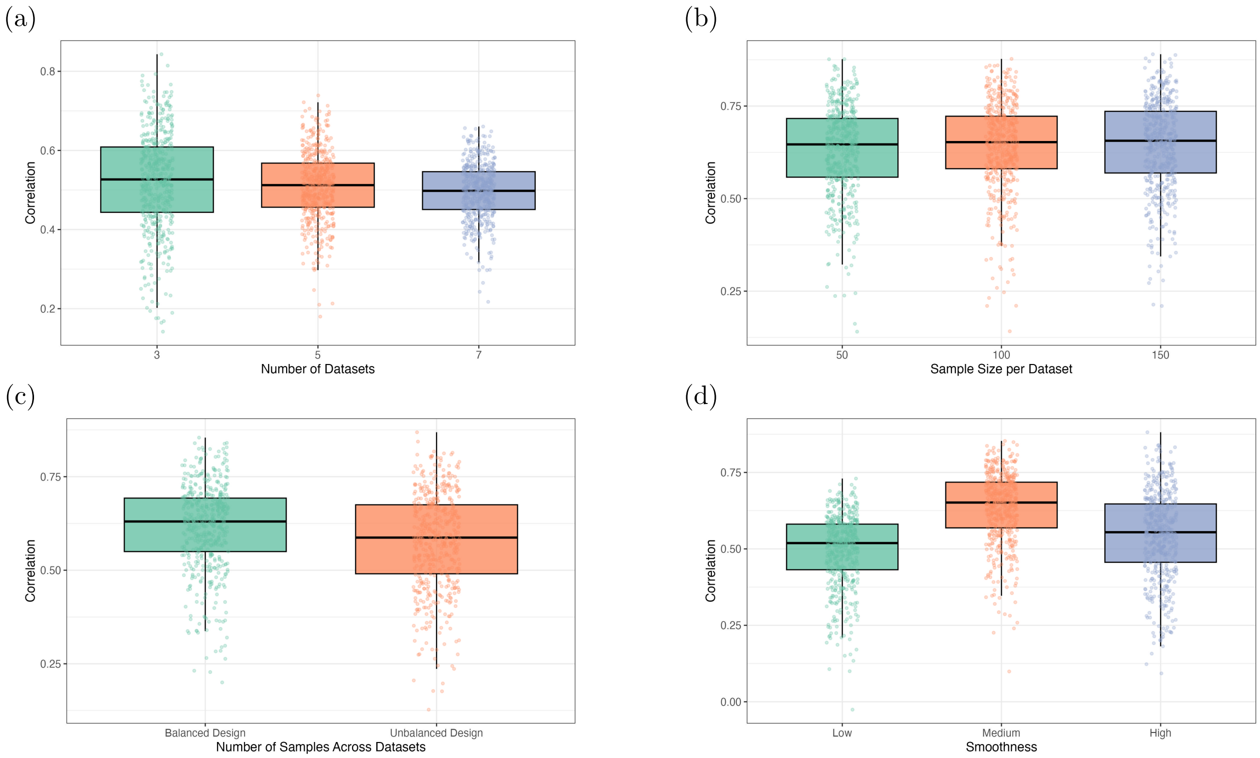


Figure S2: **Pearson correlation between estimated and true measurement efficiency in a wide range of experiment settings.** This set of experiments aims to investigate the robustness of measurement efficiency estimation under various settings. The $y$-axis is the Pearson correlation between estimated and true measurement efficiencies. The experimental settings vary by (a) the number of datasets, (b) the sample size per dataset, (c) the balance of dataset sizes, and (d) the smoothness of the measurement efficiency. MetaDICT can recover the measurement efficiencies robustly under different settings.

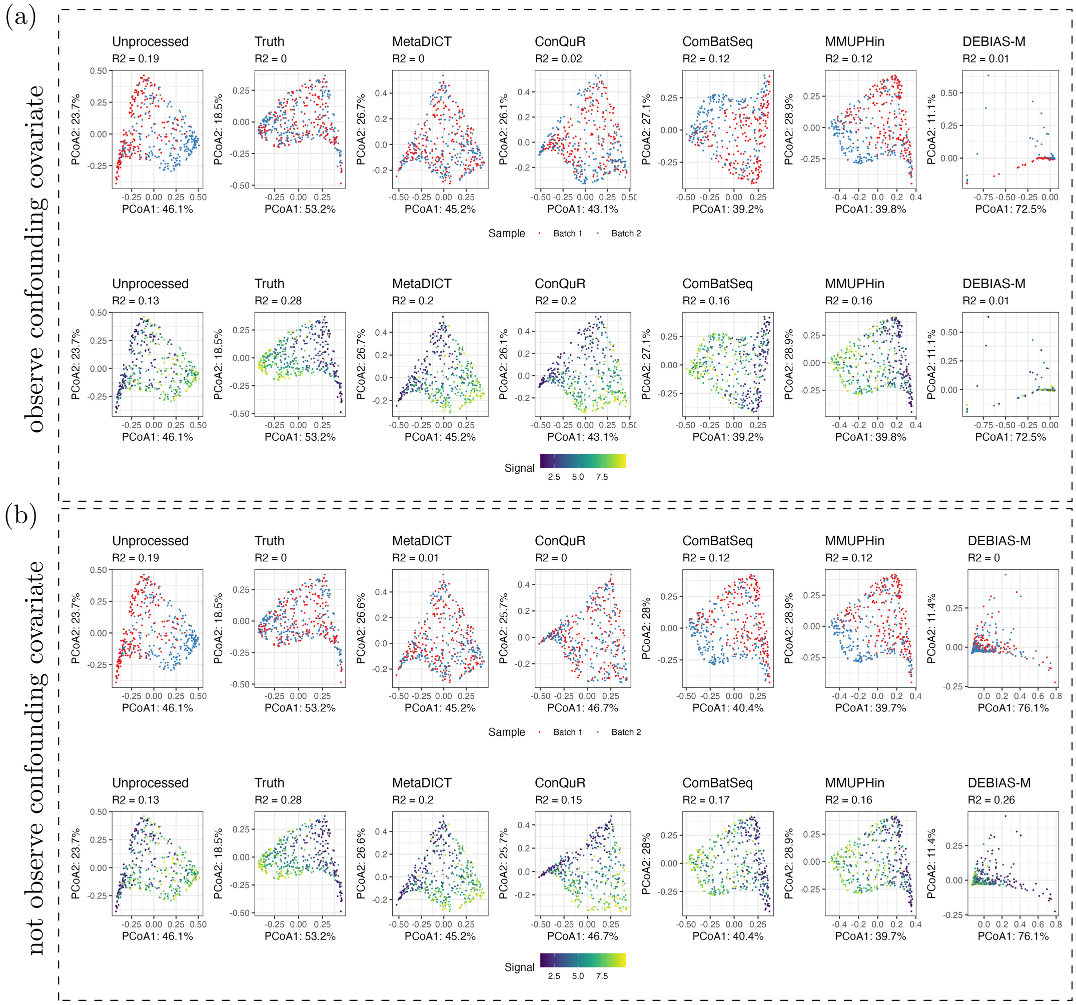


Figure S3: **Comparison of different data integration methods when the biological variable is continuous.** In this experiment, we simulated two batches of data, and the synthetic samples were associated with a randomly generated continuous variable. The unprocessed panels display the simulated data with batch effects, while the truth panels show the batch-effect-free data (i.e., the microbial absolute abundance). Figure (b) shows the PCoA plots and PERMANOVA $R^2$ values when the biological variable is not observed in advance, whereas Figure (a) presents the results when the biological variable is used as input for all methods. The data integration methods aim to recover the truth panels as accurately as possible. Note that PLSDA-batch, Percentile Normalization, and scANVI are excluded because they do not support continuous outcome variables. Bray-Curtis dissimilarity is used for all methods. The result indicates that MetaDICT can effectively correct batch effects and preserve the biological variation with continuous biological variable.

(a) Confounding covariate is observed:

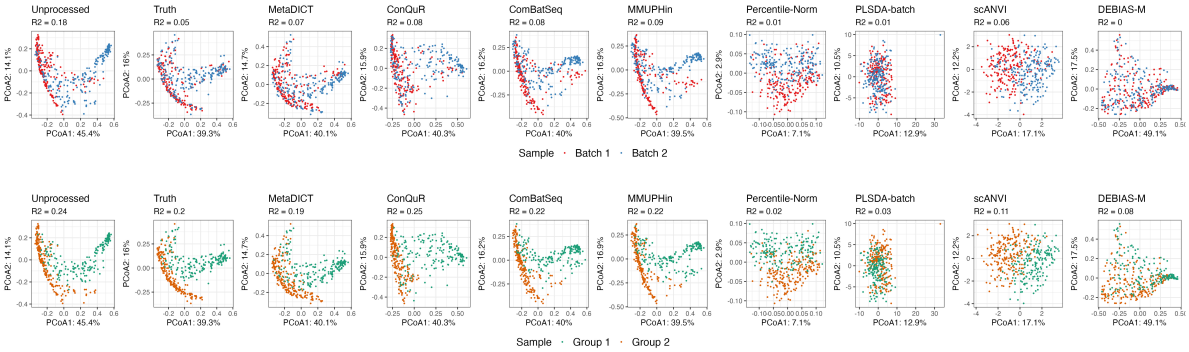


(b) Confounding covariate is not observed:

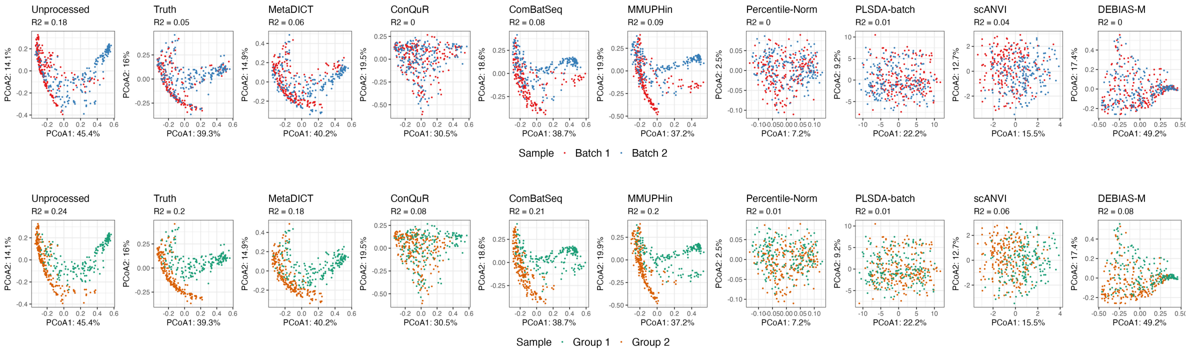


Figure S4: **Comparison of different data integration methods with both batch effects and a distribution shift in absolute abundance across datasets.** In this experiment, we simulated two batches of data, and the synthetic samples were associated with a randomly generated binary variable (Group 1 and Group 2). Specifically, there is a distribution shift in absolute abundance; that is, the distribution of the binary variable is not identical across the two batches. The unprocessed panels display the simulated data with batch effects, while the truth panels show the batch-effect-free data (i.e., the microbial absolute abundance). Figure (b) shows the PCoA plots and PERMANOVA $R^2$ values when the biological variable is not observed in advance, whereas Figure (a) presents the results when the biological variable is used as input for all methods. The data integration methods aim to recover the truth panels as accurately as possible. Euclidean distance is used for PLSDA-batch and scANVI, while Bray-Curtis dissimilarity is applied for the remaining methods. This results indicates that MetaDICT can avoid overcorrection even when confounding variable is not observed.

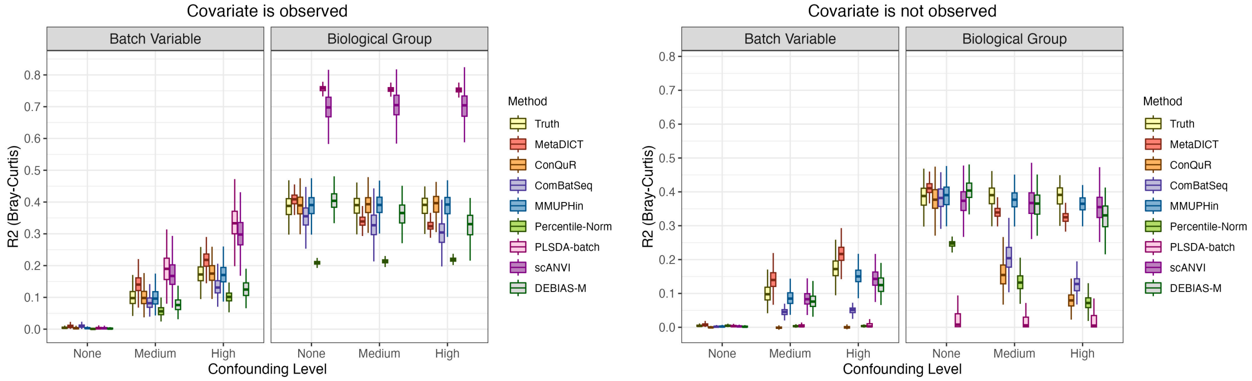


Figure S5: **Comparison of different data integration methods with no batch effects but a distribution shift in covariate.** The figures consider an ideal scenario in which there is a distribution shift in absolute abundance but no batch effect across datasets (i.e., the measurement efficiency is the same across batches). We compare the performance of eight data integration methods using the $R^2$ statistic from PERMANOVA, under varying levels of confounding between the biological variable and batch. The left panel shows results when the biological variable is not observed in advance, while the right panel shows results when the biological variable is observed in advance and used in data integration methods. The experiments are repeated 500 times, and boxplots of the resulting $R^2$ values are presented. The results demonstrate that MetaDICT performs robustly and avoids overcorrection in the presence of a distribution shift in absolute abundance. Euclidean distance is used for PLSDA-batch and scANVI, while Bray-Curtis dissimilarity is applied for the remaining methods.

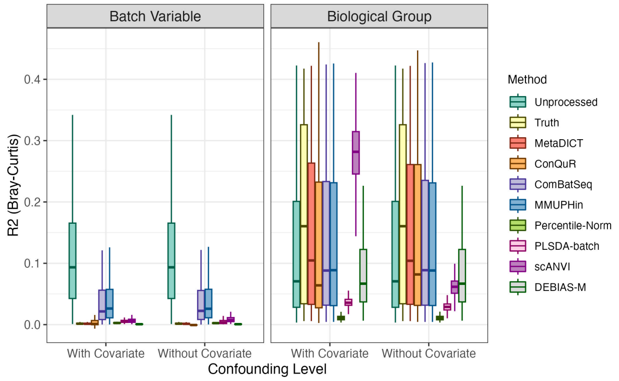


Figure S6: **Comparison of different data integration methods with batch effects and no distribution shift in covariate.** In this experiment, we simulated two batches of data, and the synthetic samples were associated with a randomly generated binary variable (Group 1 and Group 2). We compare the performance of eight data integration methods using the $R^2$ statistic from PERMANOVA under two conditions: when the biological variable is not observed in advance, and when it is observed and incorporated into the data integration methods. The figures show boxplots of the resulting $R^2$ values in the repeated 500 experiments. The results indicate that MetaDICT can preserve biological variation while effectively removing batch effects. Euclidean distance is used for PLSDA-batch and scANVI, while Bray-Curtis dissimilarity is applied for the remaining methods.

(a) Confounding covariate is observed:

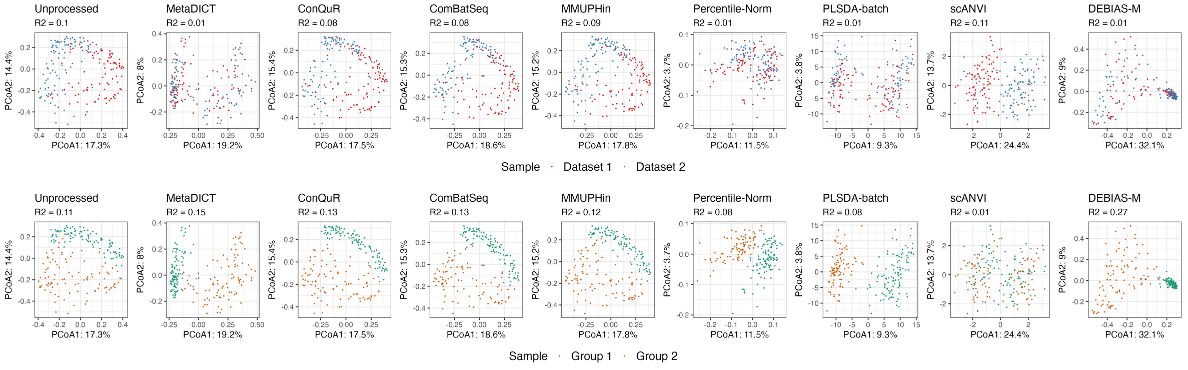


(b) Confounding covariate is not observed:

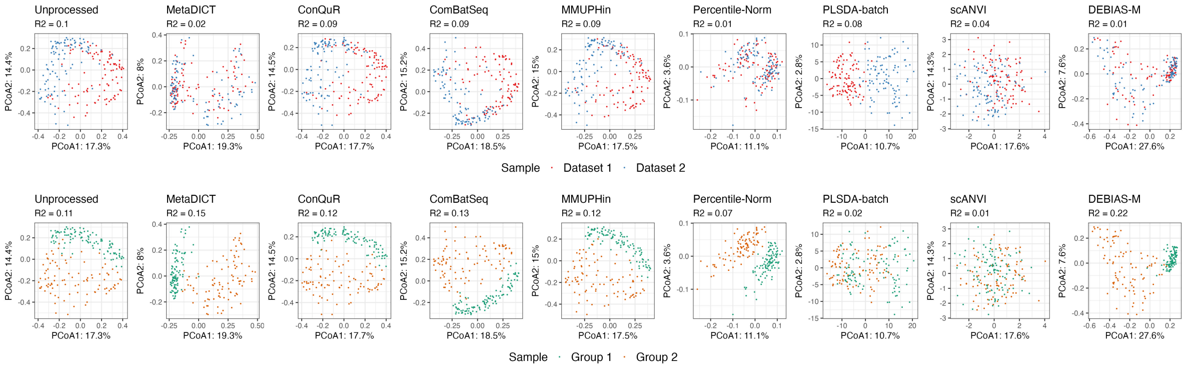


Figure S7: **Comparison of different data integration methods under sparseDOSSA simulation setting.** We use SparseDOSSA2 to generate simulated data. Control samples from one real dataset are used to fit the basis model for simulated dataset 1, while control samples from another dataset are used for simulated dataset 2. Instead of artificially simulating batch effects, we leverage the natural variation between the two datasets as batch effects. As a result, we do not have ground-truth batch-effect-free datasets. Microbial loads are varied according to a binary biological covariate that is independent of batch. The figures display PCoA plots and the $R^2$ statistic from PERMANOVA for scenarios where the biological variable is unobserved (Figure (b)) and observed (Figure (a)). This setting is similar to that used in Figure 2(c) and 2(d). Since the batch variable is independent of the biological covariate, the amount of batch effect can be represented by the variability explained by the batch variable, while the amount of biological variation can be represented by the variability explained by the biological covariate. MetaDICT and DEBIAS-M can effectively reduce batch effects while preserve biological variations. Euclidean distance is used for PLSDA-batch and scANVI, while Bray-Curtis dissimilarity is applied for the remaining methods.

(a) Confounding covariate is observed:

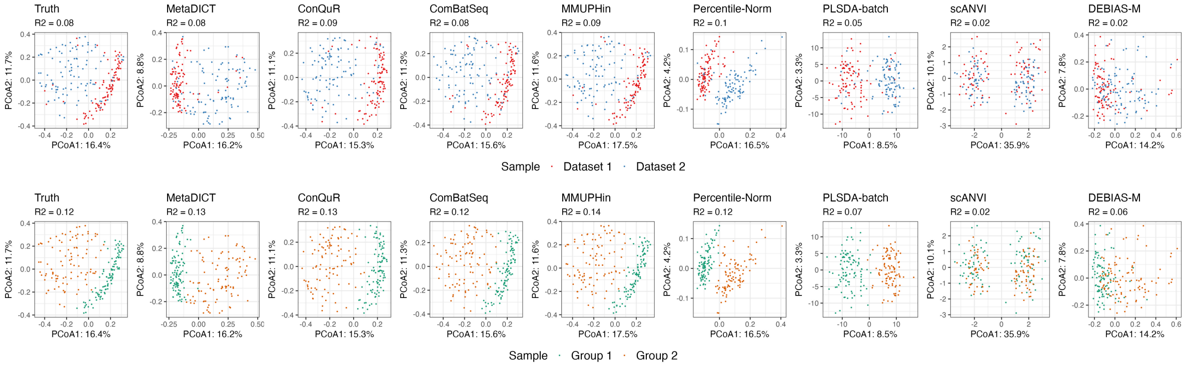


(b) Confounding covariate is not observed:

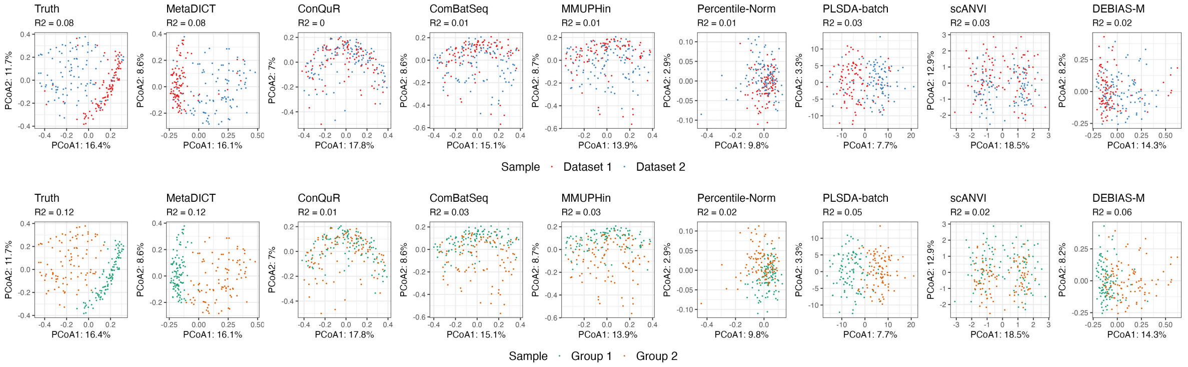


Figure S8: **Comparison of different data integration methods when there is no batch effect (measurement efficiency is the same across batches) but a distribution shift in absolute abundance under sparseDOSSA simulation setting.** The figures consider an ideal scenario in which there is a distribution shift in absolute abundance but no batch effect across datasets. We simulated two datasets using a single model fitted on control samples from a real dataset. Microbial loads are varied according to a binary biological covariate that is confounded with batch (i.e., the biological covariate is imbalanced across batches). The figures display PCoA plots and the $R^2$ statistic from PERMANOVA for scenarios where the biological variable is unobserved (Figure (b)) and observed (Figure (a)). All existing methods suffer from overcorrection when the confounding covariate is not observed, while MetaDICT remains robust. Euclidean distance is used for PLSDA-batch and scANVI, while Bray-Curtis dissimilarity is applied for the remaining methods.

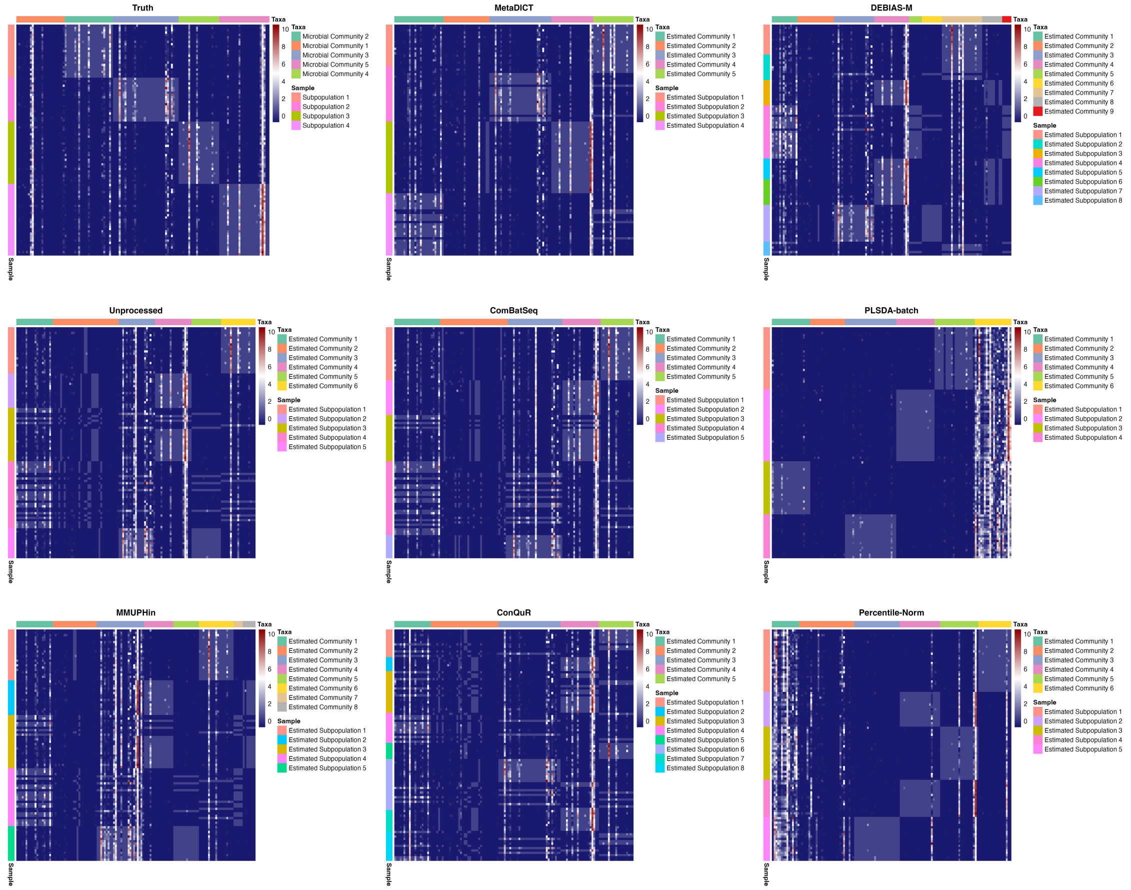


Figure S9: **MetaDICT reveals the biclustering structure of microbiome data via embeddings.** We generated synthetic datasets with five microbial communities and four sample subpopulations, and investigate whether data integration methods can help reveal these biclustering patterns. The figures show heatmaps of absolute abundance, where columns represent taxa and rows represent samples. Both rows and columns have been reordered based on the clustering results obtained from each method. Most methods tend to overestimate the number of microbial communities, while MetaDICT reveals this biclustering result effectively.

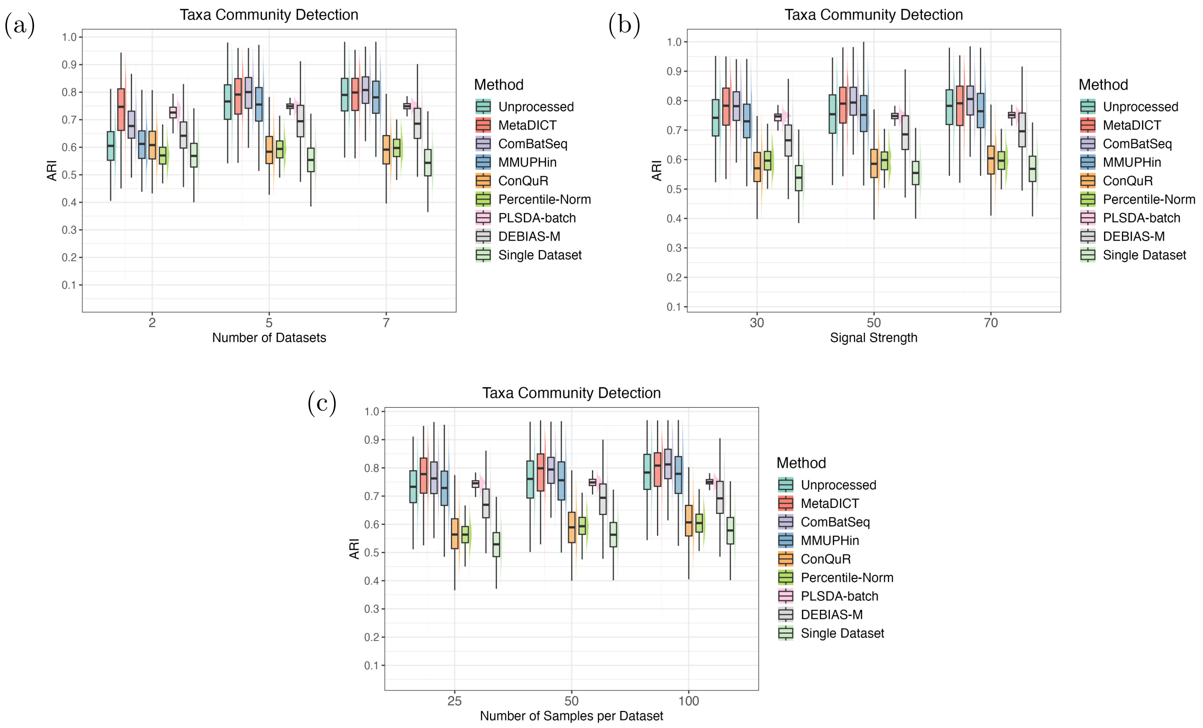


Figure S10: **Comparison of community detection at taxa level when different data integration methods are applied.** The taxa embeddings in MetaDICT can be used in microbial community detection. The figures compare the taxa community detection accuracy of different methods and present boxplots of the adjusted Rand index across 500 repeated experiments, where the number of datasets (Figure (a)), signal strength (Figure (b)), and sample size per dataset (Figure (c)) are varied. MetaDICT consistently performs well in identifying microbial community patterns.

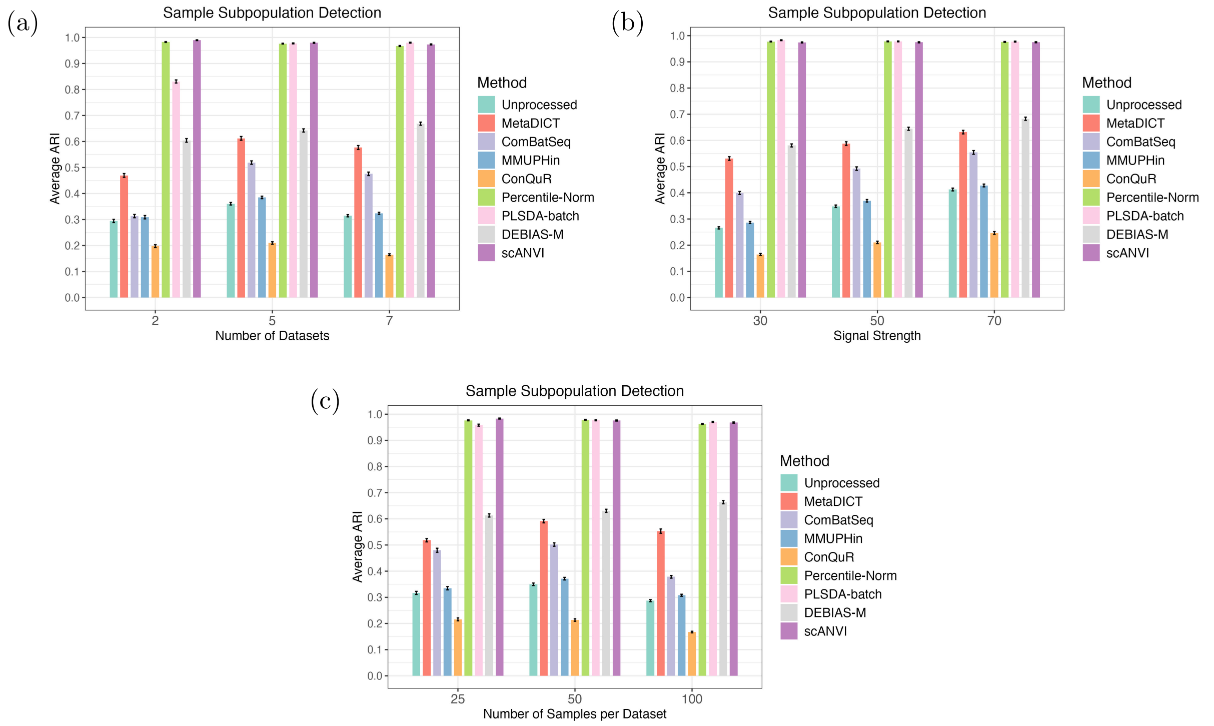


Figure S11: **Comparison of community detection at sample level when different data integration methods are applied.** The sample representations generated by MetaDICT can be used to detect subpopulations. The figures compare the sample community detection accuracy of different methods and present the average adjusted Rand index across 500 repeated experiments, where the number of datasets (Figure (a)), signal strength (Figure (b)), and sample size per dataset (Figure (c)) are varied.

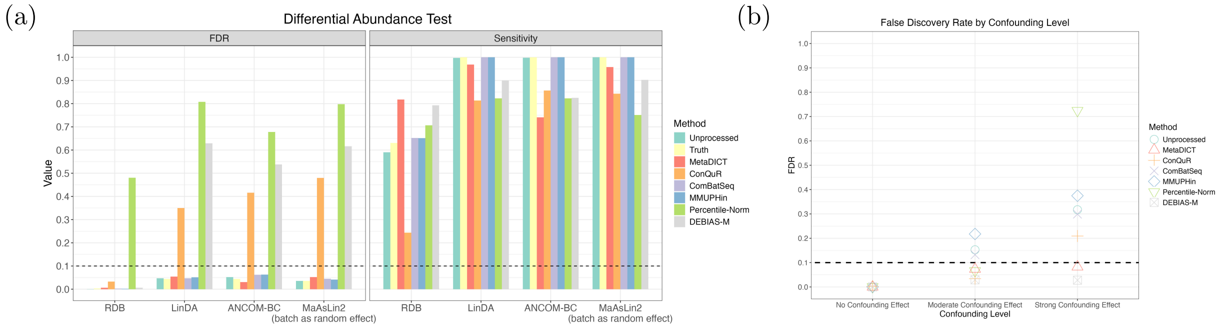


Figure S12: **Comparison of integrative methods on differential abundance tests.** Figure (a) compares false discovery rate control and power(sensitivity) when the covariate of interest is associated with microbial compositions and independent of batches. Most data integration methods can successfully correct the batch effects, and thus, the differential abundance tests on the integrated data set can control the false discovery well and maintain a decent power. In Figure (b), we compare false discovery rate control across methods when the covariate is independent of microbial composition but confounded with batches; for this analysis, control samples from three real studies are treated as different studies, and a binary variable is randomly generated. Due to the confounding effect, batch effect correction becomes more challenging and uncorrected batch effects lead to an inflated false discovery rate. MetaDICT and DEBIAS-M perform more robustly than other methods, especially under the strong confounding setting.

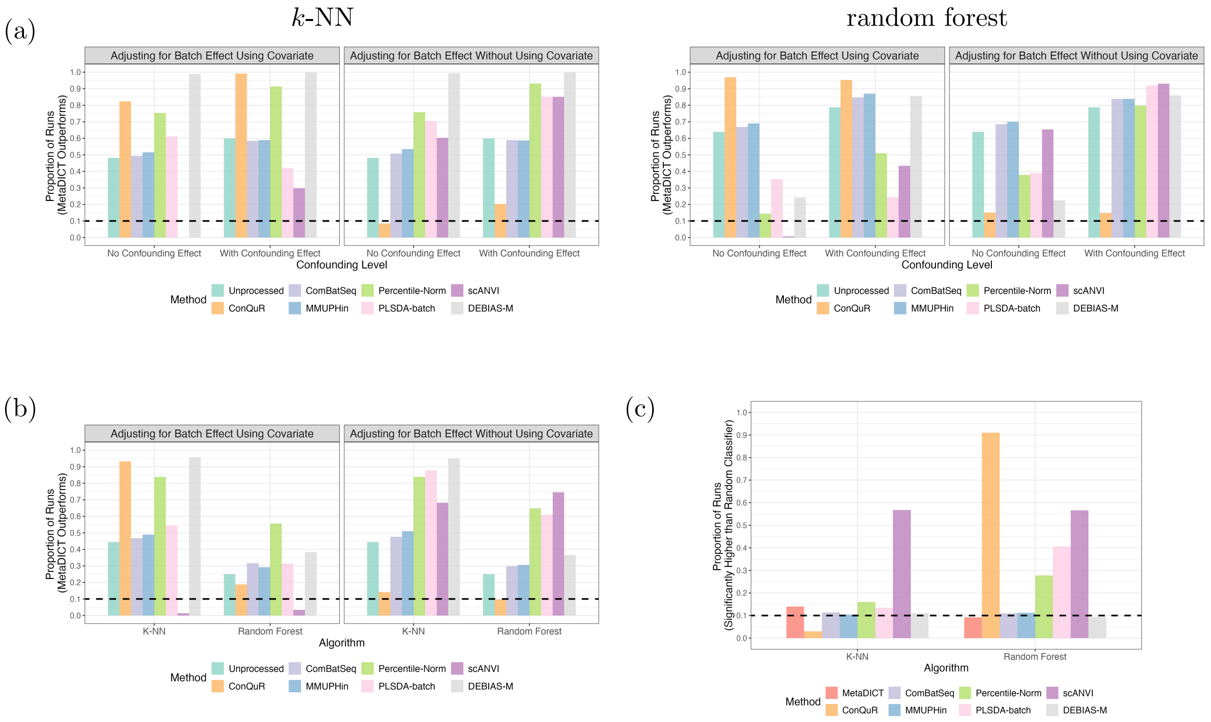


Figure S13: **Comparison of ROC-AUCs of $k$-NN and random forest classifiers using one-sided DeLong's test.** Figure (a) compares the ROC-AUCs of $k$-NN (left panel) and random forest classifiers (right panel) in a transfer learning setting, where the training and testing data sets come from different studies. The $y$-axis shows the proportion of experiments in which the ROC-AUC of the classifier trained on MetaDICT-integrated data is significantly higher than that of the comparison classifier (i.e., the $p$-value from a one-sided DeLong's test is less than 0.1). Figure (b) compares the ROC-AUCs of classifiers trained on MetaDICT-integrated data versus those trained using other integration methods, in scenarios where the training and testing data sets come from the same integrated data set. The classifier trained on MetaDICT-corrected data achieves higher ROC-AUCs than most other methods. Figure (c) considers a negative control setting in which the outcome is independent of microbial composition but is still used during data integration. We test the null hypothesis that the ROC-AUC of the classifier trained on integrated data is greater than that of a random guess classifier. The $y$-axis shows the proportion of experiments in which the null hypothesis is rejected at a significance level of 0.1. Percentile Normalization, PLSDA-batch, scANVI, and ConQuR lead to an over-optimistic random forest classifier, while scANVI also results in an over-optimistic $k$-NN classifier.

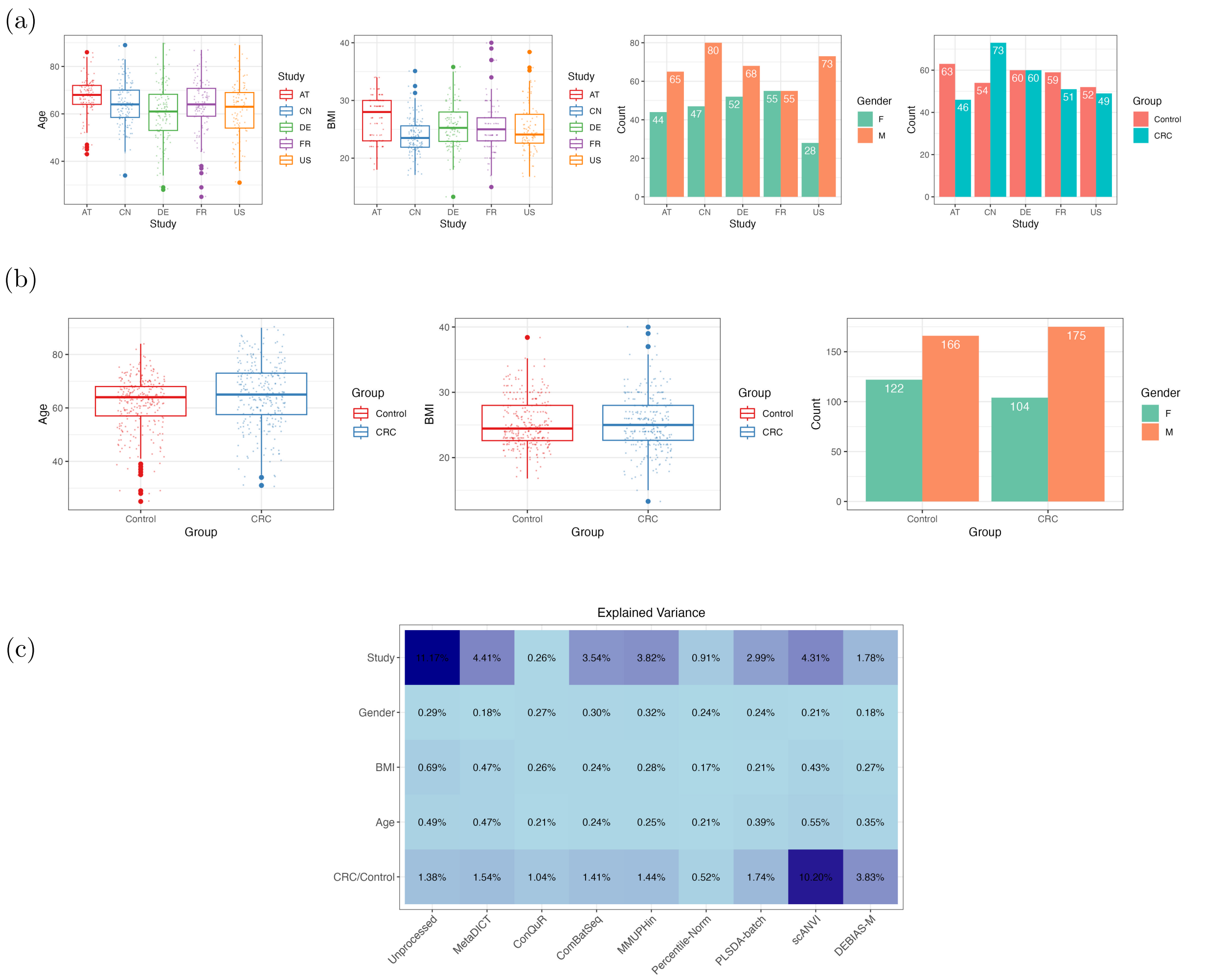


Figure S14: **Comparison of covariates' distribution across different studies and disease status in the integrative analysis of colorectal cancer studies.** Figure (a) compares the covariates' distribution across different studies, including age, BMI, gender and disease status. Figure (b) shows the boxplots of age, BMI, and gender when comparing CRC and control groups. Figure (c) shows variation explained by covariates measured by $R^2$ in PERMANOVA. Euclidean distance is used for PLSDA-batch and scANVI while Bray-Curtis dissimilarity is applied for the rest methods. The study variable had a dominant effect on the microbial profiles for unprocessed data.

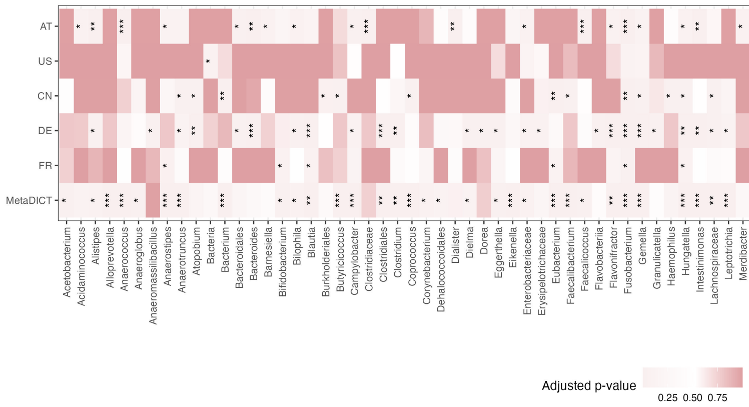


Figure S15: **Differential abundance analysis in meta-analysis of CRC.** Figure presents differentially abundant genera detected by each individual study and the integrated data using LinDA. Differentially abundant genera are marked with an $*$ and adjusted $p$-values are colored. Genera with an adjusted $p$-value smaller than 0.001 are marked with ***; those with an adjusted $p$-value smaller than 0.01 are marked with **; and those with an adjusted $p$-value smaller than 0.1 are marked with *.

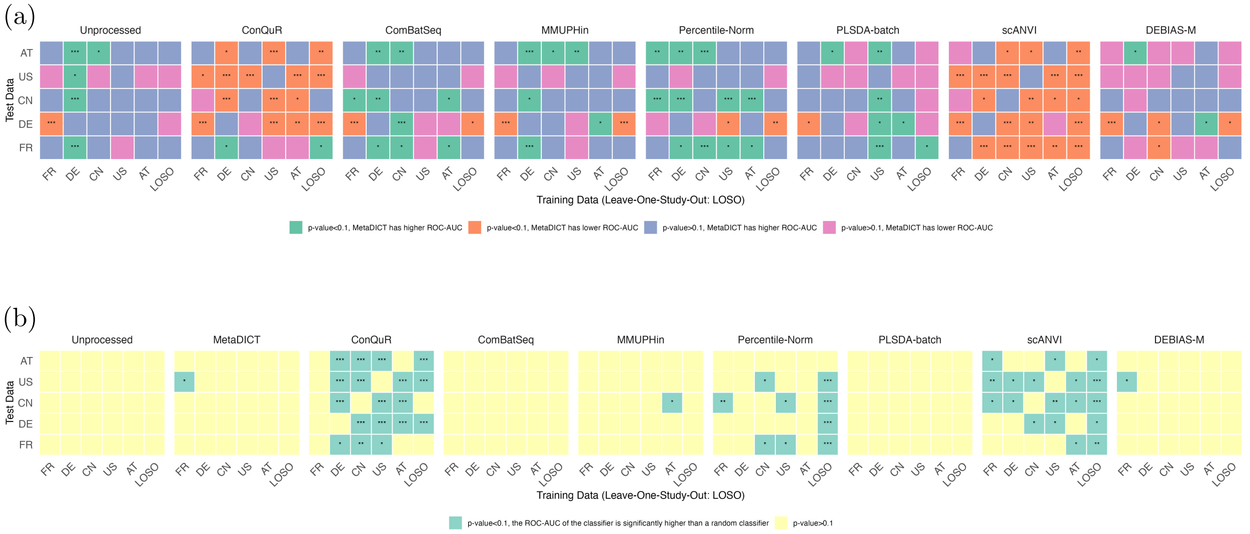


Figure S16: **DeLong's test to compare the ROC-AUCs of classifiers in the CRC integrative analysis.** Figure (a) presents the results of DeLong's test comparing the ROC-AUCs for predicting CRC status between the classifier trained on MetaDICT-integrated data and classifiers trained with other methods. The corresponding ROC-AUC values can be found in Figure 6(d). Figure (b) presents the results of one-sided DeLong's tests comparing the ROC-AUCs for predicting the negative control variable between a random guess classifier and the other classifiers. A $p$-value smaller than 0.1 indicates that the classifier trained on integrated data achieves an ROC-AUC significantly higher than 0.5. The corresponding AUC values can be found in Figure 6(e). AUC differences with $p$-values less than 0.1 are marked with *; those with $p$-values less than 0.01 are marked with **; and those with $p$-values less than 0.001 are marked with ***.

(a)

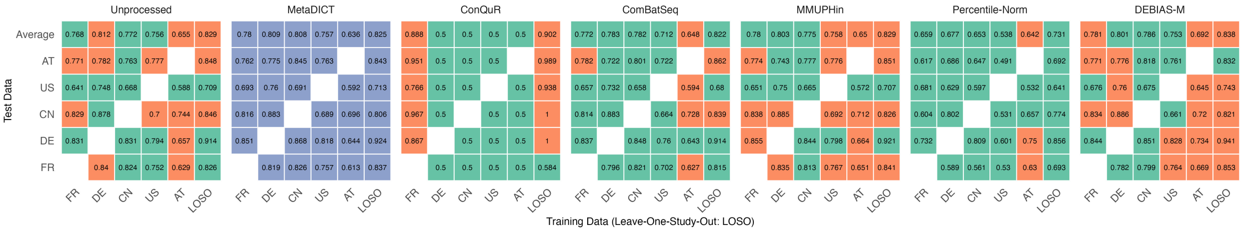


(b)

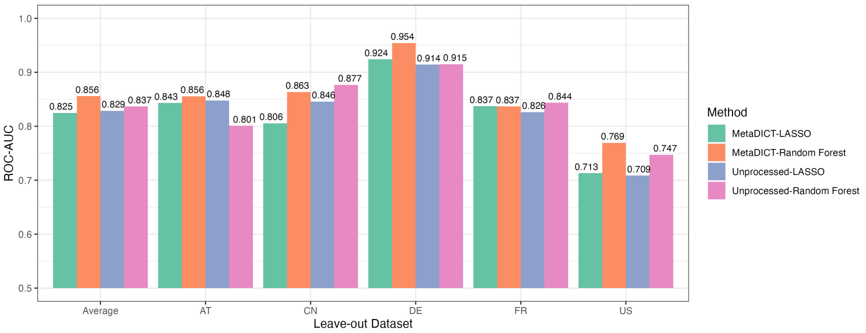


Figure S17: **ROC-AUCs of Lasso logistic regression and random forest classifier trained on integrated data in CRC integrative analysis.** Figure (a) presents the ROC-AUCs of the Lasso logistic regression trained on integrated data at the species level. Green indicates that MetaDICT achieved a higher ROC-AUC, while orange indicates that other integrated data yielded a higher ROC-AUC. Note that PLSDA-batch and scANVI are excluded from this analysis because they are incompatible with log-transformation-based standardization. Figure (b) compares the ROC-AUCs from leave-one-study-out experiments (LOSO) using both the random forest and Lasso logistic regression, applied to MetaDICT-corrected and unprocessed species-level data. The random forest model trained with MetaDICT-processed data achieves highest ROC-AUCs in LOSO experiments. These results suggest that combining an appropriate classifier with an integration method such as MetaDICT can improve prediction performance in real-world data analyses.

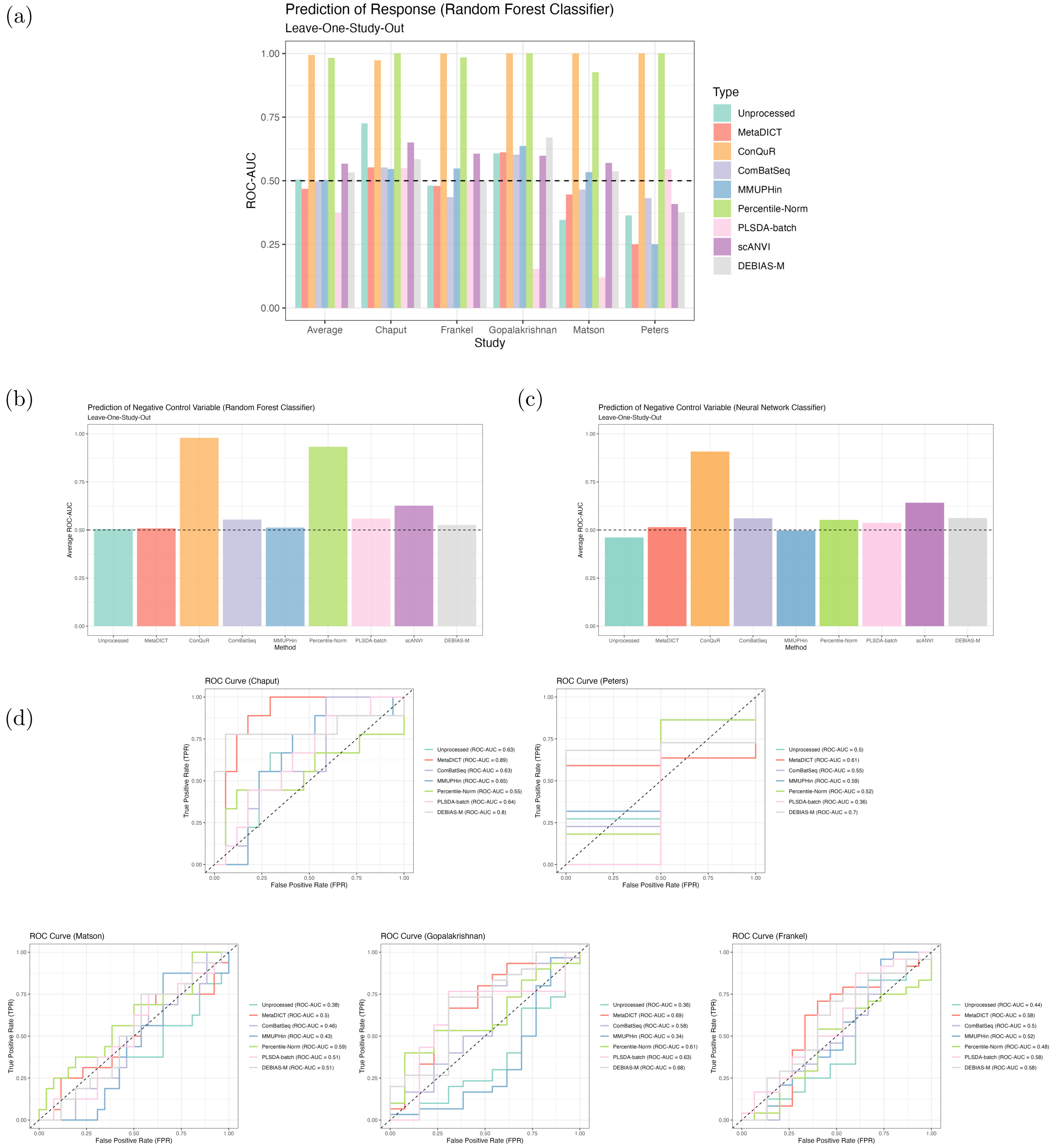


Figure S18: **ROC-AUCs of predictive model for PD-1 immunotherapy response.** Figure (a) presents the ROC-AUC values from leave-one-study-out experiments using a random forest classifier for predicting PD-1 response. Figure (b) shows the average ROC-AUC from leave-one-study-out experiments using a random forest classifier, where the outcome is a negative control—i.e., completely independent of microbial composition but still used in data integration. The random forest model cannot fit these data sets very well and only achieves mediocre performance for most methods without inflating the association in negative control experiments. Figure (d) displays the ROC-AUC curves from leave-one-study-out experiments using a neural network classifier for predicting PD-1 response, while Figure (c) presents the average ROC-AUC from leave-one-study-out experiments using a neural network classifier for predicting the negative control outcome. The comparison suggests that MetaDICT and DEBIAS-M lead to more robust classifiers than other methods.

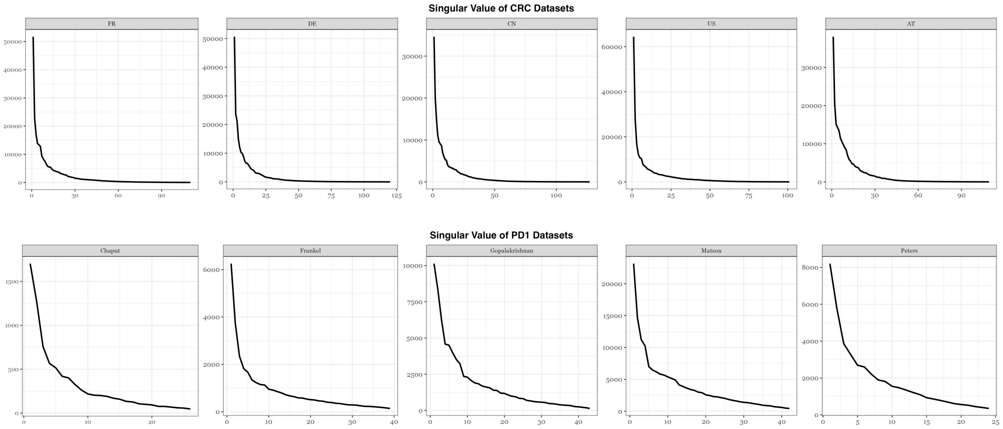


Figure S19: **Singular values of sequencing count matrices for CRC and PD-1 datasets.** A rapid decay in singular values is observed across all datasets in both CRC integrative analysis and PD-1 integrative analysis, supporting the low-rank assumption and suggesting that the data can be effectively approximated using matrix factorization.
